# Discovery of novel covalent ligands with AlphaFold3

**DOI:** 10.1101/2025.03.19.642201

**Authors:** Yoav Shamir, Ronen Gabizon, Adi Rogel, David Yin-wei Lin, Amy H. Andreotti, Nir London

## Abstract

Covalent inhibitors are a prominent modality for research and therapeutic tools. However, a scarcity of computational methods for their discovery slows progress in this field. AI models such as AlphaFold3 (AF3) have shown accuracy in ligand pose prediction, but their applicability for virtual screening campaigns was not assessed. We show that AF3 co-folding predictions and an associated predicted confidence metric ranks true covalent binders with near-optimal classification over property-matched decoys, significantly outperforming state-of-the-art covalent docking tools for a set of protein kinases. In a prospective virtual screening campaign against the model kinase BTK, we discovered a chemically distinct, novel, covalent small molecule that displays potent inhibition in vitro and in cells while maintaining marked kinome and proteomic selectivity. Co-crystallography validated the sub-angstrom accuracy of the predicted AF3 binding mode. These results demonstrate that AF3 can be practically used to discover novel chemical matter for kinases, one of the most prolific families of drug targets.

---

Covalent acting small molecules represent new opportunities in chemical biology and drug discovery (*1–4*). Such ligands typically display increased potency, selectivity and prolonged target engagement compared with their non-covalent counterparts. Advances in chemical biology have mitigated early concerns regarding off-target reactivity and promiscuous irreversible binding and enabled rational and successful design processes (*2*). This has led to the discovery of drugs for decades-long challenging targets such as K-Ras (*5*). Currently more than 50 covalent drugs have been approved by the FDA (*4*). Recent examples include Nirmatrelvir for COVID-19 (*6*), Ritlectinib for alopecia (*7*), and Adagrasib for cancer (*8*), with many others undergoing Phase III clinical trials.

Several computational tools were developed for virtual screening of covalent libraries (*9–16*). Given a protein structure or model as input, these docking tools employ physics-based scoring functions to predict and energetically score the protein-bound pose of each ligand. Multiple covalent binders targeting a range of proteins have been developed and experimentally validated based on the results of such *in silico* screening (*9*, *17–20*). Several covalent datasets of experimental structures have been curated, enabling the evaluation of binding-pose prediction accuracy of docking tools (*21–25*). However, for virtual screening and practical applications, the ability of docking tools to rank a compound library such that active compounds are at the top is arguably more important than accurate pose recapitulation. While datasets designed to evaluate such enrichment are prevalent for non-covalent docking (*26–30*), we are unaware of any for the covalent domain.

AlphaFold3 (AF3), the latest AI-based all-atom structure prediction model from Google DeepMind, was released for free use in November 2024 (*31*). AF3 facilitates high-accuracy prediction of biomolecular complexes, including the prediction of covalent protein-ligand complexes (*32*). To probe the enrichment performance of AF3 in the covalent domain, we constructed COValid, a first-of-its-kind benchmark set for the enrichment analysis of covalent virtual screening. We show that ranking the AF3-predicted compounds using a physics-based scoring function significantly outperforms the ranking produced by classical covalent docking tools. Strikingly, we find an AF3-predicted confidence metric that significantly outperforms all physics-based methods tested across all protein targets in COValid, yielding exceptionally high success rates in identifying active compounds. To mitigate concerns about training data leakage, we performed what is to our knowledge the first prospective covalent virtual screen with AF3, and identified novel, potent and selective BTK covalent inhibitors, that are active in cells.

## Results

### COValid - a novel benchmark for covalent virtual screening

Since cysteines are the nucleophile most commonly targeted by covalent binders, we focused our analysis on the most common electrophile that targets cysteines – acrylamide (Fig. S1) (*33*). We curated compounds with experimental activity annotations from two databases – BindingDB (*34*) and ChEMBL (*35*). The majority of targets we identified were protein kinases, likely due to the intense interest in covalent kinase inhibitors (*33*, *36*, *37*), as they provide superior selectivity and potency for this wide-spread protein family. We included eight of the most populated kinases in COValid (Fig. 1A) as well as K-Ras^G12C^, a prime target for covalent inhibitor design (*5*, *38*). For each target-ligand pair we also annotated the target cysteine position. We made sure that for each target the benchmark included structures with both a covalent and noncovalent binder (selected from the PDB; Dataset S1).

**Figure 1.**
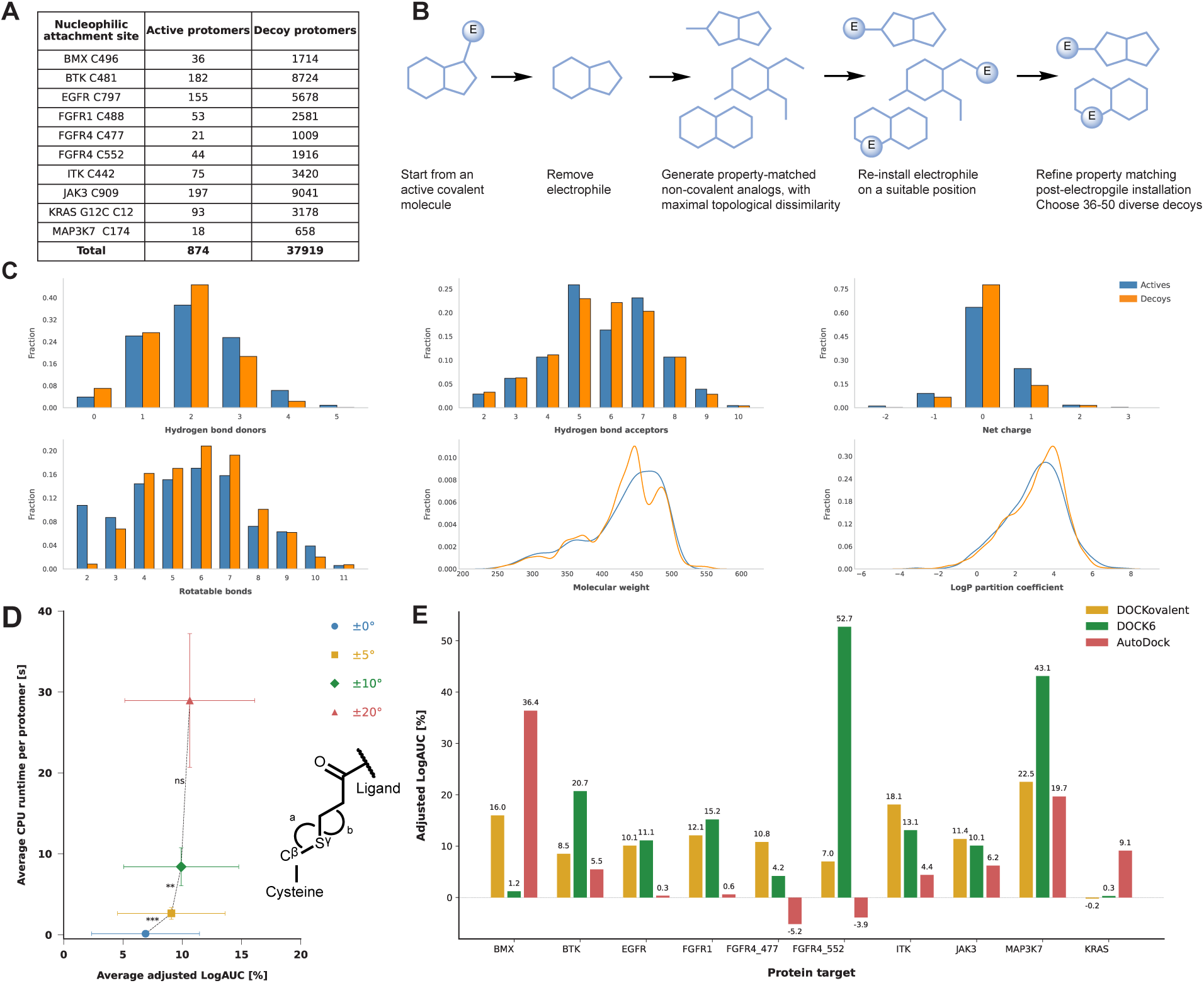
COValid enables optimization of covalent docking algorithms. **A.** Composition of the COValid set. **B.** Decoy design scheme - the electrophile (E) is removed from the covalent ‘active’, next, property-matching, topologically dissimilar non-covalent decoys are collected for all actives. Finally, 36-50 decoy protomers are matched to each active protomer by attaching an acrylamide to free amines of the non-covalent decoys and ensuring property-matching to the active. **C.** Property distributions of the active and decoy protomers (colored blue and orange, respectively) for the six properties matched during decoy design. **D.** Example of using COValid for parameter optimization. The angle range parameter of DOCovalent defines a range Δ around a user-defined angle (109.5°), such that two bonds angles (angles a and b, defined by Cysteine Cβ, Sγ, and the acrylamide Cβ atoms, and by the Cysteine Sγ, acrylamide Cβ and Cα atoms) are sampled ±Δ during docking. Sampling with ±5° and further with ±10° leads to higher averaged adjusted LogAUC with statistical significance (*p-values* are 0.0002 and 0.004, respectively), but further increasing the exhaustiveness to Δ=20° yields no significant improvement (*p-value*: 0.1053), while requiring significantly longer runtime. **E.** Comparison of enrichment for the best identified configuration of three covalent docking tools, docked to the ten covalent PDB structures.

Following the curation of active ligands, we generated 36-50 property-matched, acrylamide-bearing decoys for each ligand (see Supplementary Methods), to represent chemically feasible but likely inactive compounds. This computational design of decoys is necessary since annotations of ‘true’ inactive compounds are scarce. The decoy generation protocol (Fig. 1B) followed two design principles borrowed from the popular DUD-E benchmark set for non-covalent enrichment analysis (*27*). First - the physicochemical properties of the designed decoys should match those of the active. This mitigates the risk that different property distributions for the actives and decoys would yield artificially high enrichment owing to property mismatch (Fig. 1C, Dataset S2) (*39*). The second principle - the chemical topology of the decoys should be dissimilar to that of the actives. We therefore scrambled the connectivity of the compounds to generate decoys that are compositionally matched to actives but unlikely to conserve favorable interactions with the target. Finally, the decoy scaffolds are selected from ZINC20 (*40*) representing commercially available compounds to circumvent the risk of generating non-realistic molecules. In total, COValid includes 874 active protomers and 37,919 decoy protomers, across ten cysteine attachment sites from nine protein targets (Dataset S3).

### COValid enables comparison and optimization of physics-based covalent docking tools

We evaluated three physics-based covalent docking tools using COValid: the DOCKovalent method (*9*) based on DOCK3.7, the attach-and-grow method based on DOCK6.12 (*15*), and the flexible sidechain method of AutoDock (*10*). To quantify the enrichment performance, we used the adjusted LogAUC metric (*41*), which emphasizes early enrichment in ranking hit lists (*42*). In a real-life scenario it is more important to have active molecules at the top fraction of the hit list than overall better ranking across the entire library. An adjusted LogAUC of 0% equals random performance, whereas 85.5% is optimal performance, ranking all actives higher than all inactives (*43*).

Each docking tool offers many adjustable parameters and configurations that can be non-trivial to optimize and can result in significant increases in run-time. Since evaluating all different combinations of parameters is an intractable combinatorial problem, we chose several representative features for each software and assessed their effect on accuracy and runtime (See Fig. 1D for one example, and the Supplementary Results for the full analysis).

The performance of leading configurations of DOCK6 and DOCKovalent were similar (Fig. 1E; Table 1; see docking parameters in Table S1). AutoDock, with a similar run-time showed an average enrichment slightly lower than that of DOCKovalent and DOCK6 (*p-values* of 0.05 and 0.06, respectively). For the three docking tools, the differences in the enrichment across the covalent-bound structures, or the non-covalent structures, were insignificant (*p-values > 0.1*), indicating that non-covalent *holo* structures could be useful for covalent virtual screening when a covalent complex structure is not available.

**Table 1.**
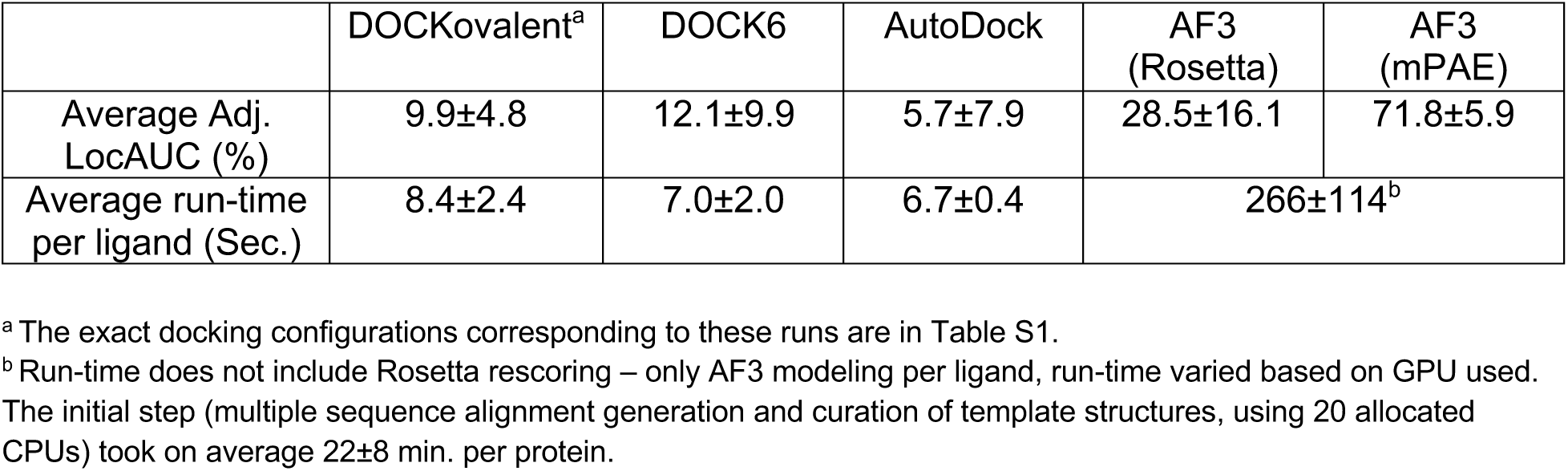
Performance over the COValid set.

### Physics-based scoring of AlphaFold3 models outperforms docking tools for multiple targets

AF3 is the latest AI model from DeepMind for all-atom prediction of biomolecular structures (*31*). It enables prediction of non-covalent as well as covalent protein-ligand complexes. On a pose-prediction benchmark set of covalent complexes curated by the AF3 authors, the model achieved a success rate (defined as RMSD<2Å across the pocket atoms) of 78.5% (*31*). To our knowledge AF3 performance in the context of covalent virtual screening (or in fact for virtual screening in general) has not been reported. We set out to use COValid to assess its capacity for screening. We used the prediction pipeline of AF3 to predict covalent complexes for all compounds in COValid using the protein sequence and geometrically optimized 3D ligand conformer as input. The input also includes specification of the ligand atom and the protein side-chain atom to be covalently bonded.

We sought to rank the predicted complex structures using a physics-based docking score. We used Rosetta (*44*) to minimize and energetically score all the predicted AF3 models and used this score to calculate the enrichment performance across COValid. Re-scoring of AF3 models yielded significantly better results for five out of the 10 sites (Fig. S2) and an overall better average adjusted LogAUC across the ten sites (Table 1). The fact that this enrichment was achieved just based on the models produced by AF3 (and not by any AI-based metric) suggests the models are sufficiently realistic to differentiate true binding modes from artificial ones. Indeed, in the few cases where a crystal structure was available for a target-ligand pair from the benchmark, the AF3 models proved accurate (SI Fig. S3). While most of these examples were likely included in the AF3 training set, one structure was determined after the training, and in this case as well AF3 predicted it accurately (0.45Å pocket-aligned RMSD; PDB:7O70, Fig. 2A).

**Figure 2.**
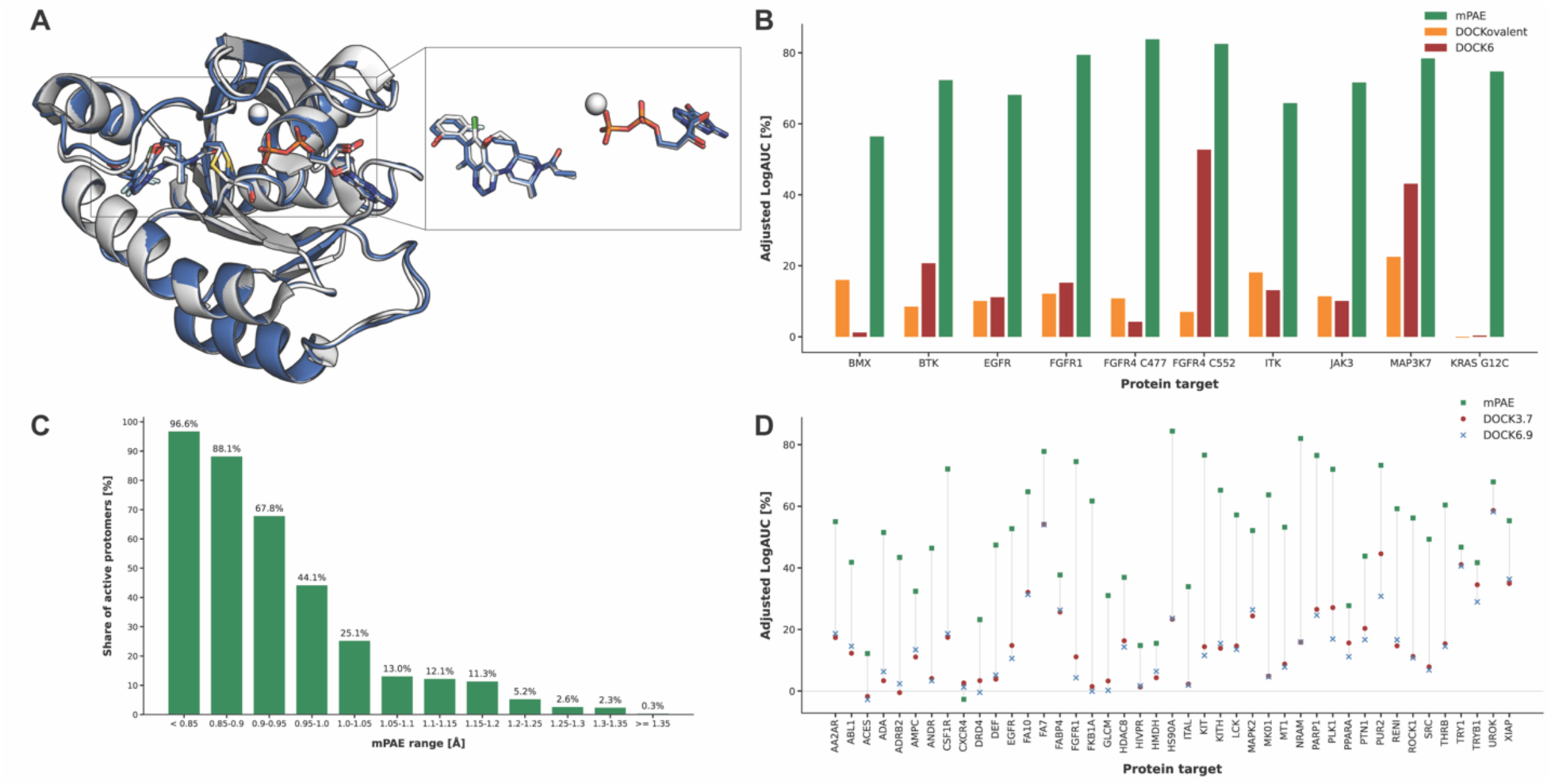
Near-optimal enrichment of true inhibitors using AF3 mPAE. **A.** Example of an accurate AF3 prediction of a covalent complex (KRAS G12C with Mg^2+^ and GDP, PDB 7O70, pocket-aligned RMSD 0.45Å) not included in the training set (the experimental structure and the predicted model are colored in gray and blue, respectively). **B.** Comparison of the two best physics-based covalent docking enrichment to AF3 structural predictions ranked by mPAE. **C.** The distribution of the proportion of active COValid protomers, according to their AF3 mPAE element. Each column indicates the share of active protomers out of all protomers within the indicated mPAE value range. This could be interpreted as the probability for being ‘active’ given a mPAE value. **D.** AF3 mPAE enrichment performance on DUDE-Z targets. Across all 43 DUDE-Z targets, we compare the adjusted LogAUC values kindly provided by Balius et al. for DOCK3.7 and DOCK6.9 (*45*) to values derived from our AF3 predictions, ranked by mPAE (actives are sorted prior to enrichment calculations, using only the best-scoring active protomer of each compound). Protein sequence input for the data step of AF3 was extracted from the DUDE-Z PDB files, and the non-covalent ligands were provided as SMILES strings from DUDE-Z.

### AF3 predicted alignment error is a superior metric for virtual screening enrichment

Along with the 3D coordinates of the predicted molecular structure, the AF3 model outputs predicted confidence metrics for the accuracy of the pose recapitulation. We were interested in the ability of such metrics to rank compound libraries and enrich actives, even though these metrics were not trained to predict activity or affinity. We evaluated the adjusted LogAUC values over the COValid set for various AF3-predicted confidence metrics (see definitions in Supplementary Methods and results in Fig. S4). When attempting to rank COValid AF3 models by these metrics, we noticed that most metrics lack the resolution required for a decisive ranking of the lists. In several cases, thousands of protomers were ranked with the same exact predicted value (Fig. S5). Nevertheless, to probe their enrichment performance, we evaluated the adjusted LogAUC value for each cysteine site as a range spanning two extremes: we assessed a best-case scenario, where for any given value we rank actives higher than decoys for the same metric value, and a worst-case scenario, where decoys are ranked higher than actives of the same value.

Ranking by a global metric (pTM) or protein-based metric (protein chain pTM) yielded large ranges between the best- and worst-case enrichment values (Fig. S4). This lack of resolution is perhaps in line with the fact that the protein is constant for any specific target. Taking into account the protein-ligand interface (ipTM and ‘ranking score’), yielded smaller, more informative enrichment ranges for some targets, while other targets still displayed a wide range of enrichment values. The metric that yielded the most decisive adjusted LogAUC ranges of this group was the ligand chain pTM. Even when using the worst-case ranking, AF3 predictions sorted by the ligand chain pTM metrics significantly outperformed the physics-based docking tools across nine of the ten COValid Cys sites (Fig. S6).

Finally, we studied the relevance of the predicted aligned error matrix (PAE), a structural confidence metric provided by AF3. In the internal representation used by the AF3 model, each protein residue and each ligand atom are represented by a single token. Each (*i,j*) element of the PAE matrix contains a prediction of the error in the positions of the AF3 token *j* when aligned to the ground-truth based on token *i* (*31*). We define mPAE as the minimal predicted error in the positions of the protein residues, when aligned to the ground-truth complex with respect to the ligand atoms (See details in Supplementary Methods; this value is provided as output by AF3).

Compared with other AF3-predicted confidence metrics, mPAE yields much better resolution between compounds, enabling conclusive ranking of all ten COValid compound lists. Remarkably, AF3 modeling followed by ranking by mPAE, significantly outperforms other metrics including all covalent docking tools across all ten cases (Fig. 2B; Fig. S4). The worst-case adjusted LogAUC values ranged from 56.4% for BMX to 83.8% for Cys477 of FGFR4, approaching optimal classification. Comparing the enrichment performance to that of the physics-based covalent docking tools, the differences are striking (Fig. 2B). If we disregard early enrichment and report the more typical ‘AUC’ (area under the receiver operating curve), the values range from 0.9082 to 0.9996 (Table S2).

We also studied the relevance of the absolute mPAE values for prospective covalent screening. Within the applicability domain of COValid, mPAE enables assignment of confidence to the classification of a compound as active (Fig. 2C). Across all COVAlid protomers, 96.6% of the protomers with the lowest mPAE values (<0.85Å) correspond to actives. As mPAE values increase (indicating a decrease in structural confidence) the probability that the protomer corresponds to an active binder drops monotonically, decreasing to less than 13% at mPAE values higher than 1.05Å.

We considered various confounding factors, such as the possibility of overlap between the AF3 training set and the COValid compounds. If the model was trained on some COValid ‘actives’, it could potentially predict higher structural confidence for these compared to the designed decoys. We conducted ablation studies by removing active compounds with high topological similarity to any compound in the PDB (along with their corresponding decoys; Fig. S7) and re-calculated the adjusted LogAUC values in the absence of the removed protomers (Fig. S8). Even at strict similarity cutoffs (Tanimoto coefficient=0.4, Morgan fingerprint, radius=2, bit=2048) that remove most of the active and decoy protomers from the ranked lists, enrichment remains high, with little effect due to ablation. This suggests that the performance of mPAE is not due to overlap with the training data. In addition, analysis of the mPAE element of all the COValid ‘actives’ against their maximal Tanimoto similarity to any PDB yielded no correlation between the two metrics (Pearson correlation coefficient −0.056, *p-value* 0.21; Fig. S9).

Another potential confounding factor is that our decoys, despite matching in physicochemical properties and being based on commercially available scaffolds, differ in some rudimentary way from the ‘actives’. To address this, we searched ChEMBL 33 (*35*) for experimental decoys - acrylamide-bearing structures that were tested against benchmark targets but showed worse than 10 μM activity (Fig. S10). We found 64 experimental decoys across six COValid kinases. We used AF3 to predict their covalent complexes and evaluated their mPAE values. The average mPAE was 1.5±0.5Å, with 86% of the decoys yielding a mPAE higher than 0.95Å.

Recent work (*46*) suggested that co-folding models do not learn the ‘physics’ of ligand binding, and demonstrated their insensitivity to local mutations in the binding site. We found a similar trend. First, there is no apparent correlation between mPAE and affinity (Fig. S11) suggesting it is a ‘coarse’ metric useful primarily for classification. Looking further into selectivity, we conducted cross-docking experiments and calculated the enrichment for each kinase active and decoy set against all non-cognate kinases. (Fig. S12). While generally the highest enrichment is still achieved against the cognate kinase, clear clusters appear, in which kinases with a similarly positioned cysteine residue perform well in enriching non-cognate ligand sets. To some extent this could be a result of non-selectivity of the active ligands, which are known to bind kinase off-targets with analogous cysteines (*47–49*). But overall it likely suggests that AF3 and mPAE are currently not suitable for selectivity prediction.

### mPAE performs well also for non-covalent virtual screening

We tested the relevance of AF3 predictions with mPAE ranking for the non-covalent case using the DUDE-Z benchmark set (*28*). Across 42 out of 43 targets, AF3-based enrichment significantly outperformed the results reported for either DOCK3.7 or DOCK6.9 (Fig. 2D; the exception is CXCR4, which yields near-random enrichment across all three screening methods). Across this diverse set, we observe a larger variability in enrichment values than in COValid, with an average enrichment of 50.9%±20.0%, suggesting some targets are more challenging than others. Indeed, kinases exhibited higher mPAE-based enrichment on average, when compared to other targets (11 kinases yielded an average adjusted LogAUC of 60.7%±11.1%, compared to 47.5%±21.3% on average for the other 32 protein targets). Across this more diverse set we do see a moderate correlation between performance and the average maximal chemical similarity of the active compounds to PDB ligands (Pearson correlation coefficient=0.425, *p-value* = 0.00451, R^2^=0.18; Fig. S13). Boltz-2, another AI based co-folding model (*50*) showed similar performance to AF3 on this set (see Supplementary Results; Fig. S14).

### Covalent prospective screening identifies novel kinase inhibitors

To evaluate AF3-based covalent screening and probe its relevance for the discovery of novel covalent binders, we conducted a prospective campaign against Bruton’s tyrosine kinase (BTK), a well-studied kinase target, whose dysregulation is implicated in B-cell malignancies (*51*). BTK has multiple FDA-approved covalent inhibitors targeting Cys481 at the ATP binding pocket (*52–55*). We constructed a diverse virtual library of ∼906K acrylamide-bearing compounds, and used AF3 to predict their covalent complexes with Cys481 of BTK (see Supplementary Methods; Dataset S4). We ranked the predictions by their mPAE scores, and filtered out compounds with mPAE greater than 0.9Å (resulting in 440 compounds; ∼0.05% of the library). In search of novel binders, we filtered out 50 compounds showing even remote chemical similarity to known BTK binders (Tc > 0.35). The remaining 390 compounds were clustered and manually inspected in search for compounds with diverse binding poses. Of these, we synthesized 13 compounds for experimental evaluation (Fig. S15).

Intact protein LC/MS experiments indicated that three of the 13 compounds reached near-100% covalent labeling of BTK within 2 hours (Fig. 3A,B; Fig. S16, S17). We further characterized the three main hits - YS1, YS2, and YS3, and used ibrutinib, an FDA-approved drug for BTK, as a control (*52*) (Fig. 3C). We note that these compounds bear no chemical resemblance to any ligands across all of ChEMBL (Fig. S18). Dose response and time course experiments indicated that YS1 covalently labels BTK rapidly, even in low concentrations, approaching the binding kinetics of ibrutinib (Fig. 3D,E; in a 2h dose response assay, ibrutinib and YS1 yield >90% labeling at 1𝛍M). To mitigate concerns that labeling arises from intrinsic reactivity of the molecules rather than target recognition, we conducted GSH reactivity assays, which indicated that binding is not driven by intrinsic electrophilicity - all three compounds are less reactive than ibrutinib (YS1 and YS2 are significantly less reactive, Fig. 3F). Differential scanning fluorimetry (DSF) experiments showed that all three compounds stabilize BTK, with YS1 leading to the largest shift in melting point (∼9℃; Fig. 3G). In vitro kinase activity assays identified YS1 as a potent inhibitor of BTK, (IC_50_=30 nM; Fig. 3H; IC_50_ values for YS2 and YS3 are 77 𝛍M and 8.4 𝛍M, respectively). YS1 showed no activity with the C481S mutant of BTK (Fig. S19), indicating its binding is driven primarily by the covalent interaction.

**Figure 3.**
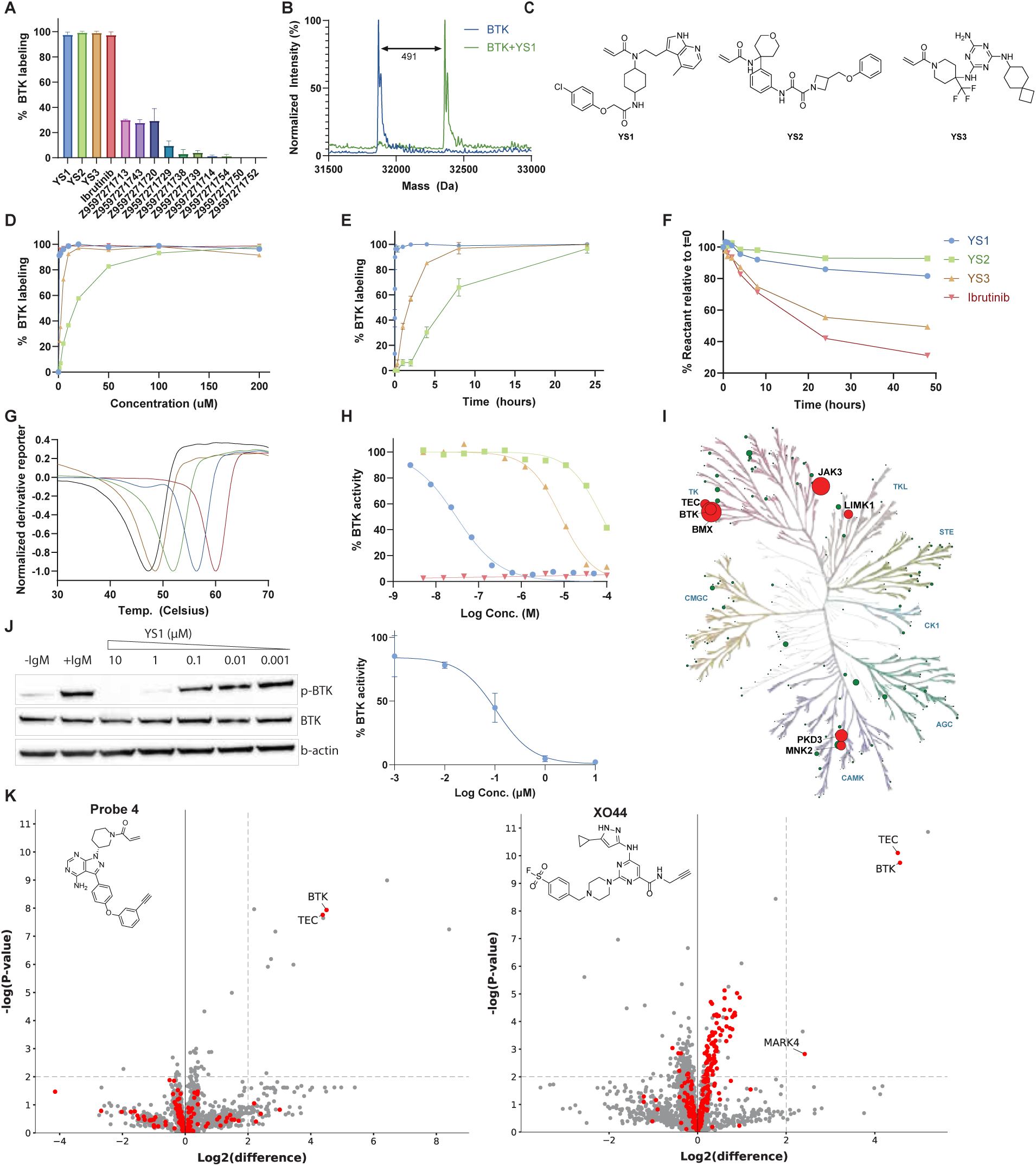
Experimental characterization of novel BTK inhibitors discovered by prospective screening. **A.** Intact protein LC/MS covalent labeling percentage for 13 synthesized compounds (1 μM BTK, 200 μM compound, pH 7.5, RT, 2h). n = 2, data represented as mean with SD as error bars. **B.** Deconvoluted LC/MS spectrum of BTK incubated with YS1. The mass difference corresponds to the ligand adduct, validating covalent binding. **C.** Three main binding hits. **D.** LC/MS dose response labeling experiment (1 μM BTK, pH 7.5, RT, 2h) for main hits and ibrutinib. **E.** LC/MS time-course experiment (1 μM BTK, 5 μM compound, RT, pH 7.5). n = 2, data represented as mean with SD as error bars. **F.** Reduced glutathione (GSH) assay for reactivity assessment (5 mM GSH, 0.2 mM compound, pH 8, 25℃). **G.** Differential scanning fluorimetry (DSF) analyzes the shift in BTK melting point after treatment with the compounds (5 μM BTK kinase domain, 0.02 mM compound, pH 7.5, overnight incubation at 25°C). The derivative reported, normalized by the absolute value of the minimum in each dataset, was used to determine the melting point. Ibrutinib, YS1, YS2 and YS3 stabilize BTK by ∼13°C, 9°C, 5°C, and 1°C, respectively. BTK baseline curve is plotted in black. **H.** Kinase inhibition analysis by a radiometric filter binding assay measuring phosphorylated substrates products (20 μM ATP, 3 nM BTK, 0.2 mg/ml pEY substrate, pH 7.5, RT). IC_50_ of YS1, YS2, and YS3 is 30 nM, 77 𝛍M and 8.4 𝛍M, respectively. **I.** Kinome phylogenetic tree depicting inhibition data from an assay conducted with YS1 against 362 kinases. Each kinase tested is represented by a dot; dot size is relative to the mean percentage of inhibition with respect to DMSO across two replicates at a dose of 300 nM YS1. Dots are colored red if mean inhibition is greater than 40%, and green otherwise. Illustration reproduced courtesy of Cell Signaling Technology, Inc. (www.cellsignal.com). **J.** Dose-dependent BTK activity assay in Mino cells as measured by autophosphorylation of BTK. The cells were incubated for 1h with either DMSO or various concentrations of YS1. The cells were activated with anti-IgM, and BTK autophosphorylation was quantified by Western Blot and normalized with respect to β-actin. IC_50_ (107 nM) was calculated by fitting the data to a dose−response curve using Prism software (n = 3). **K.** YS1 selectivity quantification via competitive pull-down experiments with Probe 4, an alkynylated probe analog of ibrutinib (left) and XO44, a generic kinase probe (right), respectively. Mino cells were treated with either DMSO or 1 μM of YS1 for 1h, followed by 45 min treatment with either 1 𝛍M Probe 4 or 2 𝛍M XO44 (n = 4). Proteins were quantified using label-free quantification. Proteins in the upper right segment show a significant change (fold change > 2; p-value < 0.01). In the Probe 4 experiment, 3,416 proteins are plotted (159 kinases and 3,257 non-kinases), out of which 11 exhibit significant competition (2 kinases and 9 non-kinases). In the XO44 experiment, 3,569 proteins are plotted (235 kinases and 3,334 non-kinases), out of which 5 exhibit significant competition (3 kinases and 2 non-kinases). Kinases were determined by a list of human kinase Uniprot entries from KinHub (*59*).

We conducted a kinome-wide inhibition assay for YS1 against 362 recombinantly expressed kinases (Fig. 3I; Dataset S5). Overall, YS1 exhibits marked selectivity across kinases, with only six off-target kinases inhibited by 40% or more. Three of the off-targets: JAK3, BMX, and TEC (85.1%, 95.3%, and 45.6% mean inhibition, respectively) are members of the TK kinase group, along with BTK, and all contain a Cys residue at an analogous position, making them susceptible to off-target covalent binding. BMX in particular is a challenging off-target since ten clinical BTK inhibitors are known to inhibit it *in vitro* (*56*). Of note, other kinases with an equivalent cysteine: ITK, BLK, TXK, HER2, HER4, EGFR and MKK7, were not significantly inhibited (Dataset S5). The remaining off-targets were LIMK1, PKD3 and MNK2.

To measure the efficacy of the compounds in cells, we conducted dose-dependent cellular BTK activity assays measuring the autophosphorylation of BTK. Mino cells were pre-incubated with YS1, followed by BTK activation with an anti-IgM antibody and activity was measured by western blotting. YS1 potently inhibits BTK activity in cells (Fig. 3J, Fig. S20, IC_50_=107 nM). To evaluate proteomic selectivity, we conducted two cellular proteomic competition assays of YS1 against “Probe 4”, an alkynylated probe analog of ibrutinib, and XO44, a generic kinase probe (*57*, *58*) (Fig. 3K; Fig. S21; Dataset S6). In both assays, YS1 is shown to be extremely selective for BTK in cells. In a competition with Probe 4, only a single kinase off-target - TEC - is significantly competed (log fold-change > 2; *p-value* < 0.01). In a competition experiment with XO44, two kinase off-targets are significantly competed - TEC and to a much lesser extent MARK4. We should note that despite XO44 pulling down LIMK1, PKD3 and MNK2, none were competed in cells by YS1 (Dataset S6).

Finally, we crystallized and acquired x-ray diffraction data for YS1, YS2 and YS3 in complex with the BTK kinase domain. The YS1 and YS2 complex structures were determined to resolutions of 1.6Å and 1.27Å (PDB: 9ZLJ, 9ZLM respectively; Fig. 4A,B; see Table S4 and Supplementary Methods). Unambiguous electron density is observed for the entirety of the ligands and the covalent attachment to Cys481 (Fig. S22). The structures closely match the AF3 predictions (Fig. 4A, ligand heavy atom RMSD of 0.50Å and 0.41Å, for YS1 and YS2 respectively, when aligned by the protein backbone). The YS1/BTK structure validates YS1 as the trans-isomer of the cyclohexyl moiety, as predicted by AF3; we were able to isolate a second isomer, presumably the cis-isomer, that produced similar, but slightly less potent results (Fig. S23).

**Figure 4.**
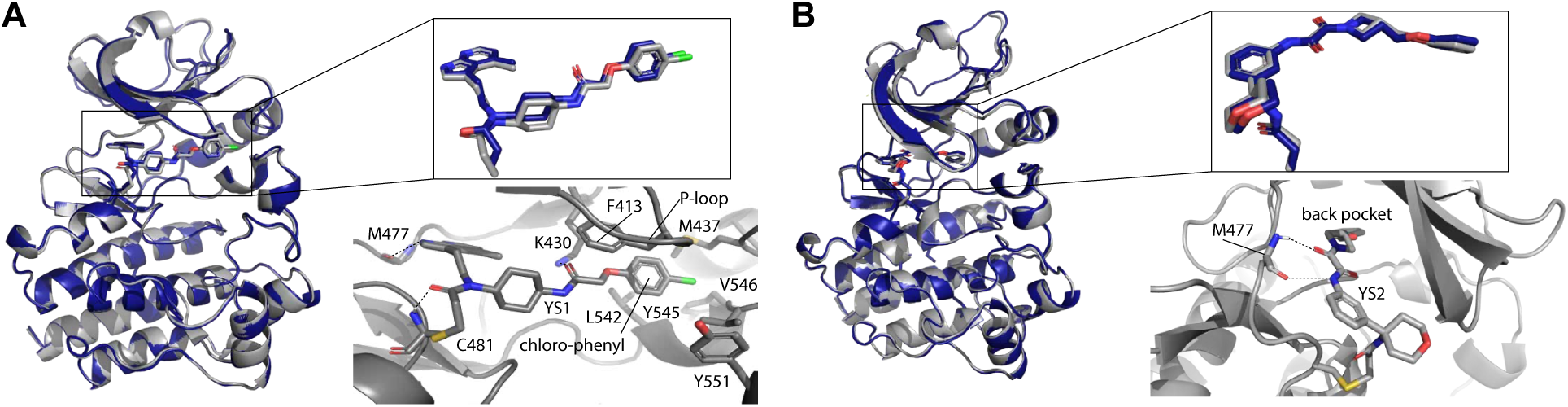
AF3 structural predictions are accurate. Overlay of AF3 predictions (blue) and X-ray crystallography experimental results (gray) for YS1 (A) and YS2 (B) covalently attached to Cys481 of WT-BTK kinase domain. Overlay of the predicted ligand pose and the experimentally determined ligand structure are also shown separately for clarity. Heavy atom ligand RMSD values are 0.50Å and 0.41Å, respectively. Close up view of each ligand in the BTK active site are shown with interacting residues labeled. Putative hydrogen bonds are depicted as dashed lines.

Both YS1 and YS2 form hydrogen bonds with the backbone of Met477; in YS1 the azaindole group forms two hydrogen bonds with the backbone of Met477 while in YS2 the atypical amide containing linker between the azetidine and benzyl groups forms the two hydrogen bonds to Met477 (Fig. 4A,B). Additional hydrogen bonds in the YS1 complex include the carbonyl of the acrylamide warhead with the backbone amide of Cys481 and a hydrogen bond between the Lys430 sidechain and the other amide carbonyl in the ligand (Fig. 4A).

Similar to many well characterized BTK inhibitors, YS2 extends into the ‘back pocket’ of the BTK active site (Fig. 4B). YS1 is not a ‘back pocket’ binder nor can it be characterized as a front pocket binder. YS1 protrudes under the kinase P-loop (or glycine rich loop) and extends its chloro-phenyl group into a subpocket formed by the side chains of M437 (next to the C-helix in the N-lobe) and L542/Y545/V546, which are located at the N-terminal end of the activation loop (Fig. 4A). This subpocket sits above the pocket normally occupied by inhibitors that bind to the front pocket. The YS1 bound structure shows that the P-loop is repositioned into an open conformation to accommodate the ligand and the sidechain of F413 adopts an unusual orientation where it points toward the ATP binding cleft rather than toward the activation loop (Fig. 4A). In fact, structures of front pocket ligands bound to BTK show that the sidechain of F413 typically occupies the pocket filled by the chloro-phenyl group of YS1 (Fig. S24). In addition, front pocket binders can ‘sequester’ the activation loop tyrosine, Y551 (*60*); but the conformation of Y551 in the YS1 complex structure does not conform to the sequestered state. Thus, AF3 predicted a unique BTK inhibitor that expands our understanding of the druggable pockets within the BTK active site.

The x-ray data for the YS3/BTK kinase domain complex only diffracted to ∼3.5Å. This resolution does not allow for detailed analysis of ligand binding as described above for YS1 and YS2, but is nevertheless sufficient to determine general ligand binding orientation in the crystal. Interestingly, the binding mode differs from the AF3 predicted binding pose of YS3 (Fig. S25). The AF3 prediction places the spiro(3.5)nonane portion of the YS3 ligand toward solvent while the x-ray diffraction data show clear electron density extending toward the back pocket of the kinase active site. The predicted pose for YS3 and the pose suggested by the ligand electron density are related by rotation around one bond. It is worth noting that the predicted pose for YS3 is quite similar to the structure of CC-292 bound to the BTK kinase domain (*60*) (Fig. S26). Solution studies of CC-292 bound to the BTK active site reveal multiple conformations and binding of CC-292 does not stabilize the BTK kinase domain (*61*). The observation that YS3 stabilizes BTK by only 1°C (Fig. 3G) and the fact that the AF3 prediction differs from the x-ray diffraction data for this ligand may be consistent with an inhibitor that adopts more than one conformation when covalently attached to BTK C481. Nevertheless, YS3 effectively inhibits BTK in a range of biochemical assays (Fig. 3).

## Discussion

Virtual screening of large ligand libraries is a cornerstone of modern drug discovery (*62*). Despite the growing prominence of covalent drug discovery, prospective covalent virtual screening remains underdeveloped. We sought to test whether AF3 can address this gap. Inspired by the importance of benchmarks of noncovalent molecules for the development of virtual screening in that domain (*27*) we developed COValid. Our key finding was that AF3 vastly outperformed traditional methods. This is especially encouraging as it only requires the sequence of the protein and the target ligand rather than a crystallographic structure of the protein as do conventional virtual screening approaches.

Previous studies have shown that AI models trained on ligand binding data could not surpass AutoDock Vina on the DUD-E enrichment test set (*63*). Moreover, an analysis conducted prior to the development of AF3 revealed that deep learning–based methods lagged behind classical docking tools, particularly in generating physically valid poses (*64*). Most such analyses focused, however, on pose reproduction (*65*). In terms of screening, AI models were only evaluated for their capacity to generate structural protein models to support virtual screens using traditional docking tools (*66–68*). In contrast, we show for the first time that AF3 outperformed classical docking in both covalent and non-covalent screening scenarios (Fig. 2).

Several factors may contribute to the exceptional performance. First, specifying the covalent attachment point may improve accuracy; prior work demonstrated that indicating the ligand binding pocket increased pose recapitulation accuracy from 81% to 93% in AF3 (*31*). Second, by incorporating backbone flexibility, AF3 can resolve minor clashes that typically confound classical fixed-backbone docking methods or expose cryptic pockets. This point is exemplified by our prospective screen. When we used DOCKovalent to screen the same library against a static structure of BTK (covalent complex with Ibrutinib; PDB: 5P9J), YS1 was ranked only at the 15th percentile of the list (as opposed to the top 0.05% via mPAE-ranked AF3) and its pose did not resemble the crystal structure, likely due to a clash with the side-chain of F413 (Fig. S27).

A more cautious explanation, however, is that the structural data used to train AF3 might overlap with our test set. Kinases—common drug targets that share similar features—are heavily represented in both the PDB and in drug discovery generally (*63*, *64*). Our dataset is heavily biased toward kinases (nine out of ten targets), highlighting the potential for data leakage. Yet, this bias also underscores the importance of kinases in the field, given that only this class of proteins provided a critical mass of annotated covalent binders. It is possible, and even likely, that the impressive performance we report is limited to kinases; even so, the impact on accelerating ligand discovery for this crucial class would still be immense. Encouragingly, two observations suggest broader generalization. First, AF3 achieved high enrichment against KRas—a GTPase unrelated to kinases—even when considering only actives dissimilar to any PDB ligand (Fig. S8). Second, AF3 demonstrated strong non-covalent enrichment across dozens of non-kinase targets, albeit not as high, on average, as the non-covalent enrichment for kinases (Fig. 2D).

AF3 confidence metrics were trained to quantify modeling accuracy. Most predicted metrics lack the resolution to rank large libraries of models (Fig. S4), particularly those focusing on the protein side. A recent adversarial experiment suggested that AF3 does not rely on specific physical interactions for ligand placement (*46*), and indeed, there is no apparent correlation between the mPAE score and the binding affinity of experimental actives (Fig. S11). Nevertheless, the fact that rescoring AF3 models with Rosetta—an established physics-based scoring method—significantly outperformed covalent docking tools in terms of enrichment for five of the ten sites suggests that the models generated by AF3 are physically sensible. In that respect, while AF3’s covalent docking tends to produce malformed poses with respect to the bond length and angle geometry (Fig. S28), it compensates so that the overall ligand pose remains accurate. An advantage of mPAE is its apparent insensitivity to the protein target, allowing comparisons across complexes and enabling the assignment of binding probabilities based on mPAE values (Fig. 2C). Cross-docking experiments (Fig. S12) demonstrated that mPAE can discriminate when covalent docking to different cysteine locations, but not so well within the same location, suggesting it should not be used currently for selectivity predictions.

In practical terms, despite AF3’s superior enrichment, its long run-time may render it impractical for screening the extremely large virtual libraries that have become standard (*69–71*). A feasible solution is to use a fast classical docking method to screen very large virtual libraries—potentially comprising billions of compounds—followed by re-docking the top million or so predicted compounds with AF3 to more effectively enrich true binders.

To mitigate the aforementioned confounding factors, the ultimate validation of real-world applicability is experimental validation. The fact that we were able to discover novel chemical matter in a prospective manner, clearly indicates that at least for well-studied protein targets with well-defined pockets, such as kinases, AF3-based prospective screening can go beyond any examples it has observed during training. Such screening opens the door for incorporation of protein flexibility in large scale virtual screening for the first time, and may unlock more challenging, flexible targets that are recalcitrant to state-of-the-art docking methods.

## Supporting information

Supp_Dataset_1_experimental_structures

Supp_Dataset_2_properties_per_protein

Supp_Dataset_3_active_and_decoy_protomers

Supp_Dataset_4_prospective_virtual_library

Supp_Dataset_5_YS1_kinome panel

Supp_Dataset_6_YS1_proteomics

## Acknowledgements

We thank Sarel Fleishman for critical reading of the manuscript. The Irwin lab in UCSF for access to their cluster for DOCK6 and DOCKovalent ligand generation and in particular Dr. Khanh Tang for technical assistance. We thank Dr. Trent Balius for assistance with DOCK6.12, as well as sharing DUDE-Z enrichment calculation data, and Dr. Alexey Orlov for sharing his ligand generation pipeline. Y.S. is funded by the CHE fellowship for data sciences. This research was generously supported by the Knell Family Institute of Artificial Intelligence, the Honey and Dr. Barry Sherman Lab, the Dr. Barry Sherman Institute for Medicinal Chemistry, the Abisch-Frenkel RNA Therapeutics Center, the Moross Integrated Cancer Center, the Goldhirsh-Yellin Foundation and Celia Zwillenberg-Fridman. D.Y.L. and A.H.A. thank the Roy J. Carver Charitable Trust for financial support.

## References

1. T. A. Baillie, Targeted covalent inhibitors for drug design. Angew. Chem. Int. Ed Engl. 55, 13408–13421 (2016).

2. L. Boike, N. J. Henning, D. K. Nomura, Advances in covalent drug discovery. Nat. Rev. Drug Discov. 21, 881–898 (2022).

3. N. London, Covalent proximity inducers. Chem. Rev. 125, 326–368 (2025).

4. F. Sutanto, M. Konstantinidou, A. Dömling, Covalent inhibitors: a rational approach to drug discovery. RSC Med. Chem. 11, 876–884 (2020).

5. J. M. L. Ostrem, K. M. Shokat, Targeting KRAS G12C with covalent inhibitors. Annu. Rev. Cancer Biol. 6, 49–64 (2022).

6. J. Hammond, H. Leister-Tebbe, A. Gardner, P. Abreu, W. Bao, W. Wisemandle, M. Baniecki, V. M. Hendrick, B. Damle, A. Simón-Campos, R. Pypstra, J. M. Rusnak, EPIC-HR Investigators, Oral nirmatrelvir for high-risk, nonhospitalized adults with Covid-19. N. Engl. J. Med. 386, 1397–1408 (2022).

7. H. A. Blair, Ritlecitinib: First approval. Drugs 83, 1315–1321 (2023).

8. P. A. Jänne, G. J. Riely, S. M. Gadgeel, R. S. Heist, S.-H. I. Ou, J. M. Pacheco, M. L. Johnson, J. K. Sabari, K. Leventakos, E. Yau, L. Bazhenova, M. V. Negrao, N. A. Pennell, J. Zhang, K. Anderes, H. Der-Torossian, T. Kheoh, K. Velastegui, X. Yan, J. G. Christensen, R. C. Chao, A. I. Spira, Adagrasib in non-small-cell lung cancer harboring a KRASG12C mutation. N. Engl. J. Med. 387, 120–131 (2022).

9. N. London, R. M. Miller, S. Krishnan, K. Uchida, J. J. Irwin, O. Eidam, L. Gibold, P. Cimermančič, R. Bonnet, B. K. Shoichet, J. Taunton, Covalent docking of large libraries for the discovery of chemical probes. Nat. Chem. Biol. 10, 1066–1072 (2014).

10. G. Bianco, S. Forli, D. S. Goodsell, A. J. Olson, Covalent docking using autodock: Two-point attractor and flexible side chain methods: Covalent Docking with AutoDock. Protein Sci. 25, 295–301 (2016).

11. Y. Wu, C. L. Brooks Iii, Covalent docking in CDOCKER. J. Comput. Aided Mol. Des. 36, 563–574 (2022).

12. D. Toledo Warshaviak, G. Golan, K. W. Borrelli, K. Zhu, O. Kalid, Structure-based virtual screening approach for discovery of covalently bound ligands. J. Chem. Inf. Model. 54, 1941–1950 (2014).

13. S. De Cesco, S. Deslandes, E. Therrien, D. Levan, M. Cueto, R. Schmidt, L.-D. Cantin, A. Mittermaier, L. Juillerat-Jeanneret, N. Moitessier, Virtual screening and computational optimization for the discovery of covalent prolyl oligopeptidase inhibitors with activity in human cells. J. Med. Chem. 55, 6306–6315 (2012).

14. V. Katritch, C. M. Byrd, V. Tseitin, D. Dai, E. Raush, M. Totrov, R. Abagyan, R. Jordan, D. E. Hruby, Discovery of small molecule inhibitors of ubiquitin-like poxvirus proteinase I7L using homology modeling and covalent docking approaches. J. Comput. Aided Mol. Des. 21, 549–558 (2007).

15. Y. S. Tan, M. Chakrabarti, R. M. Stein, L. E. Prentis, R. C. Rizzo, T. Kurtzman, M. Fischer, T. E. Balius, Development of receptor desolvation scoring and covalent sampling in DOCK 6: Methods evaluated on a RAS test set. J. Chem. Inf. Model., doi: 10.1021/acs.jcim.4c01623 (2025).

16. M. Rachman, A. Scarpino, D. Bajusz, G. Pálfy, I. Vida, A. Perczel, X. Barril, G. M. Keserű, DUckCov: A Dynamic Undocking-based virtual screening protocol for covalent binders. ChemMedChem 14, 1011–1021 (2019).

17. S. Zhang, J. Tan, Z. Lai, Y. Li, J. Pang, J. Xiao, Z. Huang, Y. Zhang, H. Ji, Y. Lai, Effective virtual screening strategy toward covalent ligands: identification of novel NEDD8-activating enzyme inhibitors. J. Chem. Inf. Model. 54, 1785–1797 (2014).

18. A. Shraga, E. Olshvang, N. Davidzohn, P. Khoshkenar, N. Germain, K. Shurrush, S. Carvalho, L. Avram, S. Albeck, T. Unger, B. Lefker, C. Subramanyam, R. L. Hudkins, A. Mitchell, Z. Shulman, T. Kinoshita, N. London, Covalent docking identifies a potent and selective MKK7 inhibitor. Cell Chem. Biol. 26, 98–108.e5 (2019).

19. C. I. Nnadi, M. L. Jenkins, D. R. Gentile, L. A. Bateman, D. Zaidman, T. E. Balius, D. K. Nomura, J. E. Burke, K. M. Shokat, N. London, Novel K-Ras G12C switch-II covalent binders destabilize Ras and accelerate nucleotide exchange. J. Chem. Inf. Model. 58, 464–471 (2018).

20. E. A. Fink, C. Bardine, S. Gahbauer, I. Singh, T. C. Detomasi, K. White, S. Gu, X. Wan, J. Chen, B. Ary, I. Glenn, J. O’Connell, H. O’Donnell, P. Fajtová, J. Lyu, S. Vigneron, N. J. Young, I. S. Kondratov, A. Alisoltani, L. M. Simons, R. Lorenzo-Redondo, E. A. Ozer, J. F. Hultquist, A. J. O’Donoghue, Y. S. Moroz, J. Taunton, A. R. Renslo, J. J. Irwin, A. García-Sastre, B. K. Shoichet, C. S. Craik, Large library docking for novel SARS-CoV-2 main protease non-covalent and covalent inhibitors. Protein Sci. 32, e4712 (2023).

21. A. Scarpino, G. G. Ferenczy, G. M. Keserű, Comparative evaluation of covalent docking tools. J. Chem. Inf. Model. 58, 1441–1458 (2018).

22. M. Gao, A. F. A. Moumbock, A. Qaseem, Q. Xu, S. Günther, CovPDB: a high-resolution coverage of the covalent protein-ligand interactome. Nucleic Acids Res. 50, D445–D450 (2022).

23. X.-K. Guo, Y. Zhang, CovBinderInPDB: A structure-based covalent binder database. J. Chem. Inf. Model. 62, 6057–6068 (2022).

24. H. Du, X. Zhang, Z. Wu, O. Zhang, S. Gu, M. Wang, F. Zhu, D. Li, T. Hou, P. Pan, CovalentInDB 2.0: an updated comprehensive database for structure-based and ligand-based covalent inhibitor design and screening. Nucleic Acids Res. 53, D1322–D1327 (2024).

25. C. Wen, X. Yan, Q. Gu, J. Du, D. Wu, Y. Lu, H. Zhou, J. Xu, Systematic studies on the protocol and criteria for selecting a covalent docking tool. Molecules 24, 2183 (2019).

26. N. Huang, B. K. Shoichet, J. J. Irwin, Benchmarking sets for molecular docking. J. Med. Chem. 49, 6789–6801 (2006).

27. M. M. Mysinger, M. Carchia, J. J. Irwin, B. K. Shoichet, Directory of useful decoys, enhanced (DUD-E): better ligands and decoys for better benchmarking. J. Med. Chem. 55, 6582–6594 (2012).

28. R. M. Stein, Y. Yang, T. E. Balius, M. J. O’Meara, J. Lyu, J. Young, K. Tang, B. K. Shoichet, J. J. Irwin, Property-unmatched decoys in docking benchmarks. J. Chem. Inf. Model. 61, 699–714 (2021).

29. M. R. Bauer, T. M. Ibrahim, S. M. Vogel, F. M. Boeckler, Evaluation and optimization of virtual screening workflows with DEKOIS 2.0--a public library of challenging docking benchmark sets. J. Chem. Inf. Model. 53, 1447–1462 (2013).

30. S. G. Rohrer, K. Baumann, Maximum unbiased validation (MUV) data sets for virtual screening based on PubChem bioactivity data. J. Chem. Inf. Model. 49, 169–184 (2009).

31. J. Abramson, J. Adler, J. Dunger, R. Evans, T. Green, A. Pritzel, O. Ronneberger, L. Willmore, A. J. Ballard, J. Bambrick, S. W. Bodenstein, D. A. Evans, C.-C. Hung, M. O’Neill, D. Reiman, K. Tunyasuvunakool, Z. Wu, A. Žemgulytė, E. Arvaniti, C. Beattie, O. Bertolli, A. Bridgland, A. Cherepanov, M. Congreve, A. I. Cowen-Rivers, A. Cowie, M. Figurnov, F. B. Fuchs, H. Gladman, R. Jain, Y. A. Khan, C. M. R. Low, K. Perlin, A. Potapenko, P. Savy, S. Singh, A. Stecula, A. Thillaisundaram, C. Tong, S. Yakneen, E. D. Zhong, M. Zielinski, A. Žídek, V. Bapst, P. Kohli, M. Jaderberg, D. Hassabis, J. M. Jumper, Accurate structure prediction of biomolecular interactions with AlphaFold 3. Nature 630, 493–500 (2024).

32. A. Stecula, R. Paul, K. Litchfield, S. E. Dalton, C. M. R. Low, C. R. Reis, M. Congreve, The rise of AlphaFold in drug design. Prog. Med. Chem. 64, 99–147 (2025).

33. A. Abdeldayem, Y. S. Raouf, S. N. Constantinescu, R. Moriggl, P. T. Gunning, Advances in covalent kinase inhibitors. Chem. Soc. Rev. 49, 2617–2687 (2020).

34. M. K. Gilson, T. Liu, M. Baitaluk, G. Nicola, L. Hwang, J. Chong, BindingDB in 2015: A public database for medicinal chemistry, computational chemistry and systems pharmacology. Nucleic Acids Res. 44, D1045–53 (2016).

35. B. Zdrazil, E. Felix, F. Hunter, E. J. Manners, J. Blackshaw, S. Corbett, M. de Veij, H. Ioannidis, D. M. Lopez, J. F. Mosquera, M. P. Magarinos, N. Bosc, R. Arcila, T. Kizilören, A. Gaulton, A. P. Bento, M. F. Adasme, P. Monecke, G. A. Landrum, A. R. Leach, The ChEMBL Database in 2023: a drug discovery platform spanning multiple bioactivity data types and time periods. Nucleic Acids Res. 52, D1180–D1192 (2024).

36. Q. Liu, Y. Sabnis, Z. Zhao, T. Zhang, S. J. Buhrlage, L. H. Jones, N. S. Gray, Developing irreversible inhibitors of the protein kinase cysteinome. Chem. Biol. 20, 146–159 (2013).

37. Z. Zhao, P. E. Bourne, Progress with covalent small-molecule kinase inhibitors. Drug Discov. Today 23, 727–735 (2018).

38. L. S. Rathod, P. S. Dabhade, S. N. Mokale, Recent progress in targeting KRAS mutant cancers with covalent G12C-specific inhibitors. Drug Discov. Today 28, 103557 (2023).

39. M. L. Verdonk, V. Berdini, M. J. Hartshorn, W. T. M. Mooij, C. W. Murray, R. D. Taylor, P. Watson, Virtual screening using protein-ligand docking: avoiding artificial enrichment. J. Chem. Inf. Comput. Sci. 44, 793–806 (2004).

40. J. J. Irwin, K. G. Tang, J. Young, C. Dandarchuluun, B. R. Wong, M. Khurelbaatar, Y. S. Moroz, J. Mayfield, R. A. Sayle, ZINC20-A free ultralarge-scale chemical database for ligand discovery. J. Chem. Inf. Model. 60, 6065–6073 (2020).

41. B. J. Bender, S. Gahbauer, A. Luttens, J. Lyu, C. M. Webb, R. M. Stein, E. A. Fink, T. E. Balius, J. Carlsson, J. J. Irwin, B. K. Shoichet, A practical guide to large-scale docking. Nat. Protoc. 16, 4799–4832 (2021).

42. I. S. Knight, S. Naprienko, J. J. Irwin, Enrichment Score: a better quantitative metric for evaluating the enrichment capacity of molecular docking models, arXiv [q-bio.QM] (2022). http://arxiv.org/abs/2210.10905.

43. M. M. Mysinger, B. K. Shoichet, Rapid context-dependent ligand desolvation in molecular docking. J. Chem. Inf. Model. 50, 1561–1573 (2010).

44. H. Park, P. Bradley, P. Greisen Jr, Y. Liu, V. K. Mulligan, D. E. Kim, D. Baker, F. DiMaio, Simultaneous optimization of biomolecular energy functions on features from small molecules and macromolecules. J. Chem. Theory Comput. 12, 6201–6212 (2016).

45. T. E. Balius, Y. S. Tan, M. Chakrabarti, DOCK 6: Incorporating hierarchical traversal through precomputed ligand conformations to enable large-scale docking. J. Comput. Chem. 45, 47–63 (2024).

46. M. R. Masters, A. H. Mahmoud, M. A. Lill, Investigating whether deep learning models for co-folding learn the physics of protein-ligand interactions. Nat. Commun. 16, 8854 (2025).

47. L. A. Honigberg, A. M. Smith, M. Sirisawad, E. Verner, D. Loury, B. Chang, S. Li, Z. Pan, D. H. Thamm, R. A. Miller, J. J. Buggy, The Bruton tyrosine kinase inhibitor PCI-32765 blocks B-cell activation and is efficacious in models of autoimmune disease and B-cell malignancy. Proc. Natl. Acad. Sci. U. S. A. 107, 13075–13080 (2010).

48. H. Y. Estupiñán, A. Berglöf, R. Zain, C. I. E. Smith, Comparative analysis of BTK inhibitors and mechanisms underlying adverse effects. Front. Cell Dev. Biol. 9, 630942 (2021).

49. T. Barf, T. Covey, R. Izumi, B. van de Kar, M. Gulrajani, B. van Lith, M. van Hoek, E. de Zwart, D. Mittag, D. Demont, S. Verkaik, F. Krantz, P. G. Pearson, R. Ulrich, A. Kaptein, Acalabrutinib (ACP-196): A covalent Bruton tyrosine kinase inhibitor with a differentiated selectivity and in vivo potency profile. J. Pharmacol. Exp. Ther. 363, 240–252 (2017).

50. S. Passaro, G. Corso, J. Wohlwend, M. Reveiz, S. Thaler, V. R. Somnath, N. Getz, T. Portnoi, J. Roy, H. Stark, D. Kwabi-Addo, D. Beaini, T. Jaakkola, R. Barzilay, Boltz-2: Towards accurate and efficient binding affinity prediction, bioRxivorg (2025)p. 2025.06.14.659707.

51. M. D. Stanchina, S. Montoya, A. V. Danilov, J. J. Castillo, A. J. Alencar, J. C. Chavez, C. Y. Cheah, C. Chiattone, Y. Wang, M. Thompson, P. Ghia, J. Taylor, J. P. Alderuccio, Navigating the changing landscape of BTK-targeted therapies for B cell lymphomas and chronic lymphocytic leukaemia. Nat. Rev. Clin. Oncol. 21, 867–887 (2024).

52. F. Cameron, M. Sanford, Ibrutinib: first global approval. Drugs 74, 263–271 (2014).

53. Y. Y. Syed, Zanubrutinib: First approval. Drugs 80, 91–97 (2020).

54. A. Markham, S. Dhillon, Acalabrutinib: First global approval. Drugs 78, 139–145 (2018).

55. C. Labanca, E. A. Martino, E. Vigna, A. Bruzzese, F. Mendicino, G. Caridà, E. Lucia, V. Olivito, V. Manicardi, N. Amodio, A. Neri, F. Morabito, M. Gentile, Rilzabrutinib for the treatment of immune thrombocytopenia. Eur. J. Haematol. 115, 4–15 (2025).

56. A. Corrionero, X. Zhang, P. Alfonso, P. J. Morris, C. Klumpp-Thomas, C. Melani, C. McKnight, J. D. Phelan, D. Holland, K. Wilson, S. B. Hoyt, M. Roschewski, P. J. Tonge, W. Wilson, M. Ceribelli, L. M. Staudt, C. J. Thomas, An assessment of kinase selectivity, enzyme inhibition kinetics and in vitro activity for several bruton tyrosine kinase (BTK) inhibitors. ACS Pharmacol. Transl. Sci., doi: 10.1021/acsptsci.5c00412 (2025).

57. B. R. Lanning, L. R. Whitby, M. M. Dix, J. Douhan, A. M. Gilbert, E. C. Hett, T. O. Johnson, C. Joslyn, J. C. Kath, S. Niessen, L. R. Roberts, M. E. Schnute, C. Wang, J. J. Hulce, B. Wei, L. O. Whiteley, M. M. Hayward, B. F. Cravatt, A road map to evaluate the proteome-wide selectivity of covalent kinase inhibitors. Nat. Chem. Biol. 10, 760–767 (2014).

58. Q. Zhao, X. Ouyang, X. Wan, K. S. Gajiwala, J. C. Kath, L. H. Jones, A. L. Burlingame, J. Taunton, Broad-spectrum kinase profiling in live cells with lysine-targeted sulfonyl fluoride probes. J. Am. Chem. Soc. 139, 680–685 (2017).

59. S. Eid, S. Turk, A. Volkamer, F. Rippmann, S. Fulle, KinMap: a web-based tool for interactive navigation through human kinome data. BMC Bioinformatics 18, 16 (2017).

60. A. T. Bender, A. Gardberg, A. Pereira, T. Johnson, Y. Wu, R. Grenningloh, J. Head, F. Morandi, P. Haselmayer, L. Liu-Bujalski, Ability of bruton’s tyrosine kinase inhibitors to sequester Y551 and prevent phosphorylation determines potency for inhibition of Fc receptor but not B-cell receptor signaling. Mol. Pharmacol. 91, 208–219 (2017).

61. R. E. Joseph, N. Amatya, D. B. Fulton, J. R. Engen, T. E. Wales, A. Andreotti, Differential impact of BTK active site inhibitors on the conformational state of full-length BTK. Elife 9 (2020).

62. A. V. Sadybekov, V. Katritch, Computational approaches streamlining drug discovery. Nature 616, 673–685 (2023).

63. M. M. Attwood, D. Fabbro, A. V. Sokolov, S. Knapp, H. B. Schiöth, Trends in kinase drug discovery: targets, indications and inhibitor design. Nat. Rev. Drug Discov. 20, 839–861 (2021).

64. P. Cohen, D. Cross, P. A. Jänne, Kinase drug discovery 20 years after imatinib: progress and future directions. Nat. Rev. Drug Discov. 20, 551–569 (2021).

65. P. Škrinjar, J. Eberhardt, J. Durairaj, T. Schwede, Have protein-ligand co-folding methods moved beyond memorisation?, Bioinformatics (2025). https://www.biorxiv.org/content/10.1101/2025.02.03.636309v2.

66. J. Lyu, N. Kapolka, R. Gumpper, A. Alon, L. Wang, M. K. Jain, X. Barros-Álvarez, K. Sakamoto, Y. Kim, J. DiBerto, K. Kim, I. S. Glenn, T. A. Tummino, S. Huang, J. J. Irwin, O. O. Tarkhanova, Y. Moroz, G. Skiniotis, A. C. Kruse, B. K. Shoichet, B. L. Roth, AlphaFold2 structures guide prospective ligand discovery. Science 384, eadn6354 (2024).

67. M. Holcomb, Y.-T. Chang, D. S. Goodsell, S. Forli, Evaluation of AlphaFold2 structures as docking targets. Protein Sci. 32, e4530 (2023).

68. M. Karelina, J. J. Noh, R. O. Dror, How accurately can one predict drug binding modes using AlphaFold models? Elife 12 (2023).

69. J. Lyu, S. Wang, T. E. Balius, I. Singh, A. Levit, Y. S. Moroz, M. J. O’Meara, T. Che, E. Algaa, K. Tolmachova, A. A. Tolmachev, B. K. Shoichet, B. L. Roth, J. J. Irwin, Ultra-large library docking for discovering new chemotypes. Nature 566, 224–229 (2019).

70. F. Gentile, J. C. Yaacoub, J. Gleave, M. Fernandez, A.-T. Ton, F. Ban, A. Stern, A. Cherkasov, Artificial intelligence-enabled virtual screening of ultra-large chemical libraries with deep docking. Nat. Protoc. 17, 672–697 (2022).

71. A. A. Sadybekov, A. V. Sadybekov, Y. Liu, C. Iliopoulos-Tsoutsouvas, X.-P. Huang, J. Pickett, B. Houser, N. Patel, N. K. Tran, F. Tong, N. Zvonok, M. K. Jain, O. Savych, D. S. Radchenko, S. P. Nikas, N. A. Petasis, Y. S. Moroz, B. L. Roth, A. Makriyannis, V. Katritch, Synthon-based ligand discovery in virtual libraries of over 11 billion compounds. Nature 601, 452–459 (2022).

