## Supplementary Methods for "Discovery of novel covalent ligands with AlphaFold3"

#### Benchmark set development

##### *Curation of active compounds*

A challenging aspect in the curation of covalent active compounds and targets is the lack of explicit annotation of the binding mechanism in public databases. As a result, we use a series of filters to maximize the likelihood that the selected compounds act through a covalent binding mechanism. As a main filter, we focused the domain of COValid on Cys-binding acrylamides - Cysteines are the most targeted nucleophilic residue in covalent ligand discovery, and acrylamides are the most common covalent warhead used.

Raw data was retrieved from BindingDB (April 2024 data) as well as ChEMBL 33 (*1*),(*2*). For each of the two datasets, the following steps were taken to filter the compounds to a relevant set of putative actives:

1. Only compounds with exactly one substructure match of the acrylamide warhead were used (multiple acrylamide moieties may lead to ambiguity regarding the covalent attachment point).
2. The molecular weight and the number of rotatable bonds of the compounds were limited to no more than 500[Da] and 12 bonds, respectively. This limits the benchmark set to the regime relevant for performance measurement of current docking tools, which are known to generally perform poorly for larger molecules.
3. A strict threshold was enforced on activity annotations - we required that active compounds in COValid have at least one affinity metric annotation ($K_I$/$K_D$/$EC_{50}$/$IC_{50}$) not higher than 100 nM. We used a more strict threshold than the 1 μM affinity threshold used in DUD-E, in order to increase the likelihood that the curated actives act via a covalent mechanism, which is known to often yield higher affinity than non-covalent analogs (*3*).
4. Substitutions on the acrylamide - we allowed no substitutions on the acrylamide Cα atom. This mostly rules out cyanoacrylamides, which often bind in a covalent, reversible fashion (*4*, *5*). We wished to focus the domain of COVAlid to irreversible covalent binders. For

non-cyanoacrylamides with substitutions on the acrylamide Cα, confidence of a covalent binding mechanism is low. Regarding the acrylamide Cβ atom, we only allowed a privileged set of three substitutions, once again filtering out unusual substitutions, for which the mechanism confidence is low. These include methylation on the Cβ atom as well as two substitutions included in FDA-approved covalent drugs - Afatinib and Dacomitinib (*6*, *7*) (Fig. S1).

5. For compounds that appeared in the database with SMILES representations twice - once with stereochemical specification and another time without it, we selected the more informative SMILES for COVAlid, and discarded the other entry.
6. Duplicate removal - for pairs of compounds sharing both the same canonical SMILES representation as well as the same protein target (as determined by their Uniprot ID annotation), one of the two compounds was discarded. We do allow two active compounds annotated as binders of different protein targets in the set to share the same canonical SMILES.

Some database-specific filters were used to ensure the relevance of the curated data:

1. BindingDB – compounds lacking annotation of the Uniprot ID of their primary target were discarded. Compounds with multiple associated ChEMBL IDs were discarded as well. Compounds annotated with a number of protein chains larger than one, implying a biomolecular complex, were also removed from the curated set.
2. ChEMBL - similarly to DUD-E, we enforced a ChEMBL confidence score of 4 or higher, keeping ligands that are assigned to protein targets. (*3*). We filtered the compounds entries for human protein targets only, and discarded compounds whose target was not annotated as a single protein.

Next, the sets curated from both datasets were concatenated. Inter-database duplicate removal and stereochemistry filtration was conducted, as described above. Since source-text analysis was conducted in later stages, we discarded compounds that lacked annotation of a source text. As a final filter, we discarded compounds whose source text did not include any mention of covalency and/or irreversibility (we required that at least one of the following phrases appear in the source text - “irreversibly”, “irreversible”, or “covalent” not prefixed by “non-”).

##### *Actives - diverse subset selection*

In order to have COValid represent a large portion of chemical space while keeping it compact for iterative optimizations of docking protocol, we filtered large compound sets by topological diversity. For Cys attachment sites annotated with less than 100 compounds, all compounds were included in the final benchmark. In cases where the Cys site had more than 100 compounds, we selected a diverse subset based on chemical topology. Morgan fingerprints were calculated for all putative active compounds of the Cys site. These are circular fingerprints that encode the presence or absence of multiple substructures in the form of a bit vector. We used a radius of 2 (for each atom, substructures which include it are searched among atoms that are up to bonds apart from it), and a bit vector length of 2048. Next, the sphere exclusion clustering algorithm implemented in RDkit was used. This algorithm outputs a set of compounds whose maximal pairwise fingerprint Tanimoto similarity is set by a user-specific input. We enforced a maximal pairwise similarity of 0.6. If the clustering output included more than 100 compounds, we randomly selected 100 of the output compounds.

##### *Enumeration of active protomers*

For each active compound in the set, Dimorphite-DL (*8*) was used to enumerate putative protonation states at a pH range of 7.0-7.8. Next, each protonated version of the active was enumerated for putative stereoisomers via RDkit (at each stereocenter which does not have stereochemistry annotation specified in the original active compound). The protonated, stereochemically enumerated SMILES representations are the final representation of the actives in COValid, and are referred to as the active protomers.

##### *Assignment of actives to Cys sites*

The following protocol was conducted for each of the ten compound sets in COValid:

1. By manual inspection of the protein binding pocket, we listed indices of Cys sites within the pocket or in its vicinity.
2. For each active compound, we scraped its source text (publication or patent, according to source annotation of the activity assay), and searched for instances of each putative Cys site (given Cys with index X, a match to any of the following patterns was considered an instance of the Cys – “cysteineX”, “cysteine X”, “cysteine-X”, “cysX”, “cys X”, “cys-X” (capitalized or not), as well as “CX”.

3. For each Cys site, we considered all its putative Cys sites according to the results of the scraping protocol – if the source text of all actives included the same Cys site A or no mention of any of the putative Cys sites explicitly, we assigned all the actives to Cys site A. This was the case for seven of the nine proteins in COValid.
4. The eighth protein, EGFR, had a single compound with multiple Cys sites detected in the text, therefore the compound was discarded.
5. For the ninth protein, FGFR4, the scraping led to 15 actives annotated with C477, 37 annotated with C552, 4 compounds had both Cys indices mentioned, and 22 had no mentions of the putative Cys sites. The pocket of FGFR4 is known to have two Cys sites to which covalent binding was reported – C477 and C552 (PDB structures 5NWZ and 5NUD, respectively (*9*)). To avoid ambiguity in the assignment, we discarded compounds that mentioned both or none of the Cys indices, and assigned 15 actives to C477 and 37 actives to C552. As a sanity check, we read through the texts of a compound assigned by the scraping protocol to C552 and another assigned to C477, and confirmed that the publications indeed mention covalent binding to the respective sites (*10*),(*11*).

##### *Enumeration of database of non-covalent putative decoys*

For the active-decoy matching protocol of COValid, we first generated a database of putative non-covalent decoys. We used the ZINC20 database in SMILES representation as input (*12*). We filtered the dataset, keeping only compounds that have free animes as potential attachment points for an acrylamide warhead. For this purpose, we curated compounds that had one or more primary/secondary aliphatic/aromatic amines (we used four relevant SMARTS patterns from Open Babel, and conducted substructure matching via RDkit (*13*)). We used Dimorphite-DL (*8*) to enumerate protonation states of the curated compounds in a pH range of 7.0-7.8. Finally, we used RDkit to pre-calculate the physicochemical properties that will be matched to those of the actives. We selected the same six properties matched between actives and decoys in DUD-E (*3*) - molecular weight, number of hydrogen bond donors, number of hydrogen bond acceptors, the net charge, the LogP value, and the number of rotatable bonds.

##### *Active-decoy matching*

We implemented a protocol for the decoy design process, based on the decoy generation methodology described in DUDE-Z, with the incorporation of multiple adaptations required for covalent binders (*14*). For each set of actives corresponding to one of the ten Cys sites in COValid, we designed decoys in a two-step process – coarse-grained property matching in non-covalent space for the curation of potential decoys, followed by fine-grained matching in covalent space.

Step 1 – curation of potential property-matched decoys for each active protomer

In the first step, we conducted physicochemical property-matching in non-covalent space - matching the properties of a non-covalent analog of each covalent active protomer in COValid to that of non-covalent putative decoys from the ZINC20-based dataset. Morgan fingerprints were calculated for all COValid active protomers (radius = 2, 2048 bits) (*15*). Next, we used the pre-calculated physicochemical properties of the active protomers and those of the non-covalent dataset of putative decoys for coarse-grained property-matching across six properties (molecular weight, number of hydrogen bond donors, number of hydrogen bond acceptors, the net charge, the LogP value, and the number of rotatable bonds). For each active protomer, the initial matching filtered the decoy database by maximal active-decoys differences as listed below:

| Physiochemical property | Maximal active-decoy difference |
|---|---|
| Mw | 10.0 [Da] |
| logP | 0.5 |
| Number of hydrogen bond donors | 1 |
| Number of hydrogen bond acceptors | 1 |
| Number of rotatable bonds | 1 |
| Charge | Exact match |

If less than 7,500 matching decoy protomers were retrieved for the current active protomer, the matching conditions were relaxed iteratively multiple times, as listed below:

| Physiochemical property | Second iteration | Third iteration | Fourth iteration | Fifth iteration | Sixth iteration | Seventh iteration | Eighth iteration |
|---|---|---|---|---|---|---|---|
| Mw | 10.0 [Da] | 20.0 [Da] | 30.0 [Da] | 40.0 [Da] | 50.0 [Da] | 50.0 [Da] | 50.0 [Da] |
| logP | 0.5 | 0.7 | 0.9 | 1.1 | 1.2 | 1.4 | 1.5 |
| Number of hydrogen bond donors | 1 | 1 | 1 | 1 | 1 | 1 | 1 |
| Number of hydrogen bond acceptors | 1 | 1 | 1 | 1 | 1 | 1 | 1 |

| | | | | | | | |
|---|---|---|---|---|---|---|---|
| Number of rotatable bonds | 1 | 1 | 1 | 1 | 1 | 1 | 2 |
| Net charge | Exact match | Exact match | Exact match | Exact match | 1 | 1 | 1 |

We conducted these curation iterations until a total of 7,500 decoy protomers were matched to the current active protomers, or until all eight iterations resumed.

We wished to prioritize closely-matching decoys. Therefore, we considered all decoys retrieved for the current active protomer, and calculated the number of closely-matching decoys protomers by the following criteria:

| **Physiochemical property** | **Maximal active-decoy difference** |
|---|---|
| Mw | 10.0 [Da] |
| logP | 0.5 |
| Number of hydrogen bond donors | Exact match |
| Number of hydrogen bond acceptors | Exact match |
| Number of rotatable bonds | Exact match |

If the number of closely matching decoys was higher than 7,500, a subset of size 7,500 compounds was randomly selected. Otherwise, all closely matching decoys were selected, and a random subset of the other curated decoys, which match the active more loosely, were added to reach a total of 7,500 compounds (or less than that, if not enough decoys matched the actives).

Next, the putative decoys of the current actives were filtered by topological similarity, only keeping decoy protomers dissimilar to all active protomers of the current Cys site. Based on Morgan fingerprints (radius = 2, bits = 2048), any decoy protomer with a Tanimoto similarity higher than 0.3 to any active of the current Cys site was discarded from the set (using the sphere exclusion algorithm implemented in RDkit).

This entire process was conducted separately for each active protomer of the current Cys site, yielding multiple putative decoys for each active protomer. In the final part of the non-covalent step, we selected a diverse subset of all decoy protomers of the current Cys site. We aggregated the decoy protomers across all actives of the current Cys site, and used the RDkit sphere exclusion algorithm to select a subset, enforcing a maximal Tanimoto similarity of 0.5 between any two non-covalent decoys.

Step 2 – active-decoy matching in covalent space

In the second step of the active-decoy matching process, we conduct fine-grained property-matching between covalent versions of the ZINC20-based putative decoy protomers and the covalent active compounds of COValid.

For each Cys site, the putative decoys of all its active protomers were aggregated, and duplicated decoys were removed. The RDkit sphere exclusion clustering algorithm was used to select a diverse subset, enforcing a maximal pairwise Tanimoto similarity of 0.5 between all decoys protomers of the Cys site.

Next, similar to DUD-E, we set to assign 50 decoy protomers to each active protomer (*3*). Starting from the most stringent thresholds on the maximal differences between active-decoy properties and iteratively relaxing these ranges, we assigned property-matched decoy protomers to each active until reaching a total of 50 decoys. In this step, properties were matched in covalent space, matching between the covalent active protomer and covalent decoys that were generated on-the-fly, by attaching acrylamide warheads (along with the same Cβ substitutions of the current active) to the potential attachment sites of the decoy protomer (free amines) and calculating physicochemical properties. It should be noted that simply adding an acrylamide warhead to an active-decoy pair that match in non-covalent space does not guarantee property-matching in covalent space, since the value of some properties varies depending on the attachment point on the ligand (e.g. LogP). If less than 50 decoys were curated after the ninth iteration resumed, additional decoys were mapped to the active regardless of property-matching to reach a total of 50. If a decoy protomer was assigned to one active protomer, we did not consider it in the mapping to other active protomers of the same protein target.

| Property | first iteration | second iteration | third iteration | fourth iteration | fifth iteration | sixth iteration | seventh iteration | eighth iteration | ninth iteration |
|---|---|---|---|---|---|---|---|---|---|
| LogP | 0.2 | 0.4 | 0.5 | 0.6 | 1 | 1.2 | 1.4 | 1.6 | 1.6 |
| Mw | 10 | 20 | 30 | 35 | 40 | 50 | 60 | 70 | 70 |
| Hydrogen bond donors | 0 | 0 | 0 | 1 | 1 | 1 | 2 | 2 | 2 |
| Hydrogen bond acceptors | 0 | 0 | 0 | 1 | 1 | 1 | 2 | 2 | 2 |
| Charge | 0 | 0 | 0 | 0 | 0 | 0 | 1 | 2 | 2 |
| Rotatable bonds | 1 | 1 | 1 | 1 | 1 | 1 | 2 | 2 | 2 |

This entire two-step process yielded the final compound set in COValid, which includes a total of 874 active protomers and 37,919 decoy protomers (Fig. 1A and Dataset S3). The reported set includes explicit mapping from each active protomer to its 50 decoy protomers, enabling users to use a subset of COValid actives in their research, and easily retrieve the corresponding subset of the decoys.

##### *Re-mapping of protomers*

Upon visual inspection of the final protomers, we detected issues with some of the protomers, yielding unrealistic protonation states (some protomers contained charged carboxamide moieties). Therefore, we re-mapped the protomers in the following fashion:

1. Active and decoy protomers that contained a charged carboxamide moiety were discarded.
2. For each protein target in COValid, the remaining decoy protomers were redistributed among the remaining active protomers of the target in the following fashion. For each remaining decoy protomer, we iterated over allowed ranges for the matched properties (9 ranges, as shown above), going from most stringent to most relaxed. For each property range, we iterated over allowed active protomers. If the decoy protomer matched for all properties given the current allowed ranges, it was temporarily assigned to the active protomer. Finally, for each active protomer, if 45 or less decoy protomers were assigned, all assigned decoy protomers were mapped to the active protomer. Otherwise (more than 45 decoys assigned), we considered the top-50 best-matching decoys for each active. We located the subsets of decoys whose charge does not match exactly to that of the active protomer, with overall poor matching of the physiochemical properties (matched at sixth iteration or further). A random subset of these poorly matching decoys was discarded, such that the final mapping of decoys to the active would include 45 protomers.

***Curation of experimental structures***

For each of the ten COValid Cys attachment sites, we selected two *holo* experimental structures from the Protein Data Bank - one with a non-covalent ligand in the binding pocket, and another with a covalent binder attached to the Cys of interest (Dataset S1). We only considered x-ray structures corresponding to the Uniprot ID of the protein target. When available, structures of higher resolution were prioritized. For the covalent complexes, when possible, preference was given to structures with an acrylamide ligand. In the case of K-ras, only protein structures containing the G12C mutant were considered. The covalent and non-covalent *holo* structures of K-ras include two cofactors - an Mg ion as well as GDP. Both cofactors were included in all docking and structural predictions in this research. Since the two Cys sites of FGFR4 were annotated with the same non-covalent structure, COValid includes a total of 19 experimental structures across the ten Cys sites.

#### Enrichment analysis

##### *Adjusted LogAUC*

Docking enrichment refers to the ability of the docking tools to rank annotated active compounds higher than a background of inactive decoy compounds. The naive method to quantify enrichment is to iterate over a compound list sorted by docking score, plot the Receiver Operating Characteristic Curve (ROC) - the share of active protomers found as a function of the share of decoy protomers found - and report the Area Under the Curve (AUC). This metric ranks the ability of the docking tool to rank the entire database, but for practical relevance for prospective screening of large libraries, we wish to emphasize the performance in terms of early enrichment. Since only a small fraction of the virtual library can be progressed to synthesis and experimental validation, enrichment performance in the top fraction of the ranked list is much more crucial than performance down the sorted list. For this reason, the LogAUC value is often reported as a more informative enrichment metric for docking tools (*16*). The LogAUC value is the area under the curve of a semi-logarithmic version of the ROC curve, where the share of actives found is plotted against the log transformation the share of the decoys found, such that the calculated area is dictated by the top fraction more so than by the rest of the ranked list. In fact, this transformation places exponentially more weight on early rather than late enrichment (*17*). In this research, we report the adjusted LogAUC value, where the LogAUC corresponding to random enrichment (14.5%) is subtracted from the LogAUC of the docking tool, such that an adjusted LogAUC value of 0% corresponds to random enrichment, and a value of 85.5% corresponds to maximum enrichment (*18*). For each compound list, we sort it, and keep the best-scoring active protomer of each compound, along with all decoy protomers. Finally, we compute the adjusted LogAUC in a scheme based on the python script distributed with DUDE-Z (*14*). Similarly to DUDE-Z, we focus on the area under the LogAUC between 0.001 to 1.0, such that the calculation of the area spans three orders of magnitude.

##### *Weighted paired t-test*

When comparing two COValid runs of the docking tools, we used a paired t-test. Due to the discrepancy in the sizes of the groups (some Cys sites have much more actives and decoys than others), we weighted the effect of each protein according to the size of its compound list. In the case of DOCKovalent and DOCK6, the adjusted LogAUC value for the same PDB structure from two different docking runs were paired (20 pairs, across the 20 PDB structures of COValid). For

AutoDock, we paired the three replicates that we ran for each PDB structure (60 pairs). In 2D plots depicting the average CPU runtime per protomer as a function of the average adjusted LogAUC values of the COVAlid run, we plotted dashed lines connecting two data points if increasing the current exhaustiveness parameter led to an increase in the average adjusted LogAUC, and conducted the weighted paired t-test to study whether or not this difference was statistically significant. Throughout the figures in the SI, we mark the p-value (two-tailed) as follows: >0.05 - insignificant (ns), 0.01-0.05 - significant (*), 0.001-0.01 - highly significant (**), and <0.001 - very highly significant (***).

##### *Error bars for reported metrics*

For each run of COValid in the optimization figures, we report either just the average adjusted LogAUC, or both the average adjusted LogAUC and the average CPU runtime per protomer. For DOCK6 and DOCKovalent, each datapoint represents a single run of the relevant compound library docked to the corresponding Cys site. AutoDock, on the other hand, uses a genetic algorithm with a random element, therefore we report each data point as the average of three replicates - three repetitions of the same run with a different random seed.

1. Average CPU runtime per protomer - in the case of DOCK6 and DOCKovalent, the output contains the total runtime for batches of protomers, and not for each protomer separately. For each of the 20 Cys sites (10 attachment points with covalent and non-covalent PDB structures), we calculated an average runtime per monomer. Finally, we report an average of the 20 values, weighted by the number of protomers associated with each Cys site. The error bars correspond to the weighted standard deviation. For AutoDock, a similar calculation is used, but since each run was executed three times, the weighted average and standard deviation are calculated across 60 values (three replicates per run).
2. Average adjusted LogAUC - similarly to runtime, for DOCK6 and DOCKovalent we report a weighted average of the 20 Cys sites along with a weighted standard deviation, and for AutoDock we do the same across a total of 60 replicates.

### Covalent Docking

##### *Ligand generation*

AutoDock

RDkit was used to add hydrogens to the ligand. The ETKDGv3 conformer generation of RDkit was used to generate a 3D pose of the ligand (*19*). In cases where generation failed, a second attempt was made, using random coordinates as the initial coordinate, instead of the default rule-based RDkit generation of initial coordinates. The maximal number of embedding attempts was also increased. If this second attempt failed, stereochemical information was removed from the ligand, and a final attempt was made to generate a 3D conformer. If conformer generation succeeded, RDkit's implementation of the MMFF94s force field was used for minimization of the conformer (*20*). A python pipeline combining RDkit and Meeko was used to align the ligand and covalently connect the ligand conformer with the cysteine attachment point. RDkit was used to calculate partial Gasteiger charges (*21*). Finally, the cysteine-ligand complex conformer was formatted in the pdbqt format required as input for AutoDock.

DOCK6

In the first step, the SMILES representations of the COValid protomers (already enumerated for protonation states and stereochemistry) were modified with the attachment of two silicon atoms to the acrylamide Cβ atom (required for the attach-and-grow method of DOCK6). Next, We used CORINA or Molconvert (multiple ligand generation schemes were tested) to generate a single 3D conformer of the compound (*22*). If this failed, stereochemical information was removed from the compound, and a second attempt was made. Next, AMSOL was used to calculate partial charges and desolvation parameters. A python script distributed with DOCK6 was used to convert both silicon atoms to dummy atoms and set their charges to zero, as well as adding per-atom solvation values to the file. This scripts also adjusts the bond angle between the two dummy atoms and the first atom of the ligand (in the domain of COVAlid - the angle between the Cys Cβ, Cys Sɣ, and the Cβ atom of the acrylamide binder) to a user-specified value. We selected an ideal value of 109.5 degrees.

#### DOCKovalent

In the first step, the SMILES representations of the COValid compounds were modified by the attachment of a single silicon atom to the Cβ atom of the acrylamide warhead (required for the DOCKovalent pipeline). Afterwards, Molconvert or Corina were used to generate a 3D conformer of the compound. If this failed, stereochemical information was removed from the ligand, and a second attempt at conformer generation was made. AMSOL was used to calculate partial charges and desolvation parameters. Next, Omega was used to expand the single 3D conformer to multiple conformers (*23*). Different values of the maximal number of Omega conformers were tested in the paper. If the initial attempt failed, another attempt was made, setting the strict_atom flag to False (making Omega's atom typing less strict). If this also failed, a final attempt was made for the run of Omega, setting both strict_atom and strict_stereo to false (setting to false means that a random stereoisomer will be used in cases where no stereochemistry is specified). The output mol2 files were converted to db2 format by a DOCK3.7 script. Finally, a db2.gz file was generated, to be provided as input to the DOCK3.7 docking protocol.

##### *Docking run*

#### AutoDock

The flexible side chain method of AutoDock4.2.6 was used for covalent docking (*24*). Out of two AutoDock methods available for covalent docking, this method performed best in pose recuperation studies. In this method, a 3D conformation of the ligand is pre-generated with ideal geometry, and covalently attached to the nucleophilic residue with random torsions. During docking, AutoDock's method for the modeling of sidechain flexibility of user-specified residues was employed sampling only torsional degrees of freedom (rotation around the cysteine Cα-Cβ bond and Cβ-Sɣ bond, as well as the internal torsions of the ligand). The protein backbone is kept rigid during docking. The docking box was set with 100 grid points in each axis, with 0.375Å spacing between grid points. The docking box was centered at the geometric center of the covalent ligand from the experimental covalent complex in COValid. For docking to the non-covalent complexes of COValid, we first aligned the covalent complexes with the non-covalent complex. Next, we used the geometric center of the aligned ligand from the covalent complex as the docking box center.

The vast majority of reported kinase binders target the conserved ATP-binding pocket, and use an ATP mimetic substructure to form hydrogen bonding interactions with two canonical residues of the kinase hinge region, deep within the cleft formed between the two lobes of the kinase (*25*). Since nine of the ten Cys sites in COValid correspond to kinases, we set to use this prior knowledge of protein-ligand interactions to guide the binding pose prediction by AutoDock. First, we annotated the two canonical hydrogen-binding residues of the hinge for each of the kinases in COValid, which are known to space apart by a single residue (*26*) (Table S3). Next, we used AutoDock Bias (ADB), a built-in method in AutoDock which enables the incorporation of prior knowledge regarding probable interactions to the docking process (*27*). ADB was shown to improve docking poses prediction and enrichment in the non-covalent case, and we set to probe its relevance for covalent enrichment (*27*), (*28*). For each kinase, an ADB script was used to calculate geometrically ideal interaction sites - for each backbone amine, a single ideal position is calculated for the positioning of a ligand hydrogen bond acceptor atom for the formation of a hydrogen bond, and for each backbone carbonyl, five ideal positions for the placement of a hydrogen bond acceptor atom are calculated (Fig. S29). ADB calculates such ideal sites for sidechain atoms as well as the backbone atoms (amine and carbonyl), but we only used the latter in this research. ADB modifies the energy grid maps of relevant atoms types using Gaussian energy wells centered at the calculated ideal sites, rewarding binding poses that allow for the formation of the desired hydrogen bonds during the sampling process (*27*). The value of the energetic reward at the ideal locations and the decay radius around these positions are user-specified. We set the decay radius to 1Å.

#### DOCKovalent

DOCKovalent, a DOCK3.7-based method, was used for covalent docking (*29*). DOCKovalent samples ligand poses in the binding pocket, while enforcing ideal covalent bond lengths and angles. Throughout all calculations reported, the covalent bond length was fixed to 1.8Å and the covalent attachment angles were fixed to 109.5°. When docking to a protein structure from a covalent complex, the experimental covalent ligand was used as the ligand input required for DOCK (called “xtal-lig”). When docking a protein structure from a non-covalent complex, we first aligned the covalent experimental complex with the non-covalent experimental complex. Then, the aligned covalent ligand was used as the “xtal-lig” input for docking.

DOCK6

The Attach-and-Grow method of DOCK6 was used for covalent docking. In preparation of the docking input, the nucleophilic CYS residue was mutated to GLY. Docking grids were prepared for DOCK3.7, and then converted for DOCK6. The coordinates of the original CYS residue were used to generate the three spheres (clarify). Except where specified explicitly, the default docking parameters from the DOCK6 covalent tutorial were used for the covalent docking. Xtal-lig was treated in the same fashion as in DOCKovalent (https://github.com/tbalius/teb_docking_test_sets/wiki/2023.05.08.6OIM_covalent#Process-SMILES-with-RDKit).

**AlphaFold3**

##### *Genetic and template search*

First, we ran the data pipeline of AlphaFold3 once for each protein target in the benchmark set (CPU-based calculation). The input was the single-letter amino acid sequence of the protein chain, derived from the FASTA file of the corresponding PDB structure (the covalent complex structures from COValid were used). Template search and genetic search (multiple sequence alignment) were performed with default settings.

##### *Compound preparation and complex prediction*

We used the default method for covalent complex prediction in AF3. First, the acrylamide double bond was reduced to a single bond to produce the adduct version of the warhead. Next, RDkit was used to add hydrogens to the compound and generate a 3D conformer to be provided as the ligand input to AF3 (using the ETKDGv3 conformer generation method of RDkit). If this failed, stereochemical information was removed from the compound, and another embedding attempt was made. The final conformer underwent force field minimization (using MMFFOptimizeMolecule function of RDKit). Finally, a script distributed with AF3 was used to convert the RDkit molecule to the molecular input format required by RDkit – CCD mmCIF. When running the prediction, this CCD mmCIF file was provided as a ligand (via the “userCCD” field of the input json file), and the covalent bond was explicitly specified (via the “bondedAtomPairs“ field of the json input file), between the  of the specific cysteine binding site residue and the acrylamide  of the ligand.

##### *Inference step*

A single seed was provided, yielding 5 samples (5 predictions). The top-ranked prediction (according to the AF3 ranking score prediction, see details in section "AF3 predicted confidence metrics") was used for all further analysis of the output.

##### *Rosetta minimization and scoring*

We used Rosetta to minimize and score the AF3 predictions (*30*). For each AF3-predicted covalent complex, we first removed the Sγ atom, detaching the covalent bond between the ligand and the residue. Next, since the output of AF3 does not contain hydrogen atoms, we used RDkit to add hydrogens (with 3D coordinates) to each compound, based on the protonation state of the input COValid protomer. Finally, we minimized the 3D non-covalent complex using Rosetta (using the lbfgs_armijo_nonmonotone minimization algorithm, with a fractional tolerance of 0.001, such that the minimum function value after minimization is likely within 0.1% of the true local minimum value). The default Rosetta scoring function, REF2015, was used for scoring. An example of a Rosetta-minimized AF3 prediction is shown in Fig. S30. The entire computational process including ligand generation for AF3, AF3 structural prediction, and the custom Rosetta minimization protocol, succeeded for 97.4% of the COValid protomers, with the per-Cys-site success rate ranging from 95.1% (JAK3 C909) to 99.1% (BMX C496).

We used the Rosetta scoring of the minimized AF3 complexes as a ranking score for enrichment analysis, and calculated adjusted LogAUC values for all ten COValid nucleophilic attachment sites. In Fig. S2, the enrichment performance of the AF3-Rosetta pipeline is compared to that of the two leading configurations of covalent docking tools (by averaged adjusted LogAUC) - DOCKovalent and DOCK6.

##### *AF3 predicted confidence metrics*

Along with the predicted structural coordinates of the biomolecular complex, AF3 also outputs multiple confidence metrics. AF3 was trained to predict these metrics, which correspond to the confidence in structural accuracy. Listed are the confidence metrics used in this research:

- pTM (predicted template modeling score) - this metric is a prediction of the template modeling (TM) score (*31*), which describes overall structural accuracy based on alignment to the ground-truth structure. As per the AF3 manual, ptM > 0.5 indicates that the overall fold of the predicted complex may be similar to that of the ground-truth structure. pTM is a scalar in range 0-1 (higher score indicates higher confidence).
- Chain-specific PTM - within the context of a biomolecular complex, this is the pTM restricted to a specific chain.
- ipTM (interface predicted template modeling) - this is a variant of the pTM metrics that only considers the interface between chains. The AF3 manual indicates that ipTM > 0.8 is indicative of a high-quality prediction, while ipTM < 0.6 could indicate a failed prediction (prediction with ipTM in range 0.6-0.8 may be correct or incorrect). ipTM is a scalar in range 0-1 (higher score indicates higher confidence).
- Ranking score - this metric is used by AF3 to select the overall prediction out of five samples predicted for each input seed (we used a single seed in this research, generating five predictions). It takes into account the pTM score and the ipTM score, as well as structural metrics which report on the share of atoms experiencing clashes and the fraction of the predicted structure which is disordered.
- PAE (predicted aligned error) - the PAE is a matrix (dimensions are determined by the total number of AF3 tokens in the biomolecular complex) that estimates the error in the relative position and orientation between each pair of tokens in the structure. Higher PAE values indicate token pairs for which the confidence in their relative positions is lower. On top of the entire PAE matrix, AF3 explicitly outputs the minimal PAE values for each pair of chains (chain_pair_pae_min) as a matrix with dimensions matching the number of chains in the complex (2x2 for a protein-ligand chain complex without any cofactors). Element (i,j) of the matrix lists the minimal value from the PAE matrix across rows restricted to chain i and columns restricted to chain j. Setting to test the potential of this metric in the space of covalent protein-ligand interactions, we considered a single entry of the minimal PAE matrix, elem (1,0) representing the minimum across all pairwise predicted errors in the positions of protein tokens when aligned by ligand tokens (hereon referred to as the minimal PAE element). We refer to this value as mPAE. The minimal PAE element describes the minimal predicted error in positions of the protein target residues, when aligned to the ground-truth complex structure by reference frames of the ligand atoms.

- In our attempts to use these predicted metrics for library ranking and enrichment analysis, we noticed that most of the metrics do not yield sufficient resolution for a decisive ranking of the libraries. That is, in many cases large numbers of protomers are predicted with the same exact value of the confidence metrics. For example, BTK and FGFR1 each have 2411 and 857 protomers that are all predicted with the same exact pTM score - 0.94 and 0.88, respectively (Fig. S5). To test the ranking enrichment performance in spite of this limitation, we ranked the ten COValid protomer lists by the AF3-predicted metric, and then reported a range between two adjusted LogAUC values for each Cys site, using the annotations available in COValid. In the best-case ranking, in cases where multiple protomers were ranked with the same value, actives were ranked higher than decoys. In the worst-case - decoys were ranked higher.

##### *AF3 predicted confidence metrics*

When comparing AF3 predictions to experimental structures, we report the pocket-aligned RMSD, calculated in a scheme similar to the one described in the AF3 publication:

1. We define the ground-truth pocket as the set of all Cα atoms of the experimental structure that are within 10Å from any heavy atom of the ground-truth ligand.
2. The AF3-predicted is aligned to the ground-truth pocket via Pymol ("align" command").
3. RDkit is used to calculate the RMSD between heavy atoms of the ground-truth ligand and the predicted ligand.

### Experimental Validation

#### *In silico prospective screening*

We constructed a virtual library of compounds based on the Enamine REAL compound library. The Enamine REAL data is available as multiple Parquet files, split into categories by the number of heavy atoms in the compounds, with each file listing the SMILES representations of the corresponding compounds. 6.71 billion compounds in the set were filtered by applying the following criteria to each Parquet file:

1. Molecular weight is in the range of 250[Da]-500[Da], and the number of rotatable bonds is no more than 12.
2. The compound includes at least a single aliphatic amine, either primary or secondary.

After implementing these filters, 327M compounds remain, aggregated across all Parquet files. Next, we selected a diverse subset of the compounds. First, we selected a diverse subset of each property-filtered Parquet file using the Rdkit sphere exclusion algorithm (threshold parameter = 0.7, enforcing a minimal Tanimoto distance between cluster centroids; used the cluster centroids provided by the algorithm as a diverse subset; used 2048-bit Morgan fingerprints with a radius of 2). Concatenated across all the files, this yielded 1.27M diverse compounds. Next, for each heavy-atom-count category, the diverse subsets of all the corresponding property-filtered Parquet files were merged, and a diverse subset of this merged set was selected using the sphere exclusion algorithm once again (this time with a threshold of 0.4; 2048-bit Morgan fingerprints with a radius of 2). Aggregated across all heavy-atom-count categories, this yielded 887,569 compounds. Next, we converted this set of Enamine-based amine precursors to covalent compounds, by attaching an acrylamide warhead to any potential attachment point considered (primary/secondary aliphatic amines). This yielded 906,510 acrylamide compounds (some of the 887,569 precursors had multiple attachment points).

For the screening campaign, We used a similar scheme to that used in the benchmarking of AlphaFold3. For the data step (MSA and structural template search), the input included the amino acid protein sequence of BTK, taken from PDB entry 5P9J (BTK-ibrutinib covalent complex). For each compound in the library, a 3D conformer with ideal coordinates was generated via RDkit and provided as input. The covalent attachment points (SG atom of Cys481 and Cb atom of the acrylamide warhead of the ligand) were also specified at the input for the prediction step. Predictions were executed with a single seed, yielding five diffusion samples. The best ranking sample out of the five was considered as the AF3 prediction for the ligand (as determined by the 'ranking score' metric reported by AF3). In total, AF3 predictions were successfully generated for 899,443 out of 906,510 compounds in the virtual libraries (99.2%) . 440 of the covalent complex predictions yielded mPAE values not greater than 0.9[Å]. Since our goal was to discover novel BTK binders, we filtered the putative binders by their similarity to known BTK binders. We filtered out compounds with Tanimoto similarity greater than or equal to 0.35 to any known submicromolar BTK inhibitor in ChEMBL (ChEMBL version 35; considered compounds annotated with $K_I/K_D/EC_{50}/IC_{50}$, and an assay confidence score no less than 4; 2048-bit Morgan fingerprints with a radius of 2). 390 compounds remained after this filter. We manually inspected them (visualized via Pymol), curating compounds with diverse binding poses, and focusing on compounds that form hydrogen bonds with the hinge region of BTK. Finally, we ordered 15 compounds for custom synthesis by Enamine, 13 of which were successfully synthesized.

##### *Intact protein LC/MS*

The LC/MS runs were performed on a Waters ACQUITY UPLC class H instrument, in positive ion mode using electrospray ionization. UPLC separation used a C4-BEH column (300 Å, 1.7 μm, 21 mm × 100 mm). The column was held at 40 °C and the autosampler at 10 °C. Mobile phase A was 0.1% formic acid in water, and mobile phase B was 0.1% formic acid in acetonitrile. The run flow was 0.4 mL/min. The gradient used was 1% B for 2 min, increasing linearly to 80% B for 2.5 min, holding at 80% B for 0.5 min, changing to 20% B in 0.2 min, and holding at 1% for 0.8 min. The MS data were collected on a Waters SQD2 detector with an m/z range of 2–3071.98 at a range of 600–1900 m/z. The desolvation temperature was 500 °C with a flow rate of 800 L/h. The voltages used were 1.00 kV for the capillary and 24 V for the cone. MassLynx version 4.2 was used to operate the LCMS and analyze the data. Raw data were processed using openLYNX and deconvoluted using MaxEnt with a range of 28000 : 36000 Da and a resolution of 1 Da/channel.

##### *Initial binding experiments*

Compounds were prepared at 10 mM stocks in DMSO. 1 μl of molecule stock was mixed with 49 μl of 1 μM BTK-KD (kinase domain) in HEPES 25 mM pH = 7.5, 50 mM NaCl (giving 200 μM molecule). Samples were incubated for either 2 hours or overnight at 25°C, and reactions were stopped by mixing 20 μl of sample with an equal volume of 20% acetonitrile + 0.25% TFA, followed by injection to intact protein LCMS. Dose response and time course experiments were conducted using the same procedure.

##### *Kinase activity assay and kinome panel*

Assays were conducted by Reaction Biology. The enzyme was diluted to 3 nM in reaction buffer (20 mM HEPES pH 7.5, 10 mM MgCl2, 1 mM EGTA, 0.01% Brij35, 0.02 mg/ml BSA, 0.1 mM Na3VO4, 2 mM DTT, 1% DMSO) and incubated with the compounds at different concentrations from DMSO stocks using acoustic dispensing (Echo 550) prior to addition of substrate for 60 minutes at room temperature. The enzyme was then incubated for 2 hours with 0.2 mg/ml pEY substrate and 33p-labeled ATP. Kinase activity was detected radiometrically by the P81 filter-binding method. ATP concentrations of 20 μM and 50 μM were used for WT BTK and C481S BTK, respectively.

For the kinome panel testing, different enzymes (with added cofactors if needed) were incubated with 0.3 μM compound and tested using the same radiometric assay.

##### *GSH assay*

Reduced glutathione (GSH) was dissolved to 100 mM and titrated to pH = 8 using NaOH. The assay buffer contained 50% acetonitrile, 25 mm NaPi pH = 8 with 400 μM 3-acetamidobenzoic acid as an internal reference. The GSH was diluted to 5 mM in the buffer, and 245 μl of diluted GSH was mixed with 5 μl of 10 mM molecule (dissolved in DMSO). The reaction was incubated at 25°C and 10 μl samples were stopped at different times by mixing with 40 μl of 0.1% TFA in 10% acetonitrile to stop the reaction, followed by injection of 10 μl to LCMS. The reaction products were run using the following gradient: 1% acetonitrile from 0 to 1 minutes, then rise to 95% acetonitrile from 1 minutes to 5.5 minutes, followed by 0.5 minutes at 95% acetonitrile and then flushing the column with 1% acetonitrile for 1 minutes. All solvents contain 0.1% formic acid. Mass spectra were collected at 80-2500 m/z at positive ionization. The areas of the nonreacted molecule in the UV chromatogram were integrated and corrected against the internal reference to measure reaction rates.

##### *DSF assay*

BTK-KD was diluted to 5 μM in HEPES 25 mM pH = 7.5, 150 mm NaCl. 80 μl of protein solution was mixed with 1 μl of 1.6 mM molecule solution (giving 4-fold excess of molecule). The samples were incubated at 25°C overnight and full labeling was confirmed using LCMS. To each sample SYPRO orange dye was added to a final concentration of X5, and 3 replicates of 22 μl were added to a 96-well plate. Melting curves were measured using StepOne Plus instrument using FAM as the target and ROX as the passive reference. The derivative reporter was used to determine the melting point.

##### *phospho-BTK assay*

Mino cells were treated for 1 h with a dose-response of the indicated compounds. Following cells treatment, BTK phosphorylation was induced with 10 μg/ml anti-human IgM (Jackson ImmunoResearch, 109-006-129) for 10 min at 37 degrees. The cells were harvested, washed with ice-cold PBS, and lysed with RIPA buffer (sigma, R0278) supplemented with protease and phosphatase inhibitors. Protein concentration was determined using BCA protein assay (Thermo Fisher Scientific, 23225). Samples containing 20 μg of total protein were prepared with 4x LDS sample-buffer (GeneScript, M00676), and were then resolved on 4-20% Bis-Tris gel (GeneScript, surePAGE). Proteins were separated by electrophoresis and were then transferred to a nitrocellulose membrane (Bio-Rad, 1704158) using the Trans-Blot Turbo system (Bio-Rad). The membrane was blocked with 5% BSA in TBS-T (w/v) for 1 h at room temperature, washed four times for 5 min with TBS-T, and incubated with the following primary antibodies: rabbit anti-phospho-BTK (#87141s, Cell-Signaling, 1:1000, overnight at 4 °C), mouse anti-BTK (#56044s, Cell-Signaling, 1:1000, 1 h at room temperature), and mouse anti-β-actin (#3700, Cell-Signaling, 1:1000, 1 h at room temperature). Membrane was washed four times for 5 min with TBS-T, and incubated with the corresponding HRP-linked secondary antibody (mouse #7076/rabbit #7074, Cell-Signaling) for 1 h at room temperature. SuperSingal West Pico PLUS chemiluminescent substrate (Thermo Fisher Scientific, 34580) was used to detect HRP activity. The membrane was stripped using Restore stripping buffer (Thermo Fisher Scientific, 21059) after each secondary antibody before blotting with the next one.

##### *Proteomics*

Mino cells were treated for 1 h with DMSO 0.1% or 1 μM of YS1, followed by 45 min treatment with either 1 μM Probe 4 or 2 μM XO44, in 4 replicates for each condition. The cells were harvested, washed with ice-cold PBS, and lysed with RIPA buffer (sigma, R0278) supplemented with protease inhibitor cocktail (Sigma, P8340). Protein concentration was determined using BCA protein assay (Thermo Fisher Scientific, 23225). 500 μg samples in a total volume of 200 μl were prepared. A click reaction was performed in a final volume of 250 μl, using final concentrations of 100 μM biotin-azide, 4.8 mM THPTA, 0.96 mM CuSO4 and 8 mM sodium ascorbate. Samples were reacted for 1.5 h at room-temperature, followed by methanol-chloroform precipitation. 250

µl Chloroform, 750 µl water, and 1 ml methanol were added and the samples were vortexed thoroughly and spun-down for 10 min at 4 degrees. The top layer was aspirated, followed by addition of 1 ml methanol. Samples were vortexed and spun-down again for 10 min at 4 degrees, the solution was removed, and the pellet was air-dried. The dry pellets were re-suspended in 250 µl of 2.5 % SDS in PBS with sonication (5 pulses, 2 sec on/ 2 sec off, 25% amplitude). Samples were then reduced with 1.25 µl of 1 M DTT (40 min at 37 degrees), alkylated with 4.6 µl of 0.8 M Iodoacetamide (40 min at room-temperature, protected from light), and then another 7.5 µl of 1 M DTT were added. The samples were diluted 20-fold with PBS, and 20 µl of PBS pre-washed Streptavidin beads were added (Cytiba, Streptavidin Sepharose High Performance, 17511301). The samples were rotated for 3 h in room-temperature. The samples were spun-down (2000 g, 3 min), the supernatant was removed and the beads were washed x3 times with 0.1% SDS in PBS, x5 times with 20% methanol, twice with 2M NaCl in PBS, once with PBS and x3 times with water. The beads were transferred to new tubes with 200 µl of 100 mM Ammonium bicarbonate, 1 µl of 0.5 µg/µl sequencing-grade trypsin were added (Promega, V5111), and samples were incubated overnight at 37 degrees with shaking. The samples were spun-down, and the supernatant was collected to new tubes. The beads were then washed with 200 µl of 100 mM Ammonium bicarbonate + 2M NaCl, which were combined with the previous supernatant. TFA was added to a final concentration of 0.1% to each sample, and the samples were desalted by Oasis (Waters) and dried using speed-vac.

The samples were dissolved in 30 µl of 0.1% formic acid in 3% acetonitrile, and 3 µl were injected per run. Samples were analyzed using EASY-nLC 1200 nano-flow UPLC system, using PepMap RSLC C18 column (2 µm particle size, 100 Å pore size, 75 µm diameter × 50 cm length), mounted using an EASY-Spray source onto an Exploris 240 mass spectrometer. µLC/MS-grade solvents were used for all chromatographic steps at 300 nL/min. The mobile phase was: (A) H2O + 0.1% formic acid and (B) 80% acetonitrile + 0.1% formic acid. Peptides were eluted from the column into the mass spectrometer using the following gradient: 1–40% B in 160 min, 40–100% B in 5 min, maintained at 100% for 20 min, 100 to 1% in 10 min, and finally 1% for 5 min. Ionization was achieved using a 1800 V spray voltage with an ion transfer tube temperature of 275 °C. MS1 spectra were collected at a resolution of 60,000 at 200 m/z, a mass range of 370–1450 m/z, normalized AGC of 300%, and the maximum injection time was set to 20 ms. Data were collected by data-independent acquisition, with MS2 resolution set to 30,000, with 32 isolation windows of 17 Da with a 1 Da overlap covering precursor m/z of 390-920, maximum injection time of 50 ms, and HCD collision energy at 27%. Four samples were analyzed per condition. The data were processed using DIA-NN (version 2.2). First, a theoretical spectral library was generated from a fasta file containing the human proteome, streptavidin, trypsin and common contaminants. The number of missed cleavage was set to 1. The data were then analyzed against this spectral library using the following additional settings: Qvalue = 0.001; --matrix-qvalue [10] --matrix-spec-q [10] --restrict-fr. The rest of the settings were standard. The resulting data was analyzed using Perseus as follows: Intensity values were converted to Log2 values, the different quadruplicates were grouped and any protein for which at least 3 valid intensity values were measured for at least one of the groups was discarded from the data (968 proteins out of 4384). Missing values were then replaced with a value of 14, which corresponds to the lower limit of the measured intensity distribution. The groups were then compared using two sided t test, and the calculated differences and P-values were used to plot the volcano plots. To determine which proteins were pulled down by the probe and competed by YS1, we filtered the data to contain only proteins for which the Log2 difference between the probe treated set and DMSO-treated set was > 2 with a P-value < 0.01, and for which the Log2 difference between the probe treated set and the competed set was > 2 with a P–value < 0.01.

##### *X-Ray Crystallography*

Protein Expression and Purification. The procedures for bacterial expression and purification of the murine BTK kinase domain (residues 382−659) and the BTK kinase domain (residues 396−659) containing six activation-loop mutations (L542M, S543T, V555T, R562K, S564A, and P565S) have been described previously (32, 33). Both constructs include the Y617P mutation to enable efficient bacterial expression.

Crystallization. Protein/compound solutions in 20 mM Tris, pH 8, 150 mM NaCl, and 10% glycerol were prepared by incubating purified Btk proteins with 1 mM of the respective compounds and 10% DMSO on ice for 30 min. Co-crystals of the BTK kinase domain with YS1 and YS2 were obtained by mixing protein/compound solutions at 16 mg/mL with a reservoir solution consisting of 0.1 M imidazole, pH 7.0, and 18% PEG 3350 at a 1:1 ratio at 4 °C. Co-crystals of the BTK kinase domain carrying the six activation-loop mutations with YS3 were obtained by mixing protein/compound solutions at 18 mg/mL with a reservoir solution of 0.1 M sodium citrate, pH 5.5, and 20% PEG 3000. The crystals were harvested in a cryoprotectant solution consisting of the reservoir solution supplemented with 20% glycerol and flash-frozen in liquid nitrogen.

Data Collection, Structure Determination, and Refinement. X-ray diffraction data were collected at the Advanced Photon Source (APS), Beamline 24-ID-E Northeastern Collaborative Access

Team (NE-CAT). Data sets were indexed, merged, and scaled using autoPROC (Global Phasing) (*34–36*). Structures of protein/compound complexes were solved by molecular replacement with PHASER (*37*) using an unpublished BTK kinase domain structure as the search model. Three-dimensional ligand structures were generated using Grade Web Server (*38*). Ligand restraints were generated with eLBOW (*39*) and subsequently adjusted based on PDB validation feedback. Structure refinement was carried out in Phenix (*40*), and model building was performed with Coot (*41*). Additional restraints defining the covalent linkage between the protein and ligand were introduced in Phenix to ensure proper geometric representation. Crystallographic data collection and refinement statistics are provided in Table S?. Atomic coordinates and structure factors have been deposited into the Protein Data Bank under accession codes: 9ZLJ (BTK/YS1) and 9ZLM (BTK/YS2).

### Supplementary Results

##### Optimization of physics-based covalent docking tools

For AutoDock, increasing the number of genetic algorithm iterations from 10 to 40 did not significantly improve enrichment (average adjusted LogAUC of 6.6±7.4% and 5.5±6.9%, respectively; Fig. S31) nor did increasing the diversification of the initial 3D conformers by sampling four conformers with ±10º around the ideal covalent bond angles (average adjusted LogAUC of 6.6±7.4% and 6.0±7.0% using the default scheme and the angle diversification scheme, respectively; Fig. S32) nor did increasing the maximal number of energy evaluations throughout the run from 50K up to 5M (despite the latter resulting in approximately 65-fold longer run time; Fig. S33). What ended up significantly improving the performance of AutoDock, was the incorporation of knowledge about the binding poses via AutoDock Bias (*27*). By explicitly biasing the docking to favor poses that form canonical hydrogen bonds with the kinase hinge residues the performance improved significantly (7.0±7.6% vs. 22.7±11.2% with the maximal bias parameter used; Fig. S34).

For DOCKovalent, we were able to find several parameter modifications that improved enrichment. Some were related to changes in the docking setup that are performed before the run such as changing the initial ligand conformer generator (Fig. S35) from CORINA (*42*) (weighted average adjusted LogAUC = 5.9±5.2%) to MolConvert (*43*) (9.0±4.9% ; p=0.0014). Using a protocol (*16*) to modify the dielectric at the protein-solvent interface (Fig. S36) improved performance for some targets, but did not lead to a significant increase to average enrichment across COValid (weighted average adjusted LogAUC of 9.0±4.9% and 9.9±4.8% for the default protocol and the modified protocol, respectively; p=0.17).

Other parameters relate to the docking of each ligand and can therefore affect run-time significantly. Because ultimately billions of ligands may be docked, run-time differences can be crucial. An example for such a parameter is the magnitude of the range sampled around an ideal bond angle. The enrichment across the benchmark without deviation from the ideal covalent bond angle was 6.9±4.6% with an average run time of only 0.13±0.03 sec. per ligand (Fig. 2A). Increasing the sampled deviation to ±5º significantly increased the enrichment to 9.1±4.5% ($p=0.0002$) but also increased the run-time to 2.6±0.7 sec. on average. Increasing the value to ±10º improved performance further to 9.9±4.8% ($p=0.004$) with a runtime of 8.4±2.3 sec. Increasing to ±20º was no longer significant ($p=0.1053$), despite tripling the run-time (28.9±8.3 sec). The step-size of sampling the bond angle also had significant effects on enrichment (Fig. S37).

Parameters that did not significantly influence the performance (but did affect run-time) included the maximal number of ligand conformers generated for each ligand (Fig. S38), and the range of covalent bond length that was sampled (Fig. S39).

For DOCK6, CORINA slightly outperformed MolConvert for generation of input conformers (6.7±6.3% vs. 5.1±5.2%, respectively; $p=0.086$; Fig. S40). Adjusting the protein-solvent dielectric significantly improved enrichment (from 6.7±6.3% to 11.9±10.0%, ; $p=0.0006$; Fig. S41). Conversely, modifying the pruning parameter of the attach-and-grow algorithm had little effect (Fig. S42) and adjusting the step size of covalent bond dihedral angle showed no effect (Fig. S43).

We are aware this is not an exhaustive exploration and do not claim to have found the optimal setup for either of these tools, but hope the process illustrates the utility of COValid in guiding such an optimization.

#### Boltz-2 predictions for DUDE-Z

Using Boltz-2 (*44*), we conducted non-covalent complex predictions for the entire DUDE-Z set, and compared enrichment with AF3 predictions, sorted by mPAE. Boltz-2 provides two main metrics of affinity predictions - a binary metric ("affinity_probability_binary", in range [0,1]) for distinguishing better binders and decoys, and a continuous metric aimed at predicting the actual affinity value ("affinity_pred_value"; predicts a binding affinity value as $\log_{10}(IC_{50})$, derived from an $IC_{50}$ value measured in μM). Boltz-2 was run with default settings ("boltz predict"), using the "use_msa_server" flag for multiple sequence alignment. Sorting the 43 DUDE-Z libraries by either one of these two metrics yield average enrichment on par with that of AF3 mPAE-based ranking (average adjusted LogAUC values of 52.1%±20.0%, 51.3%±20.6%, and 55.3%±22.4% for AF3-mPAE, Boltz-2 continuous predictions, and Boltz-2 binary prediction, respectively; Fig. S14). In the context of the non-covalent DUDE-Z set, sorting compound libraries by the leading AF3-predicted confidence metric, which were not directly designed to report on potency or binding affinity, is approximately as good as sorting them according to the output of a model that was specifically trained on binding affinity data to make such predictions. Across the 43 targets, Boltz-2-based enrichment is strongly correlated with AF3-mPAE-based enrichment (Fig. S14; Pearson correlation coefficients of 0.74 and 0.66 with p-values of 1.6E-8 and 1.4E-6 for correlations of AF3-mPAE with Boltz-2 binary and continuous affinity predictions, respectively).

### Supplementary Figures

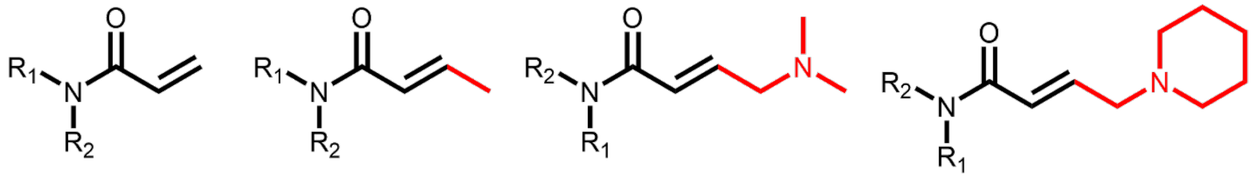

**Figure S1. The acrylamide electrophile and allowed substitutions on the acrylamide Cβ of COValid actives.**

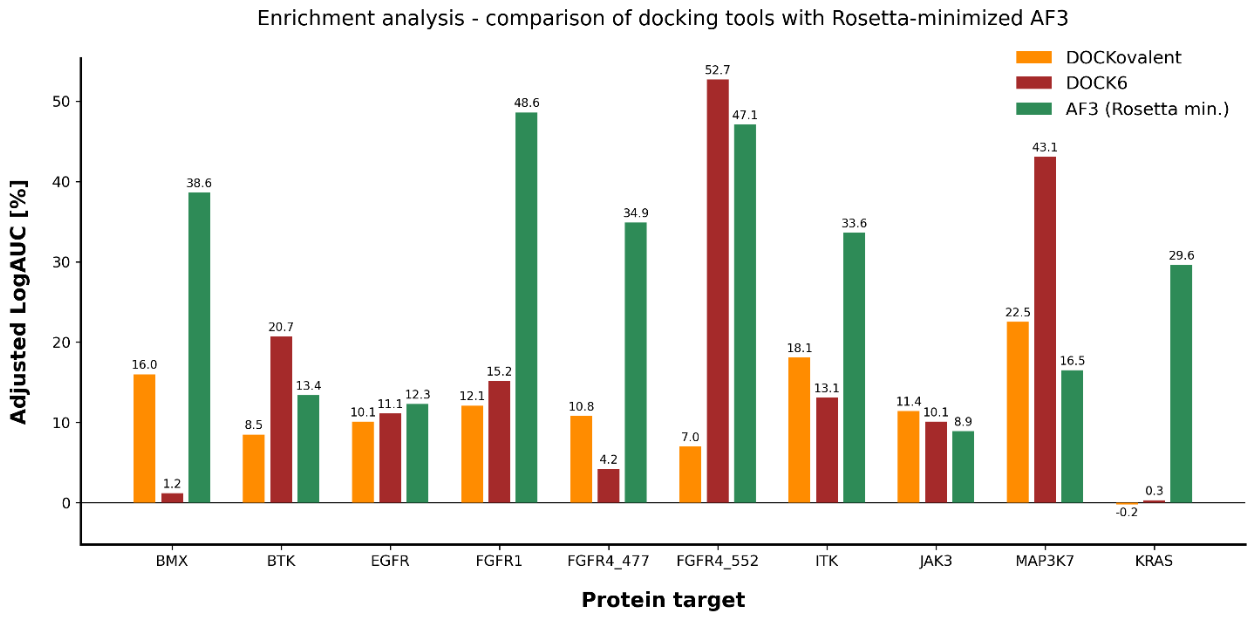


**Figure S2. Comparison of AI-based and docking-based covalent enrichment.** The plot shows the adjusted LogAUC values for the ten Cys sites of COValid, based on the docking score output of the two physics-based docking tools with leading configurations for enrichment performance (docking to the covalent PDB complexes of COValid), compared with those from sequence-based AF3 prediction followed by Rosetta minimization and scoring. For five of the nucleophilic attachment sites, Rosetta-minimized AF3 significantly outperforms the covalent docking tools. Fixed parameter values - DOCK6: Corina for conformer generation, protein preparation with thin spheres, pruning parameter = 50, covalent dihedral sampling parameter = 10°. DOCKovalent: MolConvert for generation of the initial conformer, protein preparation with spheres, maximal number of Omega conformers = 200, angle range = 10°, angle step = 2.5°, no sampling of bond length range.

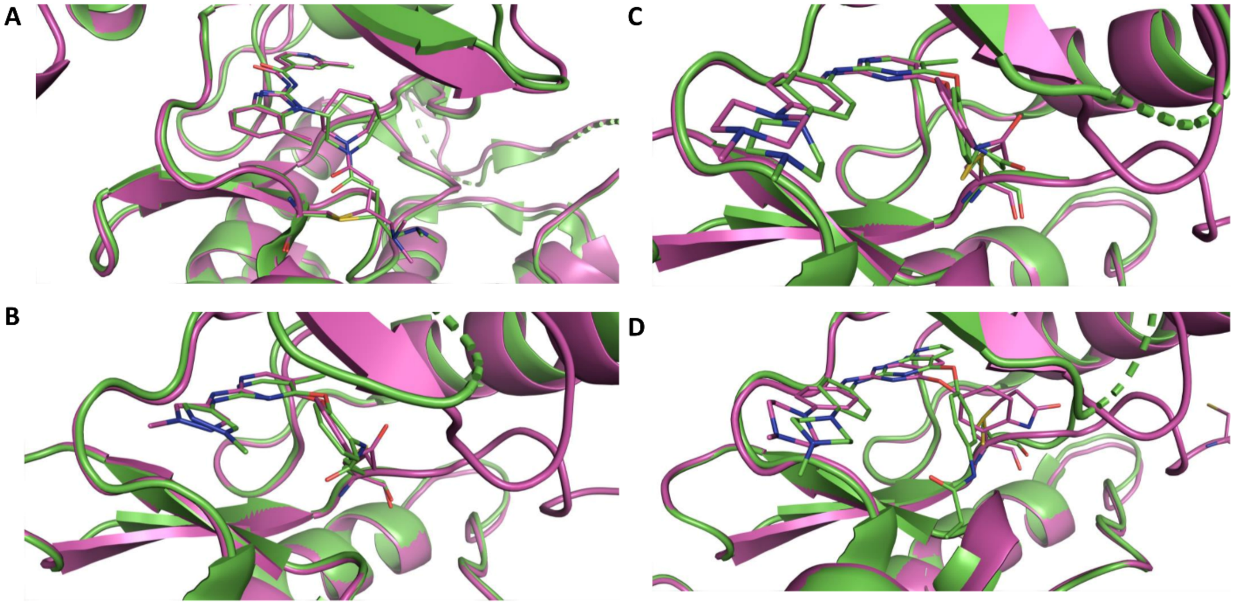


**Figure S3. Alignment of COValid structures with experimental PDB structures.** The release date of all plotted structures is before the AF3 training cutoff date. Ground-truth experimental structures are plotted in green, AF3 predictions are plotted in magenta. **A.** EGFR covalent binder (PDB ID: 5FEQ). Pocket-aligned RMSD: 0.67Å, mPAE: 1.1 **B.** MAP3K7 covalent binder (PDB ID: 5JK3). Pocket-aligned RMSD: 1.33Å, mPAE: 0.99. **C.** MAP3K7 covalent binder (PDB ID: 5J8I). Pocket-aligned RMSD: 0.84Å, mPAE: 0.96. **D.** Non-covalent MAP3K7 binder (PDB ID: 5J9L). Pocket-aligned RMSD: 3.63Å, mPAE: 1.8. This is an example of AF3 catching a mistake in COValid - while the ligand binds the pocket, it does not do so through a covalent binding mechanism. AF3 'identified' this mistake and resulted in a bad mPAE score. Structural figures were rendered via Pymol (*45*).

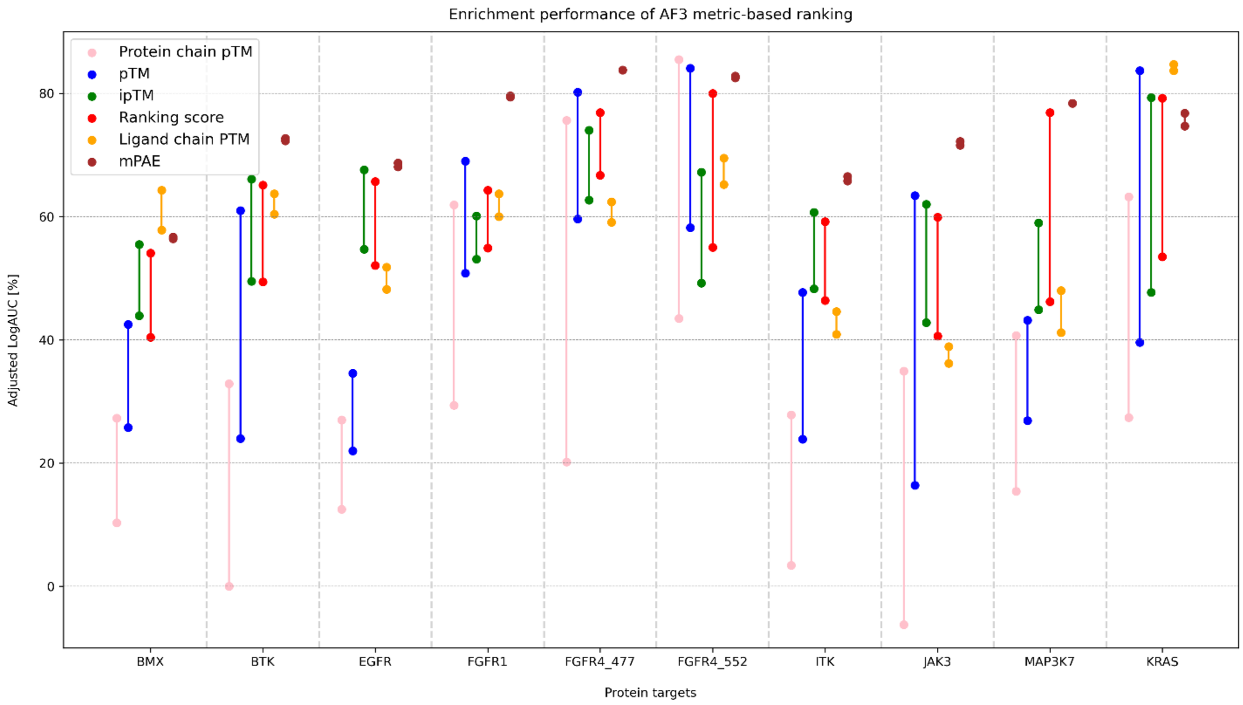


**Figure S4. Enrichment performance of metric-based ranking.** For each Cys site, the Adjusted LogAUC is reported as a range between the best-case scenario (actives are ranked higher than decoys that share the same metric value) and the worst-case scenario (decoys are ranked higher than decoys that share the same metric value).

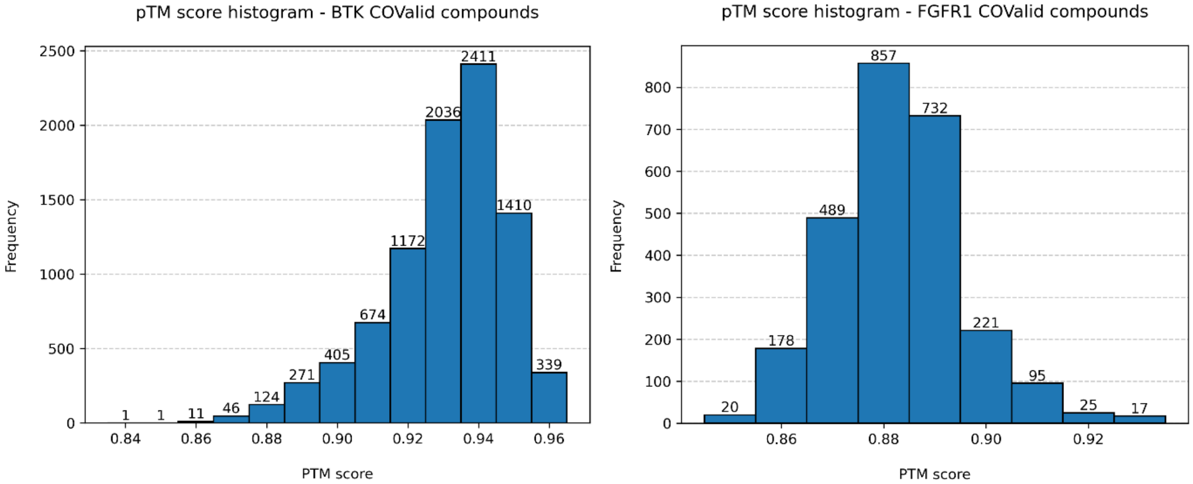


**Figure S5. pTM metric does not yield sufficient resolution for library ranking**. The histograms depict the count of pTM scores predicted for all BTK (left) and FGFR1 (right). 2411 BTK protomers and 857 FGFR1 protomers are assigned the same exact pTM value (0.94 and 0.88, respectively).

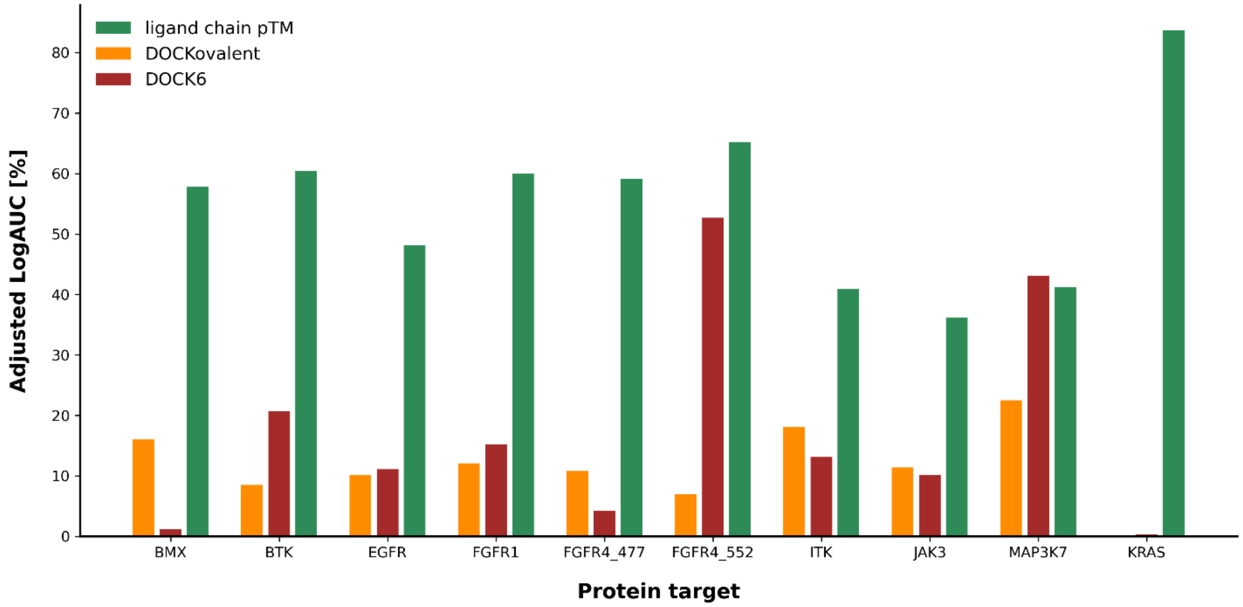


**Figure S6. Ligand chain pTM ranking outperforms docking tools for nine targets.** Plotted are the adjusted LogAUC values for the ten Cys sites of COValid, based on the docking score output of the two physics-based docking tools with leading enrichment performance (docking to the covalent PDB complexes of COValid), compared to AF3 predictions ranked by the ligand chain pTM metric (worst-case scoring).

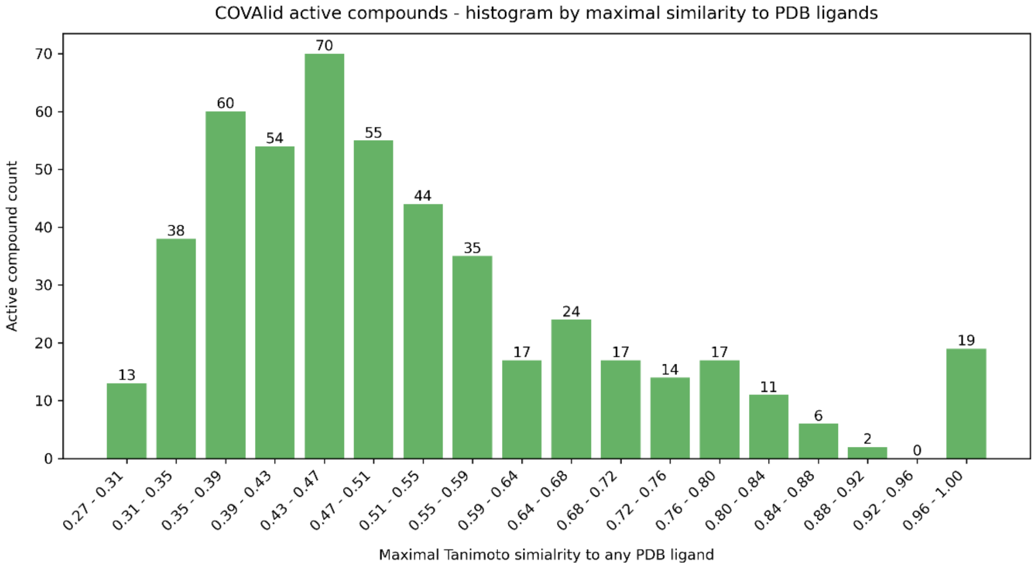


**Figure S7. COValid active compounds similarity to PDB ligands.** The histogram counts the COValid active compounds (prior to their protonation) based on the maximal Tanimoto similarity between their Morgan topological fingerprint and the fingerprints of ligands from the PDB, aggregated across the 10 COValid Cys sites.

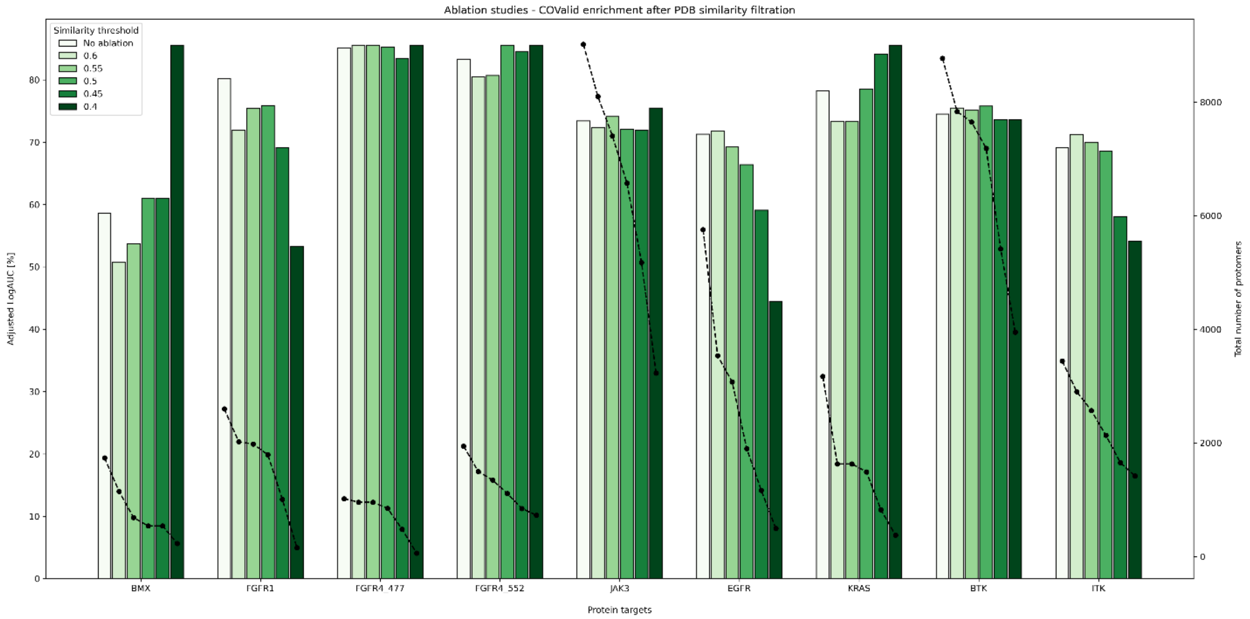


**Figure S8. COValid-PDB 'ablation' studies.** For each COValid Cys site, we plot the enrichment values if we only retain actives below different Tanimoto similarity to PDB ligands. The default dataset refers to the worst-case ranking based on mPAE. Given a cutoff level, any COValid active compound with a maximal Tanimoto similarity higher than the threshold to any PDB ligand is discarded. Specifically, the best-scoring active protomer of each discarded active is removed from the ranked compound list, and so are the 50 decoy protomers of each of the active protomers of the active compound. The columns depict the calculated adjusted LogAUC value (left vertical axis), and the black data points depict the total number of protomers in the ranked list (right vertical axis). MAP3K7 was not analyzed, since all its active protomers bear high similarity to ligands in the PDB.

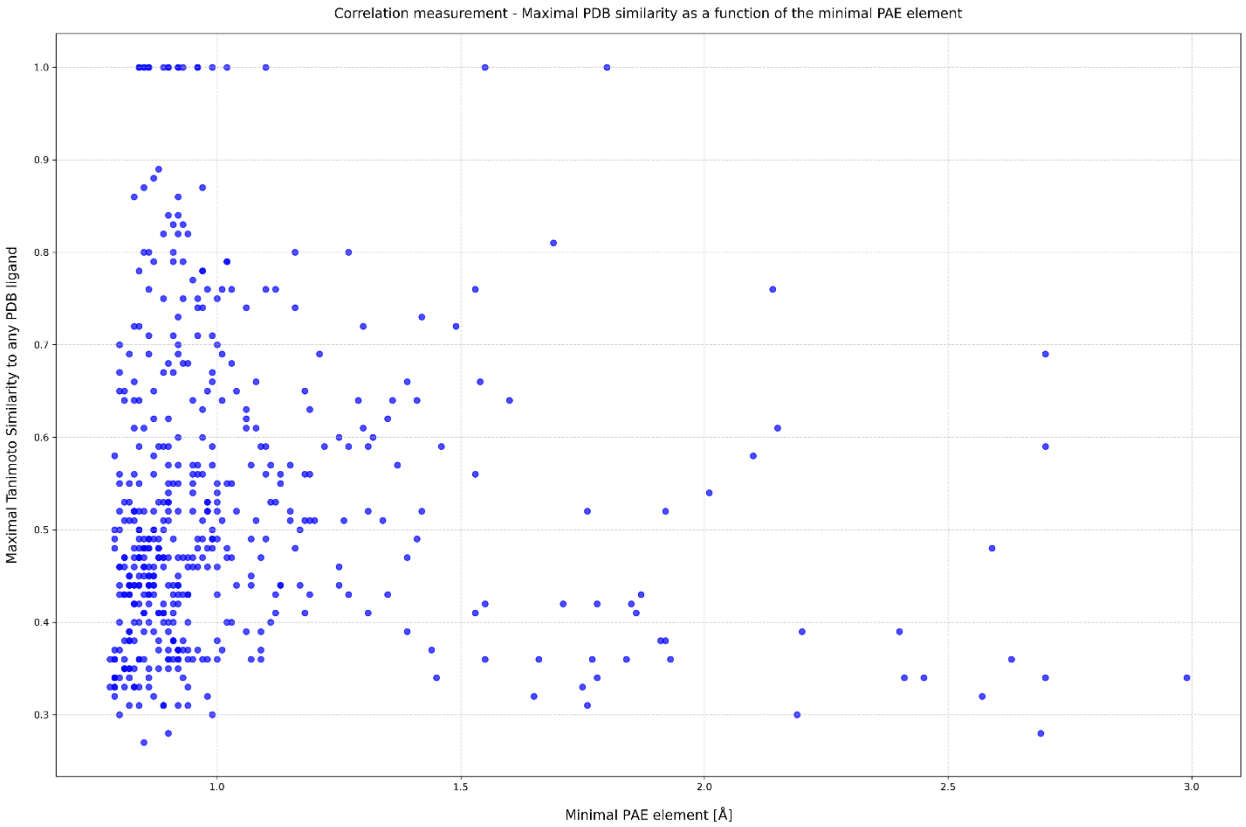


**Figure S9. No correlation between mPAE and the maximal Tanimoto similarity to PDB ligands.** We plot for the 496 active compounds in COValid the mPAE value (of the best-scoring protomer for each active compound) *vs.* the maximal TC to any PDB ligand (not necessarily with the same protein, and not restricted to AF3 training set). The Pearson correlation coefficient is -0.056, *p-value* is 0.21.

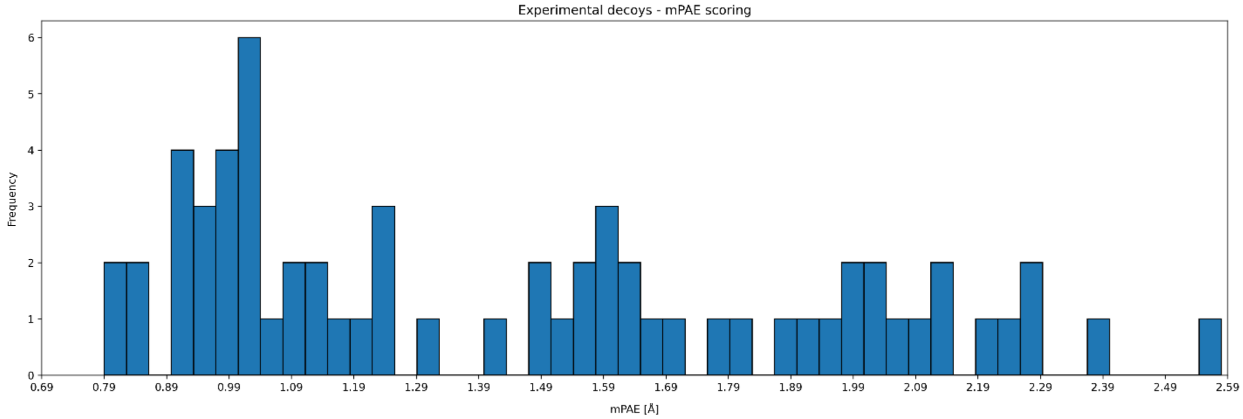


**Figure S10. mPAE values for experimental decoys.** 64 experimental decoys with $IC_{50}$ annotation were curated from ChEMBL, matching the curation process described for COValid, but with the enforcement of a lower limit on activity - 10[µM]. They are associated with six of the COValid targets - BTK(12), JAK3(5), EGFR(24), ITK(12), KRAS(10), and FGFR1(1). The average mPAE is 1.5±0.5Å.

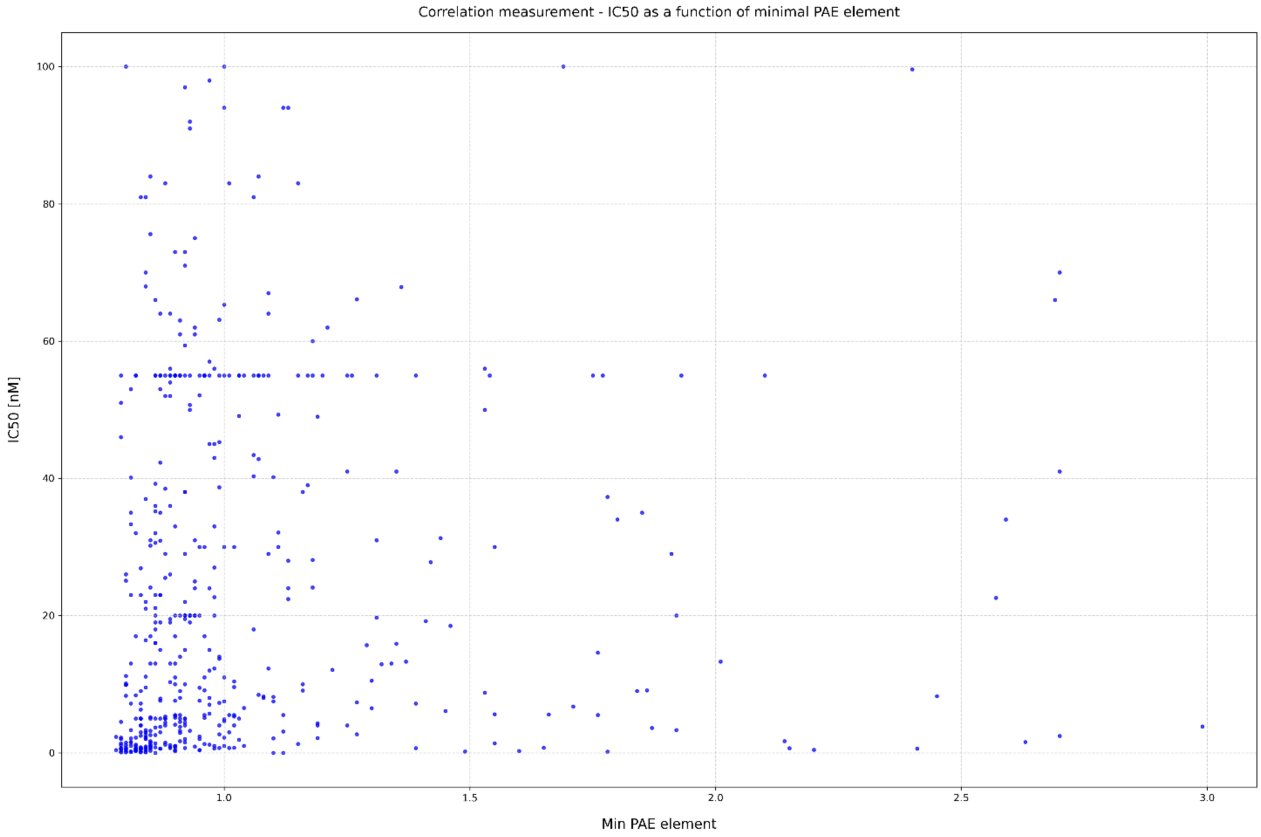


**Figure S11. mPAE does not correlate with $IC_{50}$ values.** The scatter plot depicts all COVAlid active compounds annotated with $IC_{50}$ values (481 out of the 496 active compounds), plotted against the minimal PAE element. Pearson correlation coefficient - 0.096, *p-value* - 0.035.

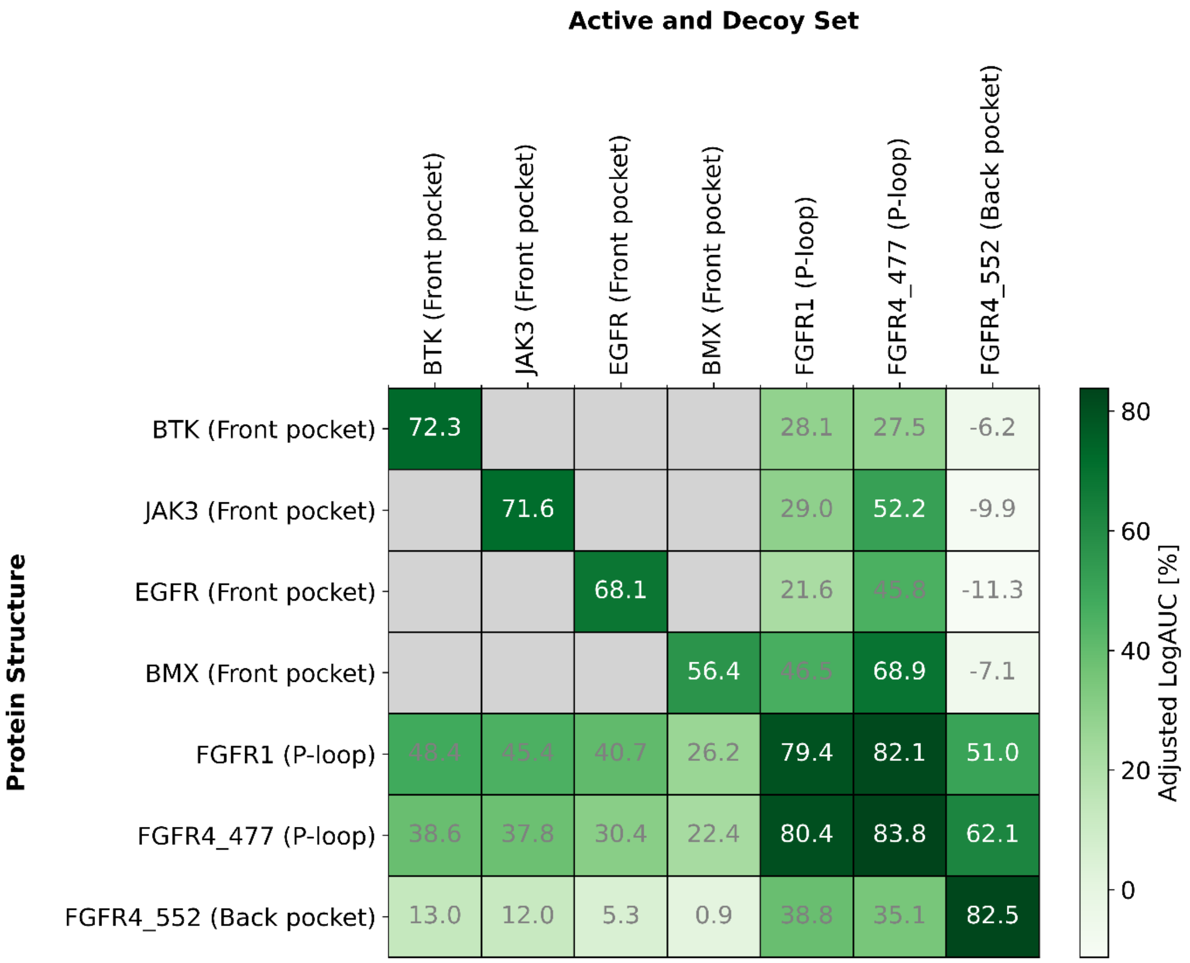


**Figure S12. AF3 cross-docking tests.** Y axis lists the COValid Cys site to which the compounds were covalently attached and the X axis lists the sets of COVAlid actives and decoys whose complexes were predicted. The location of the Cys within the kinase is depicted in parenthesis. Grey boxes indicate combinations that were not tested. Adjusted LogAUC values are presented according to mPAE-ranking (worst-case ranking). The highest cross-docking enrichment was measured when predicting a compound library associated with one P-loop Cys site in complex with the a P-loop Cys of a different protein (FGFR4 Cys477 and FGFR1 Cys488, and vice versa).

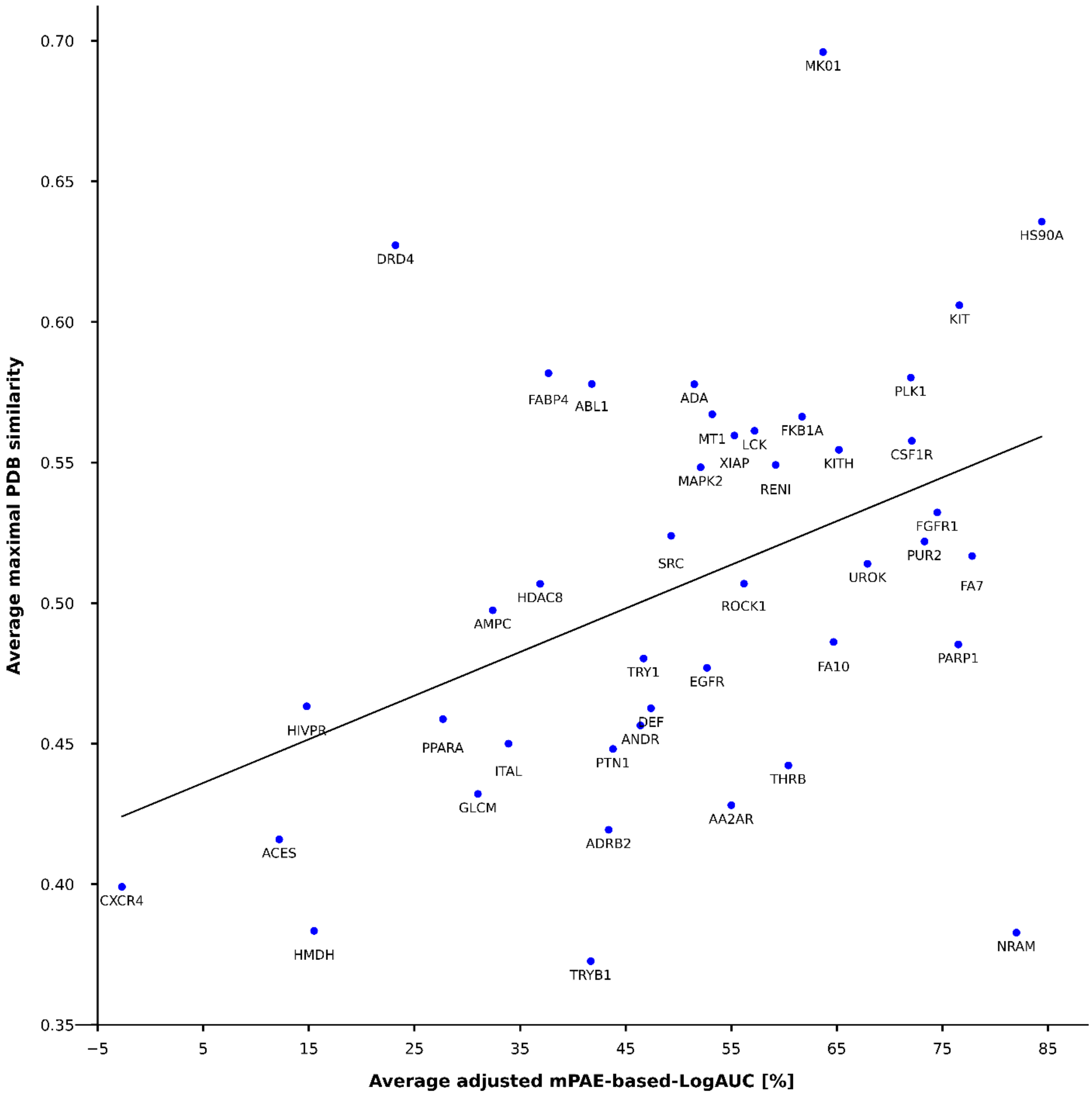


**Figure S13. Moderate correlation between average maximal PDB similarity and enrichment.** Across the 43 targets of DUDE-Z, a moderate correlation is observed between the average maximal similarity of the active compounds to any PDB compound and the average adjusted LogAUC values (Pearson correlation coefficient - 0.425, p-value - 0.00451, R-squared - 0.18).

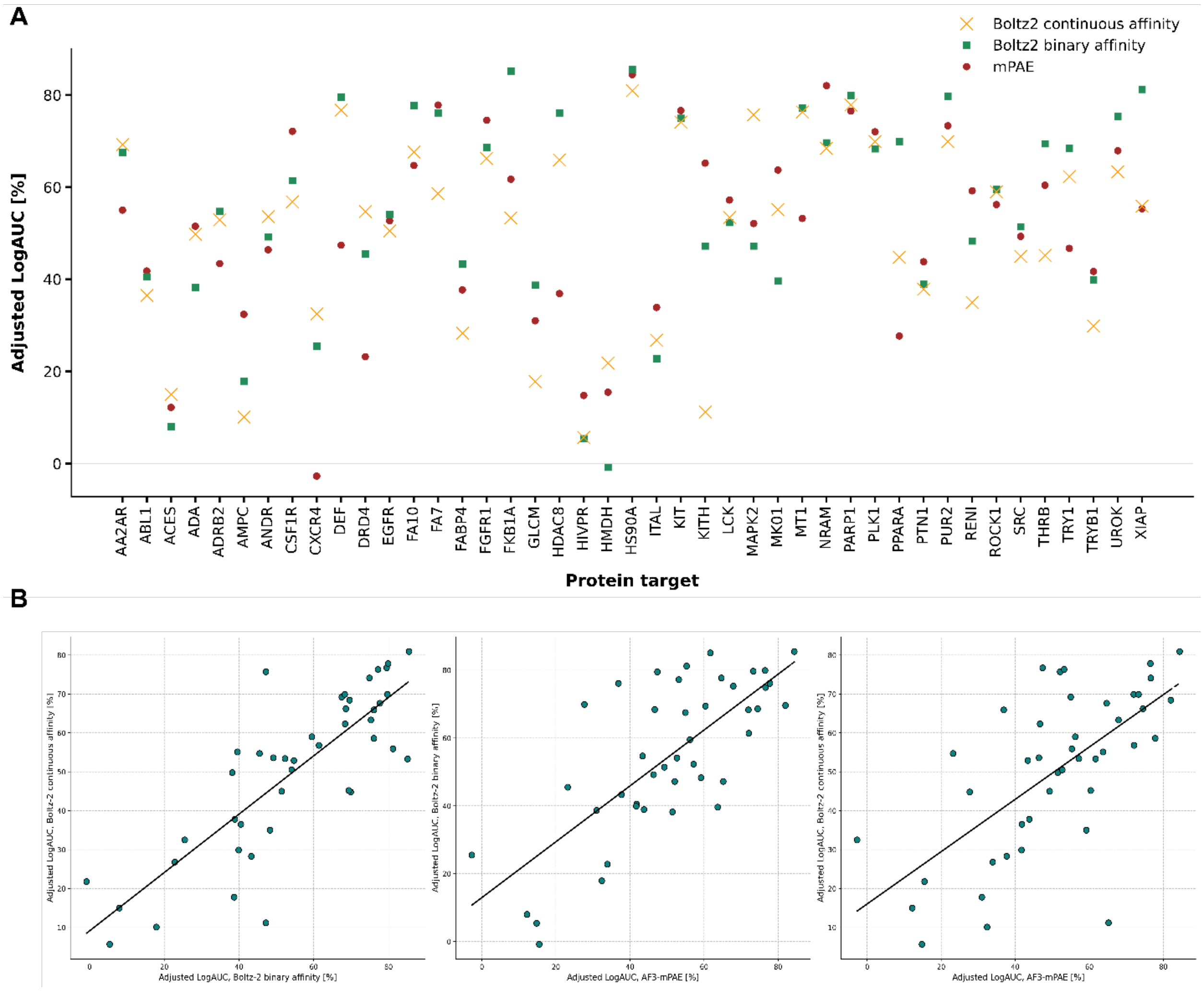


**Figure S14. Enrichment comparison - Boltz-2 and AF3 with mPAE. A.** Across the 43 targets of DUDE-Z, depicted are the enrichment values yielded when sorting the compound libraries by AF3 mPAE scores, Boltz-2 binary affinity scores, and Boltz-2 continuous affinity scores. **B.** Correlation measurements between set of enrichment values across DUDE-Z (Boltz-2 binary and continuous affinity - Pearson correlation coefficient - 0.82, p-value - 1.3E-11; AF3-mPAE and Boltz-2 binary affinity - Pearson - 0.74, p-value - 1.6E-8; AF3-mPAE and Boltz-2 continuous affinity - Pearson - 0.66, p-value = 1.4E

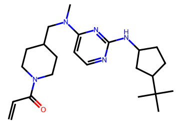

Z9597271729

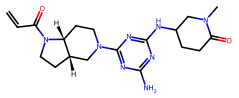

Z9597271738

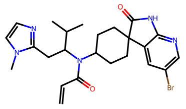

Z9597271743

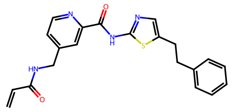

Z9597271754

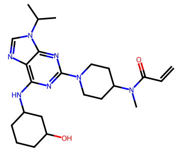

Z9597271720

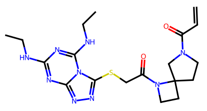

Z9597271750

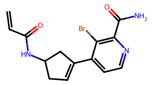

Z9597271713

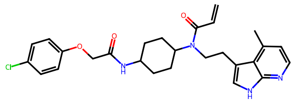

YS1

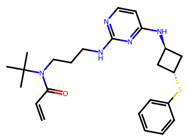

Z9597271739

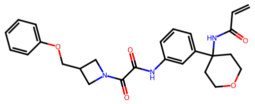

YS2

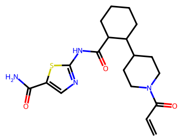

Z9597271714

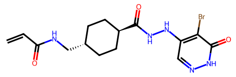

Z9597271752

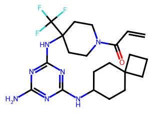

YS3

**Figure S15. Chemical structures of compounds synthesized for experimental validation.**

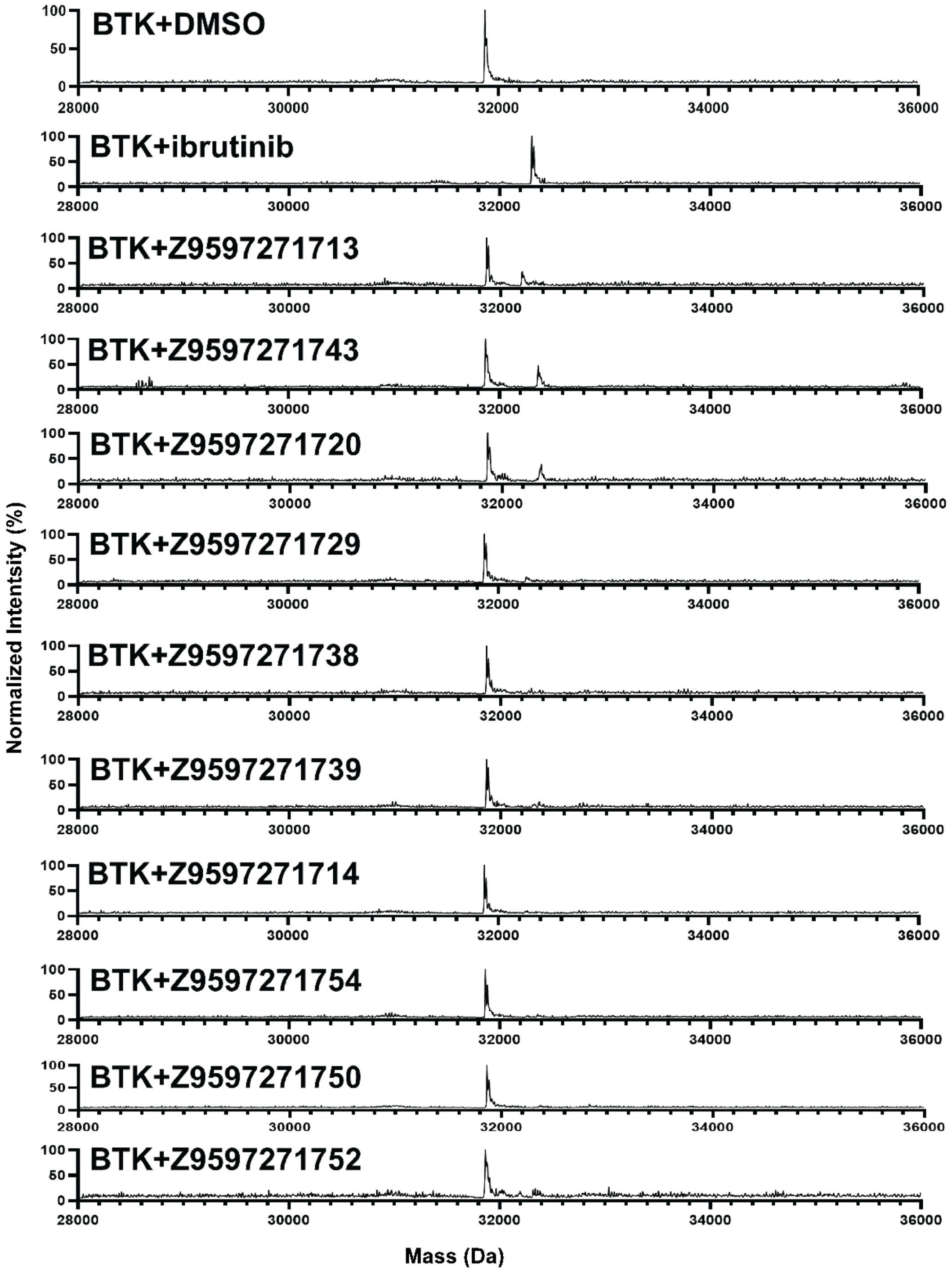


**Figure S16. Intact protein LC/MS spectra for synthesized compounds.** Deconvoluted spectra depicting protein labeling by the compounds (1 μM BTK, 200 μM compound, pH 7.5,RT, 2h). Data shown for compounds that did not reach near-100% labelling.

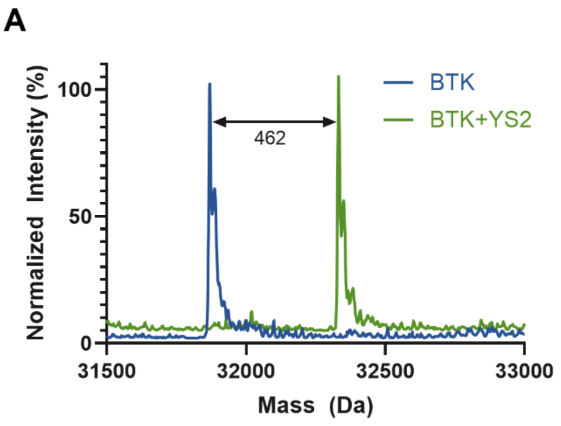


**Figure S17. Intact protein LC/MS spectra for main hit compounds.** Deconvoluted spectra depicting protein labeling by compound YS2 (A) and YS3 (B) (1 μM BTK, 200 μM compound, pH 7.5, RT, 2h).

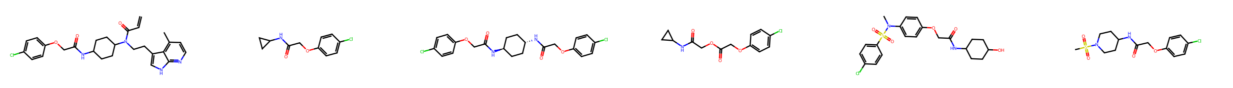

YS1 | Tanimoto Sim. = 0.46, CHEMBL1710054 | Tanimoto Sim. = 0.46, CHEMBL4303573 | Tanimoto Sim. = 0.41, CHEMBL1900394 | Tanimoto Sim. = 0.41, CHEMBL1895838 | Tanimoto Sim. = 0.4, CHEMBL3492950

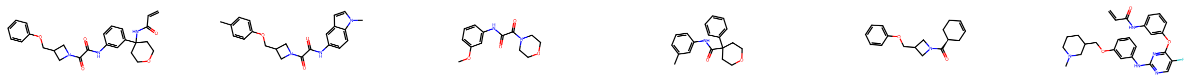

YS2 | Tanimoto Sim. = 0.41, CHEMBL4875161 | Tanimoto Sim. = 0.4, CHEMBL1564952 | Tanimoto Sim. = 0.36, CHEMBL1887133 | Tanimoto Sim. = 0.36, CHEMBL5005354 | Tanimoto Sim. = 0.35, CHEMBL6044019

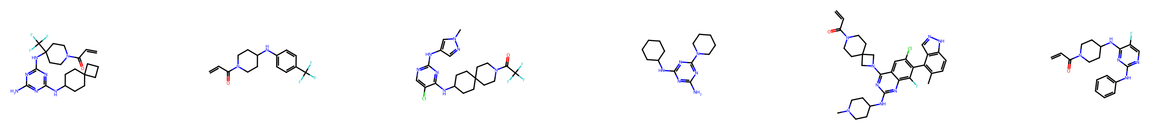

YS3 | Tanimoto Sim. = 0.36, CHEMBL5884138 | Tanimoto Sim. = 0.34, CHEMBL3953702 | Tanimoto Sim. = 0.33, CHEMBL125250 | Tanimoto Sim. = 0.33, CHEMBL5170321 | Tanimoto Sim. = 0.33, CHEMBL5835925

**Figure S18. most similar ChEMBL compounds to the main hits.** For each main hit from the BTK prospective screening, the top-5 most similar compounds from the entire ChEMBL36 dataset are plotted (Morgan fingerprints, 2048 bit, radius = 2). Similarity analysis was conducted via FPSim2.

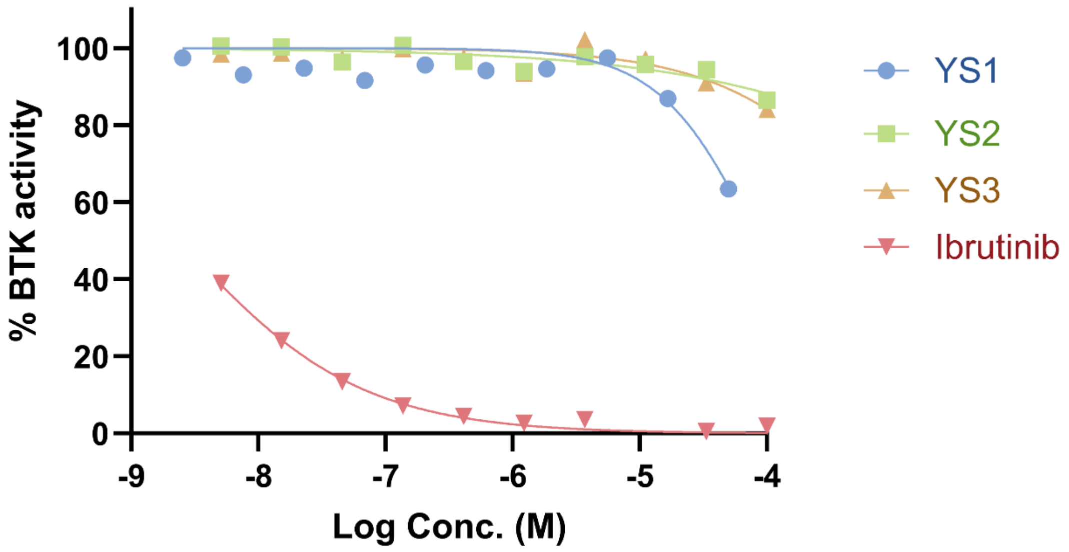


**Figure S19. In vitro kinase inhibition assay against BTK C481S.** Kinase inhibition analysis by a radiometric filter binding assay measuring phosphorylated substrates products with BTK mutant C481S (20 μM ATP, 3 nM BTK, 0.2 mg/ml pEY substrate, pH 7.5, RT).

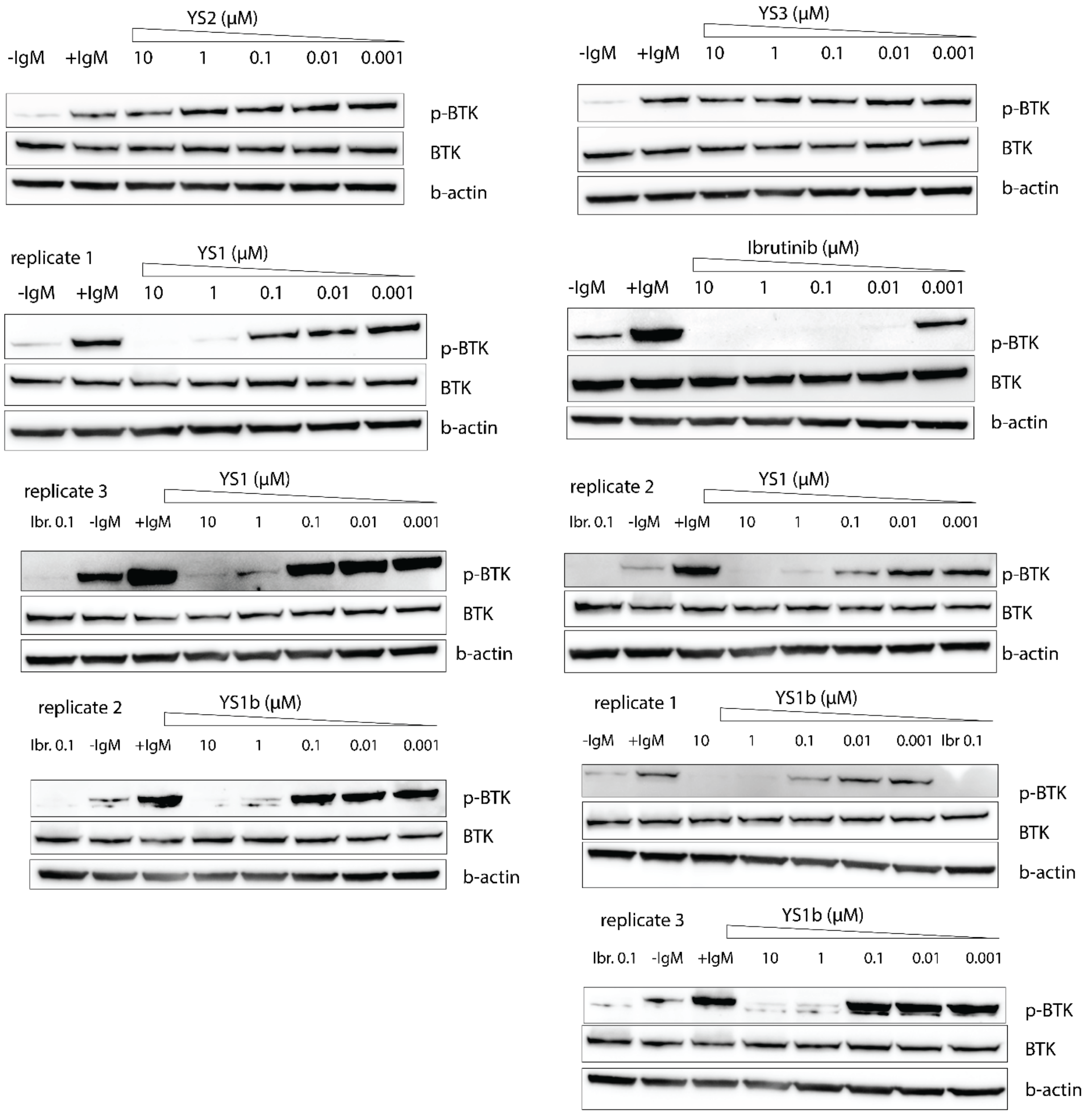


**Figure S20. Cellular pBTK inhibition assays.** Western plots depicting dose-response inhibition of BTK phosphorylation by YS1 and YS1b (n = 3), as well as YS2 and YS3 (n = 1). For quantification of YS1 activity see Fig. 3J.

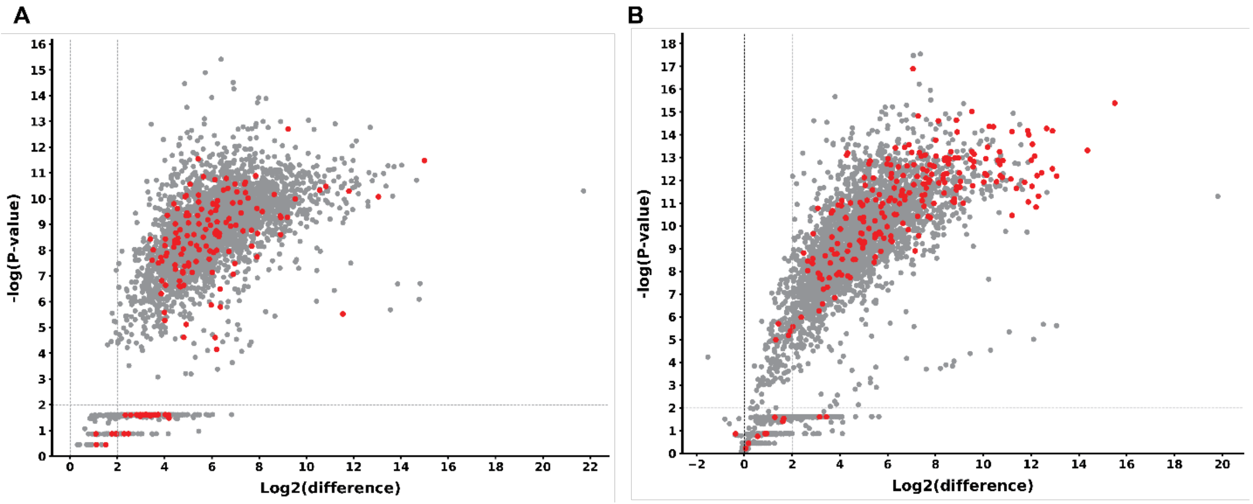


**Figure S21. Quantification of pulled-down proteins from proteomic experiments.** Mino cells were treated with 1 μM of DMSO for 1h followed by 45 min treatment with either 1 **μ**M Probe 4 or 2 **μ**M XO44 (Probe 4 in A, XO44 in B; n = 4). Proteins were quantified using label-free quantification. Proteins in the upper right segment show a significant change (fold change > 2; p < 0.01). In the Probe 4 experiment, 3,415 proteins are plotted (159 kinases, depicted in red, and 3,256 non-kinases), out of which 2,984 proteins exhibit statistical significance (131 kinases and 2,853 non-kinases). In the XO44 experiments, 3,556 proteins are plotted (235 kinases and 3,321 non-kinases), out of which 2,945 proteins exhibit statistical significance (220 kinases and 2,725 non-kinases). Kinases were determined by a list of human kinase Uniprot entries from KinHub (*46*).

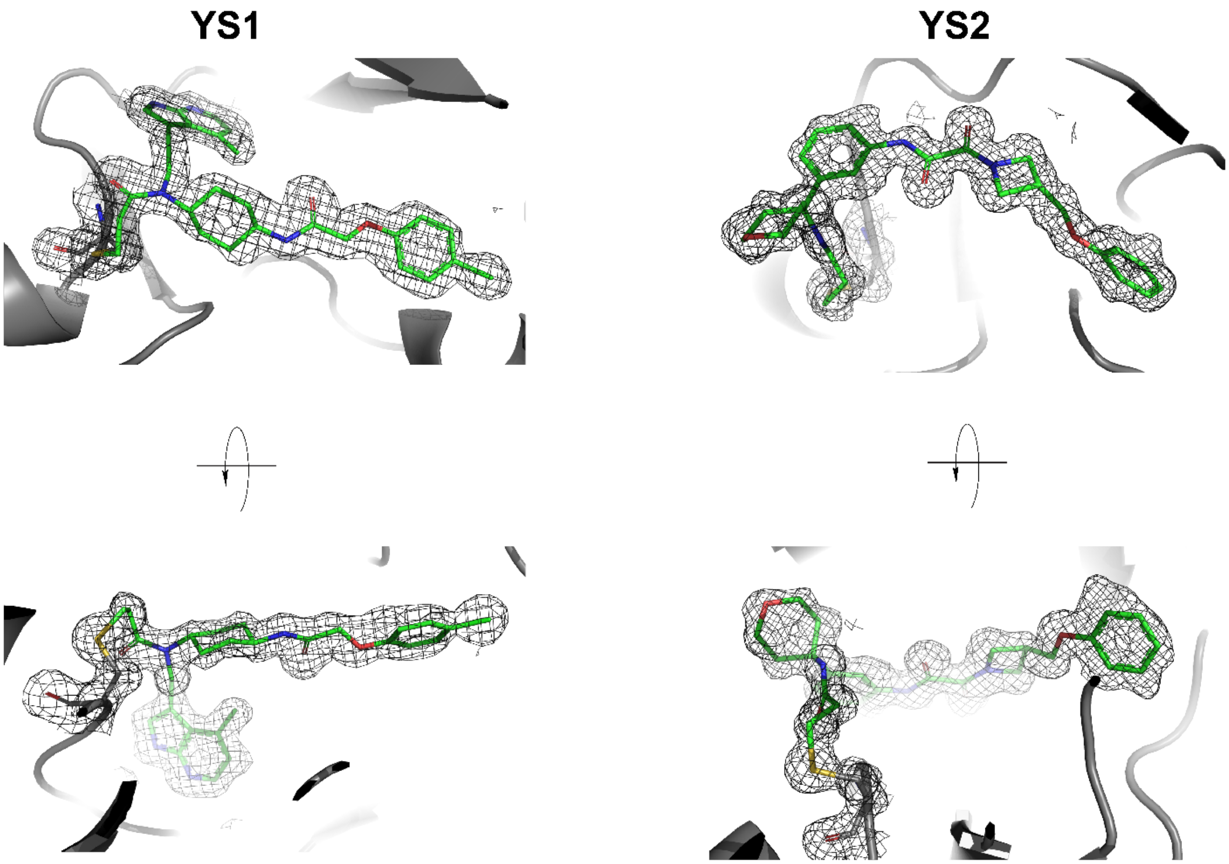


**Figure S22. YS1 and YS2 ligand electron density.** Electron density maps clearly define the YS1 (left) and YS2 (right) ligands in the BTK active site. Two views of each are shown.

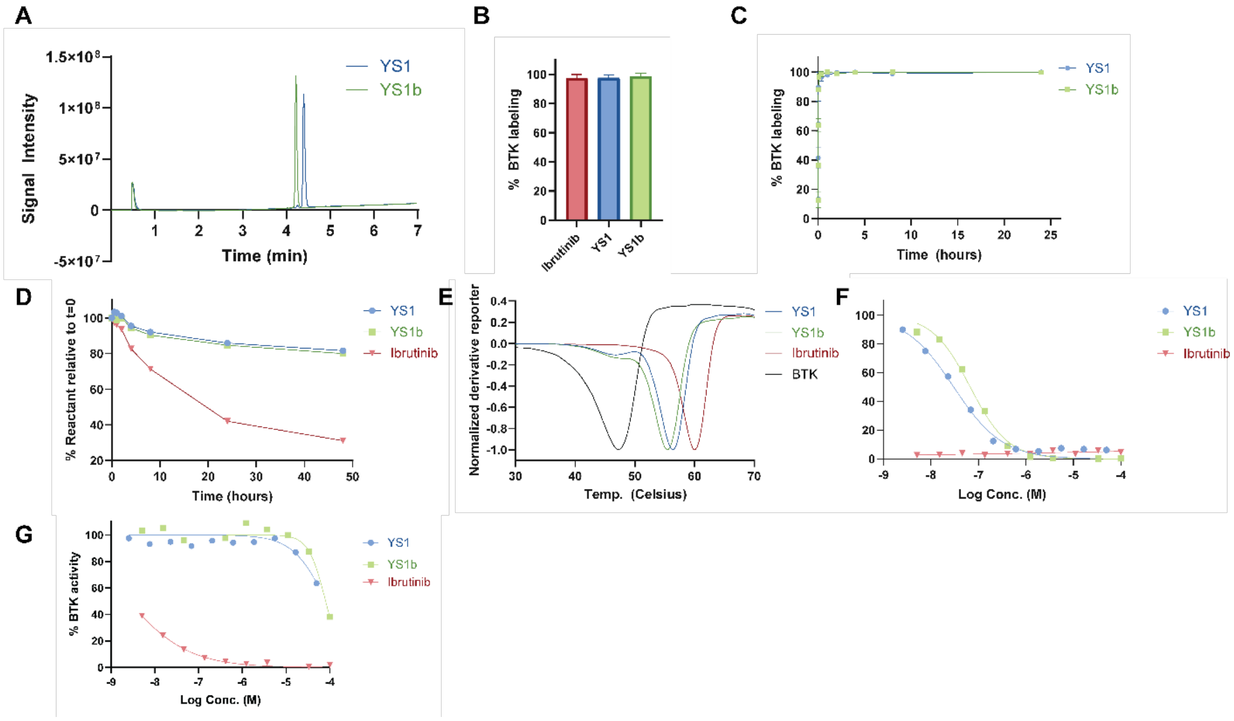


**Figure S23. Biochemical comparison of YS1 isomers.** The cyclohexyl moiety in YS1 may be found in either a cys- or trans- isomer. We were able to separate two species of YS1, presumably corresponding to these isomers. The crystal structure of YS1 with BTK (Fig. 4A) unequivocally identifies YS1 as the trans-isomer (as predicted by AF3). We name the second species (supposedly the cys- isomer) as YS1b. **A.** UPLC retention times for the two YS1 isomers. **B.** Intact protein LCMS covalent labeling percentage (1 μM BTK, 200 μM compound, pH 7.5, RT, 2h). n = 2, data represented as mean with SD as error bars. **C.** LC/MS time-course experiment (1 μM BTK, 5 μM compound, RT, pH 7.5). n = 2, data represented as mean with SD as error bars. **D.** Reduced glutathione (GSH) assay for reactivity assessment (5 mM GSH, 0.2 mM compound, pH 8, 25℃). **E.** Differential scanning fluorimetry (DSF) analyzes the shift in BTK melting point after treatment with the compounds (5 μM BTK kinase domain, 0.02 mM compound, pH 7.5, overnight incubation at 25°C). The derivative reporter, normalized by the absolute value of the minimum in each dataset, was used to determine the melting point. Ibrutinib, YS1, YS2 and YS3 stabilize BTK by ~13°C, 9°C, 5°C, and 1°C, respectively. BTK baseline curve is plotted in black. **F** Kinase inhibition analysis by a radiometric filter binding assay measuring phosphorylated substrates products (20 μM ATP, 3 nM BTK, 0.2 mg/ml pEY substrate, pH 7.5, RT). IC50 of YS1 and YS1b is 30 nM and X nM, respectively. **G.** Kinase inhibition analysis with C481S BTK mutant.

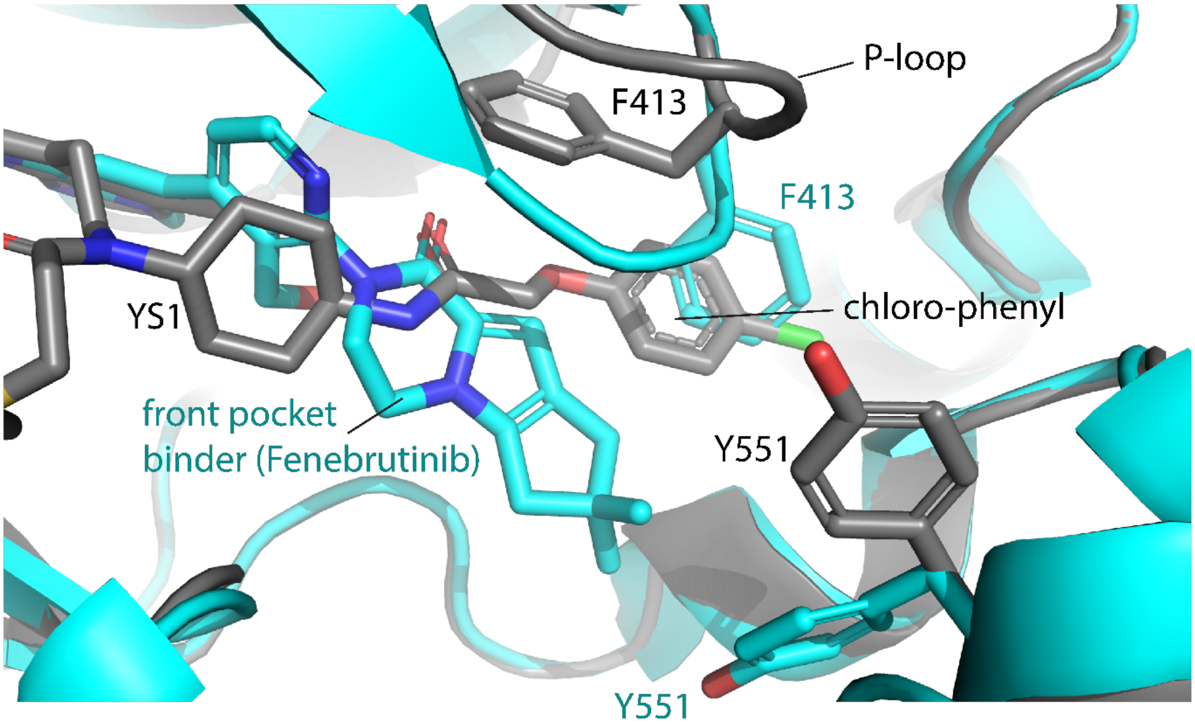


**Figure S24. Comparison of YS1 and Fenebrutinib binding poses.** Superposition of the YS1/BTK structure (gray) with a typical front pocket binder bound to the BTK kinase domain (Fenebrutinib; cyan). YS1 (shown in gray) binds into a pocket that is distinct from the well-characterized back- or front-pockets (see details in Fig 4A). The P-loop configuration is distinct in the YS1/BTK structure, leading to a completely opposite orientation of the Phe413 side chain. This likely results from the fact that the chloro-phenyl group of YS1 occupies the space normally filled by the sidechain of Phe413 in structures of front-pocket binders. The conformation of the activation loop tyrosine (Y551) is distinct from the sequestered state induced by front-pocket binders. Fenebrutinib is used here as an example of a front-pocket binder (PDB: 5VFI). There are also two covalent inhibitors for which structures are solved that bind the front pocket (PBD: 6TFP and 5J87). None are topologically similar to YS1.

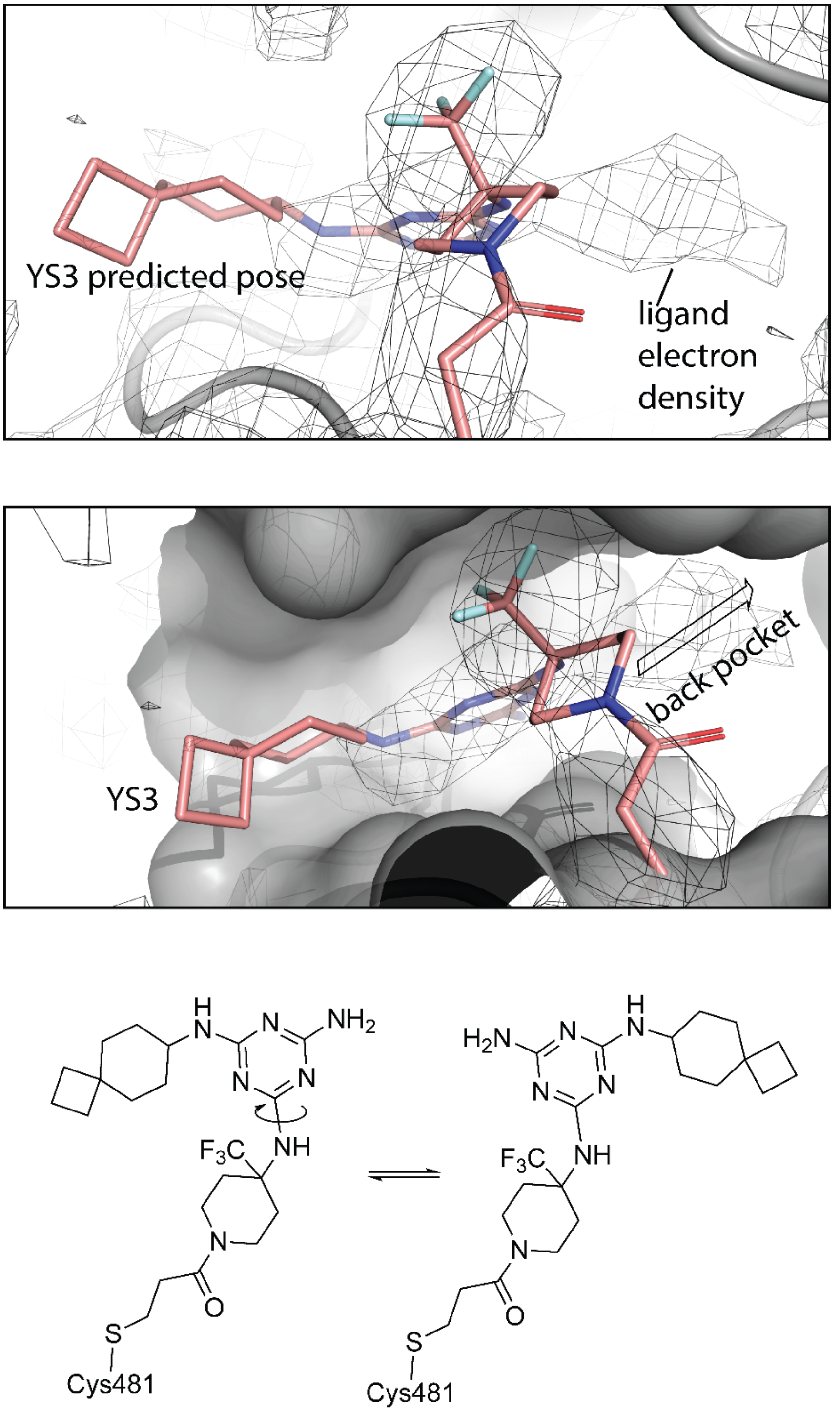


**Figure S25. Discrepancy in predicted and experimental YS3 structures.** Superposition of predicted YS3 complex with BTK kinase domain with electron density map derived from YS3 complex diffraction data. The predicted pose for YS3 is shown in salmon color, electron density as mesh. The YS3 ligand projects toward solvent in the predicted pose while the ligand electron density projects into the back pocket of the BTK active site. The two poses are related by rotation around a single bond (bottom).

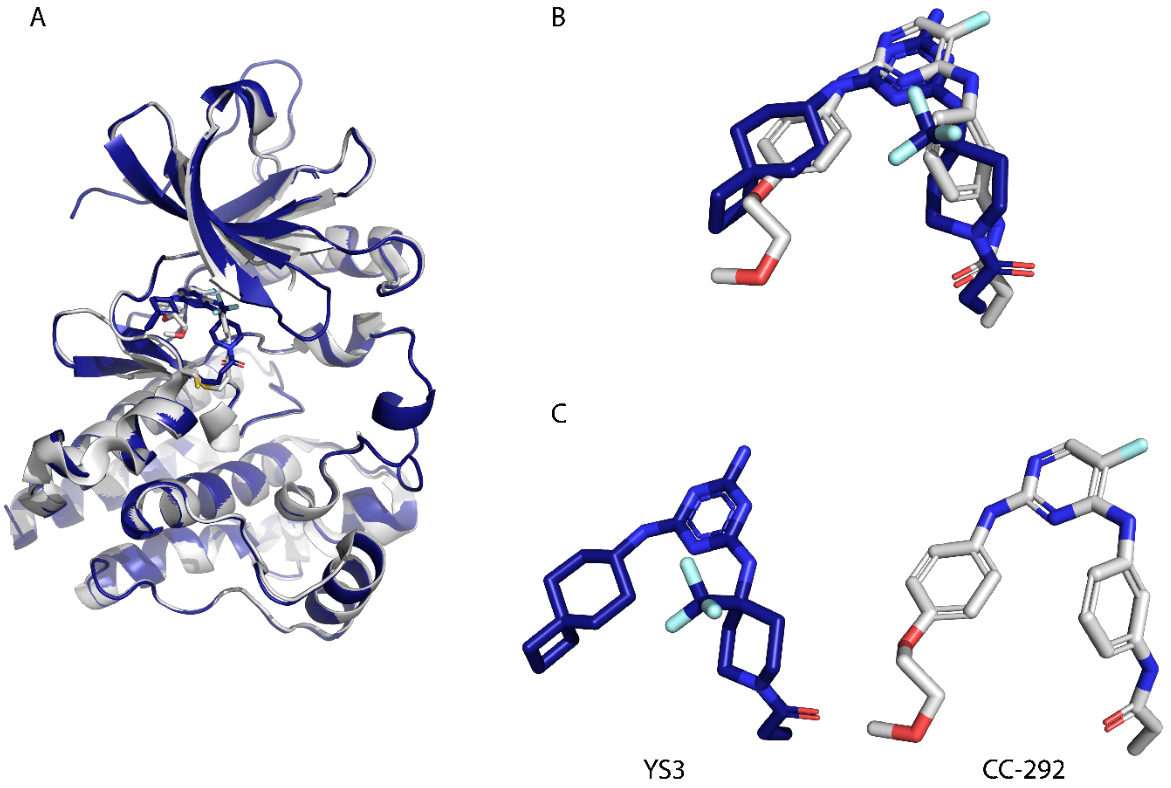


**Figure S26. Binding pose comparison of YS3 and CC-292.** (A,B) Overlay of predicted model of YS3 bound to the BTK kinase domain (blue) and crystal structure of CC-292 bound to the BTK kinase domain, PDB ID: 5P9L (gray). (C) Side by side view of bound conformations of YS3 model of CC-292 structure.

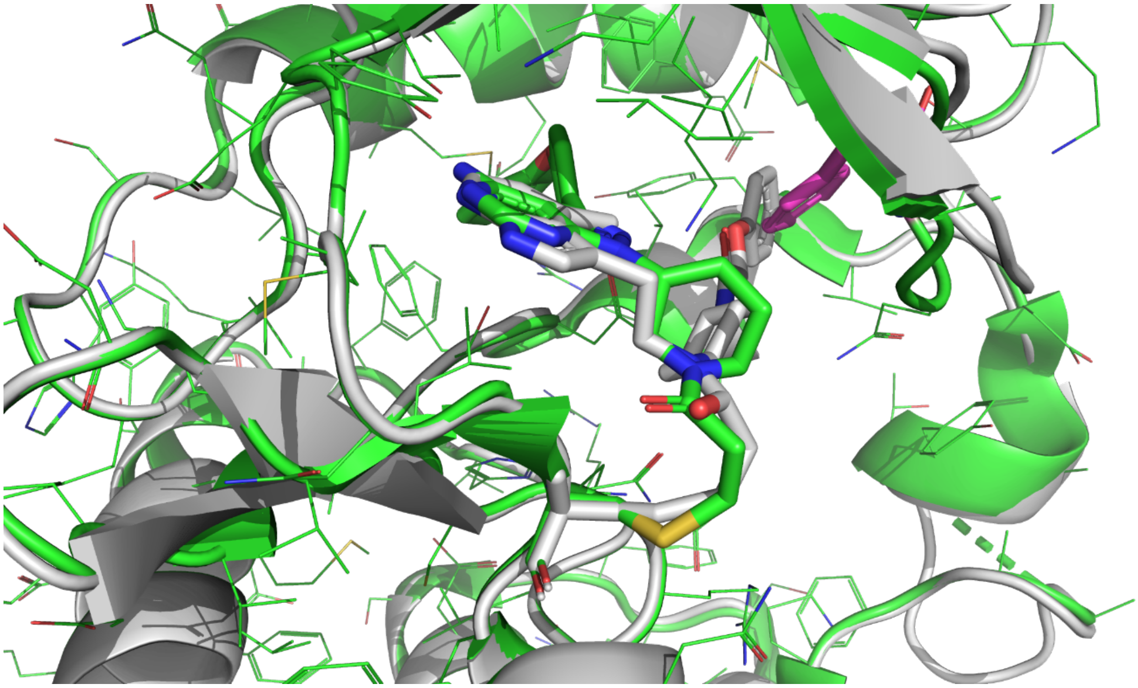

**Fig. S27. YS1 clashes with the Ibrutinib-bound BTK conformation.** Aligned structures of BTK-YS1 (grey) and BTK-Ibrutinib (green). The sidechain of Phe413 in the Ibrutinib complex is shown in magenta.

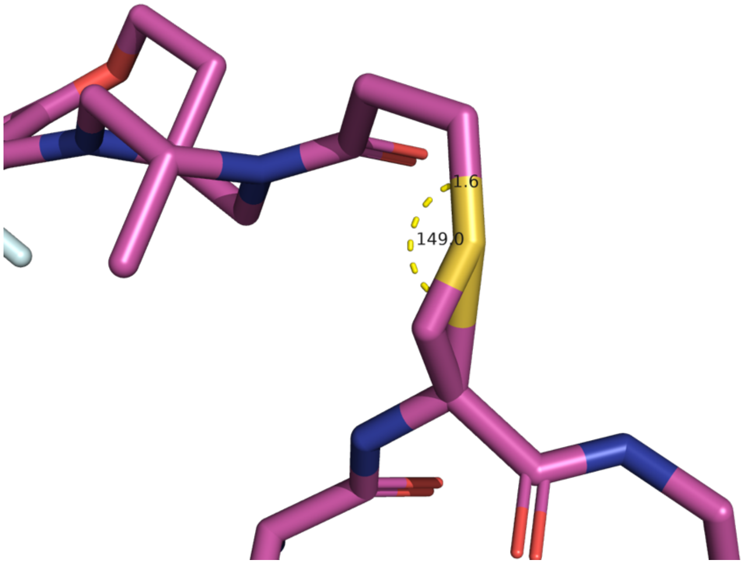


**Figure S28. AF3 produces malformed covalent bonds.** A characteristic example of an AF3 model with an unrealistically large bond angle of 149° and unrealistically short covalent bond length of 1.6Å.

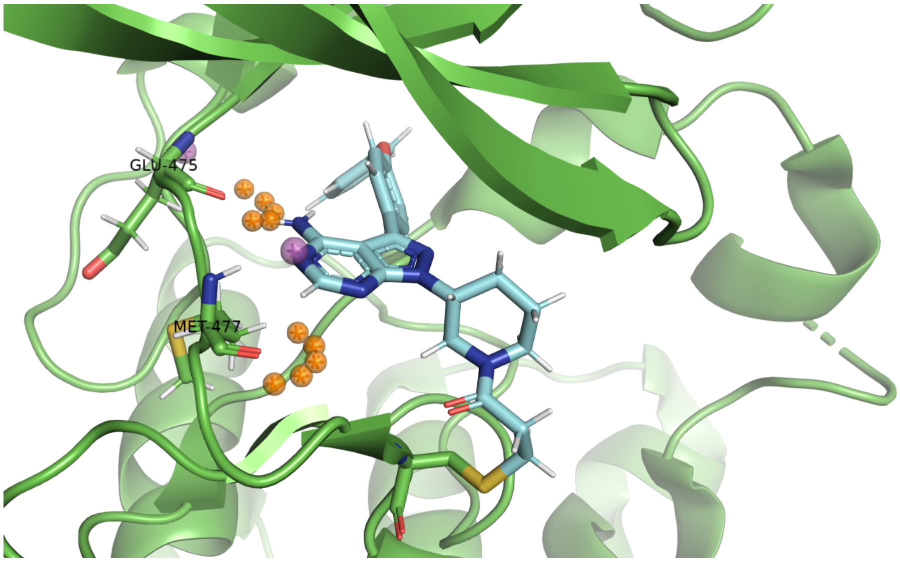


**Figure S29. AutoDock Bias - ideal interaction sites for hydrogen bond formation.** Implementation of AutoDock Bias illustrated with the covalent complex of ibrutinib with BTK (PDB ID: 5P9J). The Adenosine-mimetic substructure of the ligand forms canonical hinge interactions with two residues from the hinge region. Given the PDB structure as input, AutoDock Bias calculates ideal locations for placing ligand atoms that can form hydrogen bonds with the selected residues of the protein - for each hydrogen bond acceptor, five ideal locations for a hydrogen bond donor are calculated (in orange), for each hydrogen bond donor, a single ideal location for a hydrogen bond acceptor is calculated (in magenta). The output includes such sites according to the residue side chains as well, but we only used the sites calculated for the backbone amine and carbonyl.

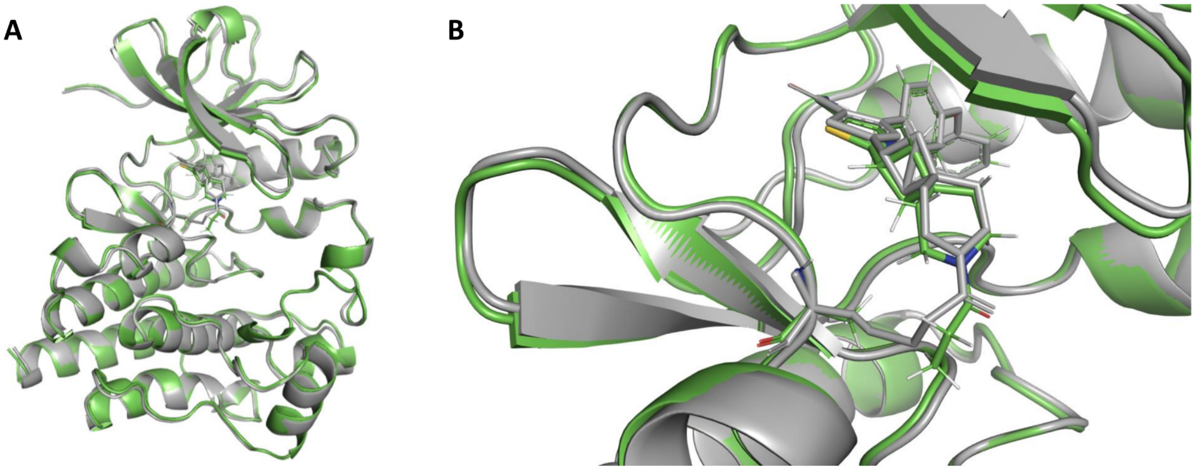


**Figure S30. Example of Rosetta minimization of AF3-predicted covalent complex**. **A.** The AF3 prediction output for one of the BTK COVValid compounds is aligned with the Rosetta-minimized structure (shown in grey and green, respectively). **B.** Close view of the ligand binding site. In the minimization process, we removed the Sγ atom of the Cys, and added hydrogen atoms to the ligand according to the protonation state of the COVAlid SMILES representation of the protomer.

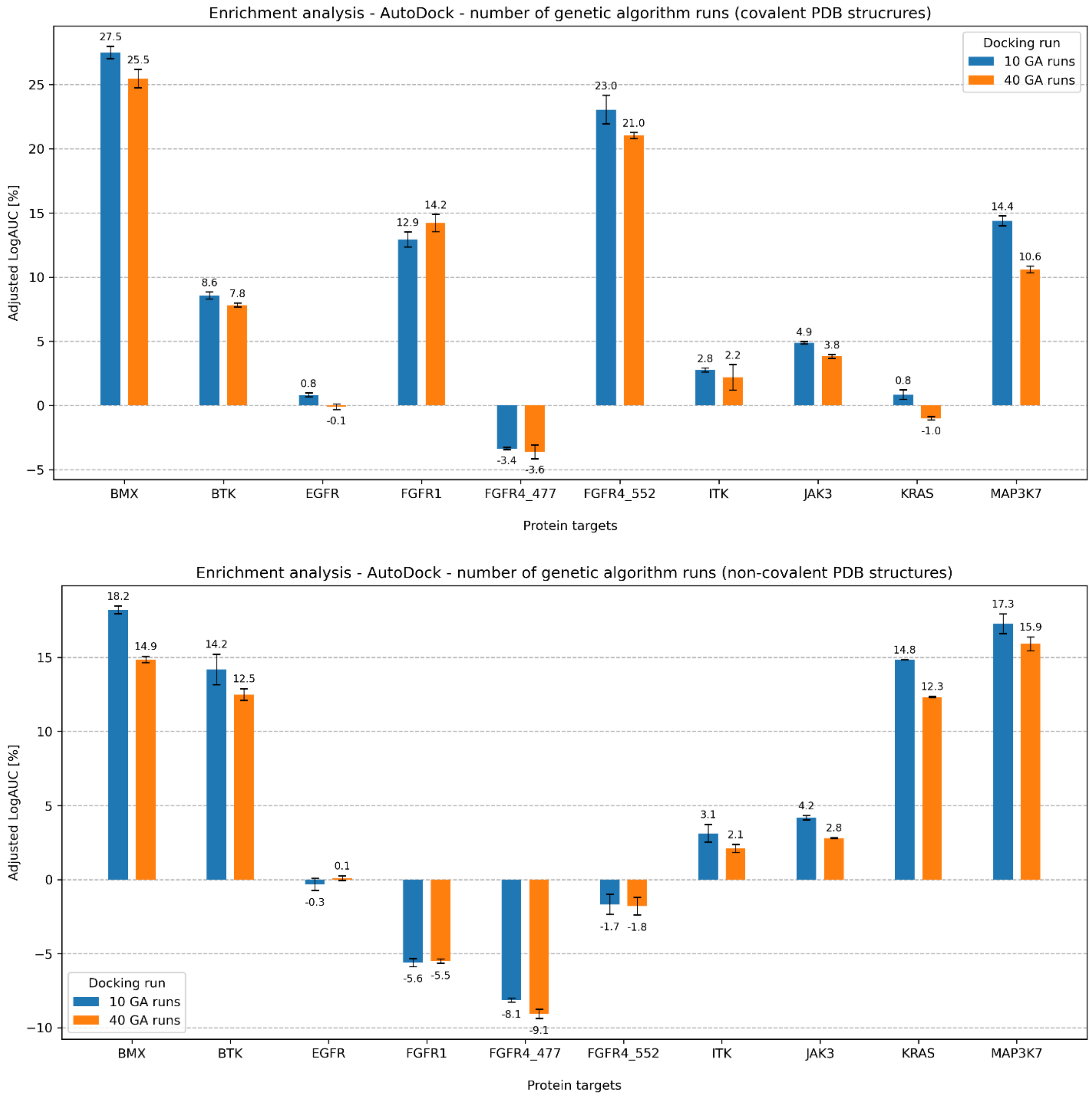


**Figure S31. AutoDock - effect of number of genetic algorithms runs on enrichment.** Due to the non-deterministic nature of genetic algorithms such as the one used by AutoDock, the algorithm is typically run multiple times, and the best-scoring output (as determined by the AutoDock score) is determined to be the binding pose prediction. In a comparison between values of 10 and 40, in spite of an approximate fourfold increase in the duration of each run, enrichment did not increase, as measured by the average adjusted LogAUC (6.6±7.4% and 5.5±6.9% for the runs with 10 and 40 GA runs, respectively).

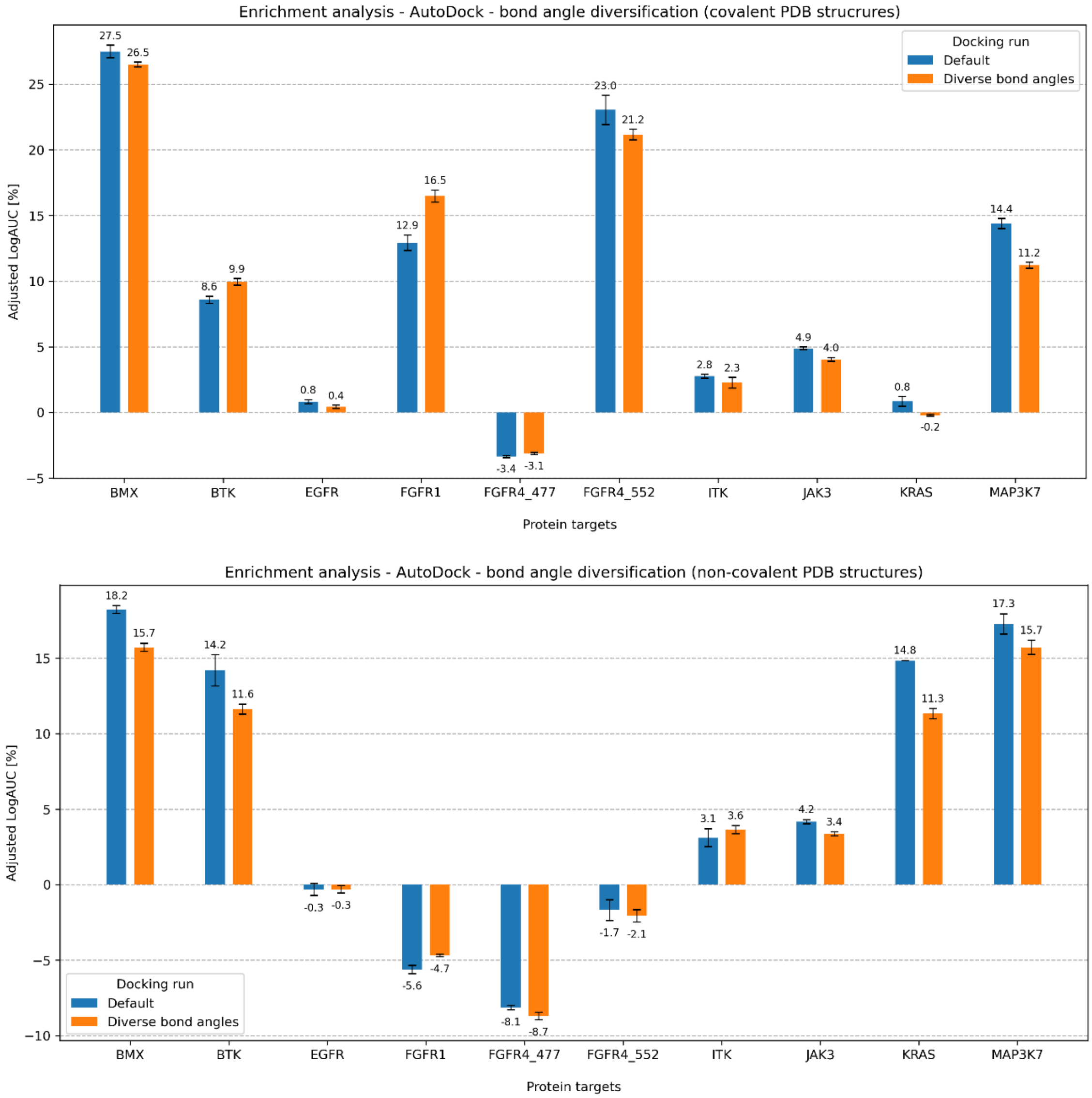


**Figure S32. AutoDock - effect of bond angle diversification on enrichment.** In the default settings of AutoDock4.2, only torsional degrees of freedom are sampled, meaning that bond angles are fixed to their value in the pre-generated input conformer. Attempting to probe the effect of bond angle sampling on enrichment, we introduced angle diversification to the ligand generation scheme. After generating an initial 3D conformer, we used it to generate four conformers by adding ±10° to two bond angles, one defined by the Cys Cβ, Cys Sγ, and the acrylamide Cβ atom, and another by the Cys Sγ, acrylamide Cβ, and acrylamide Cα atoms. We docked all four variants for each protomer, and used the best-scoring of the four in the enrichment analysis. In spite of a fourfold increase to the number of runs, the averaged adjusted LogAUC was slightly higher in the default scheme than in the angle diversification scheme (6.6±7.4% and 6.0±7.0%, respectively). Error bars reflect the standard deviation across three replicates of AutoDock. Fixed parameters: 10 GA runs.

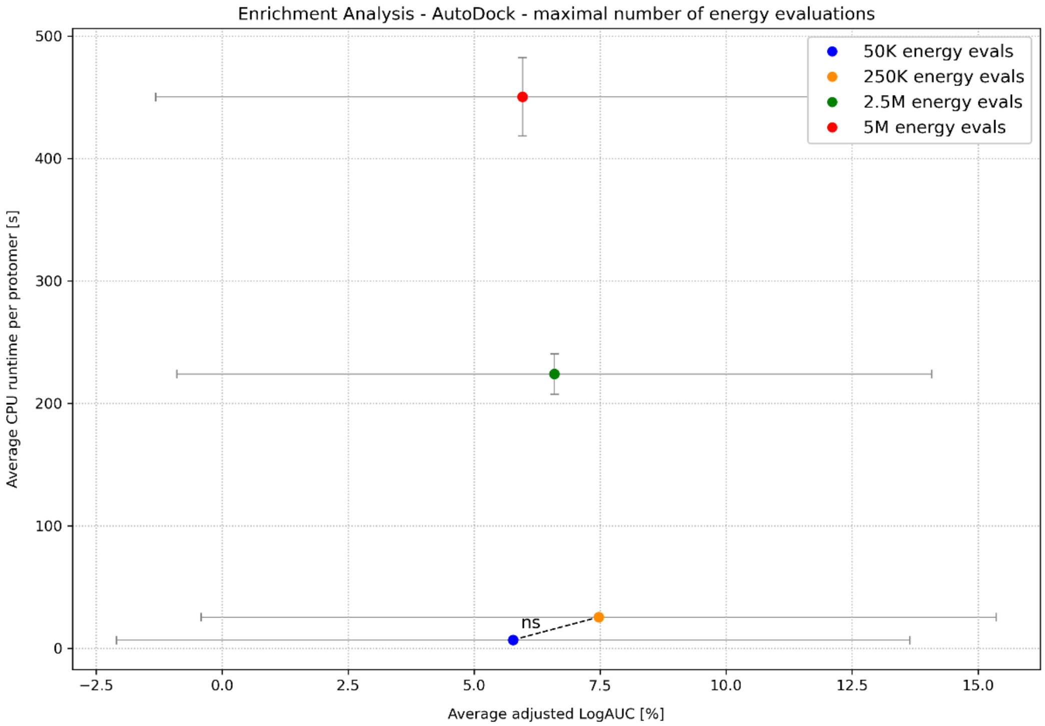


**Figure S33. AutoDock - effect of maximal number of energy evaluations on enrichment.** In genetic algorithms such as the one used by AutoDock, the maximal number of energy evaluations (named ga_num_evals in AutoDock) define the maximal number of energy evaluations conducted throughout an entire run of the docking algorithm, such that the algorithm halts if this maximal value is reached. Increasing the value from 50K to 250K yielded an insignificant increase to the average adjusted LogAUC (*p-value*: 0.0516). Interestingly, increasing the value further yielded no improvement to enrichment. By this analysis, using a low value can reduce runtime significantly without an effect on enrichment. Fixed parameters: 10 GA runs, no bond angle diversification.

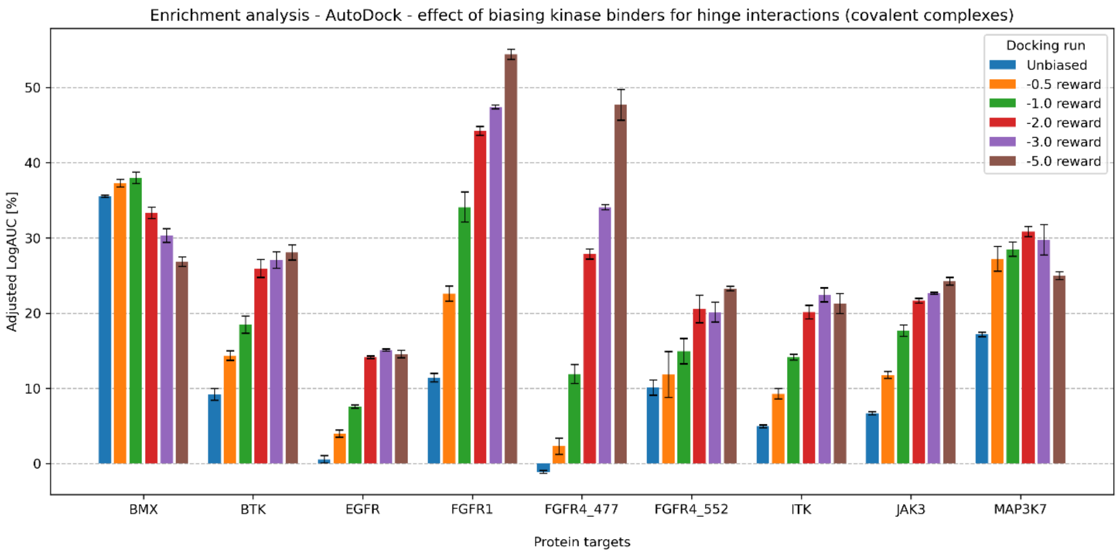


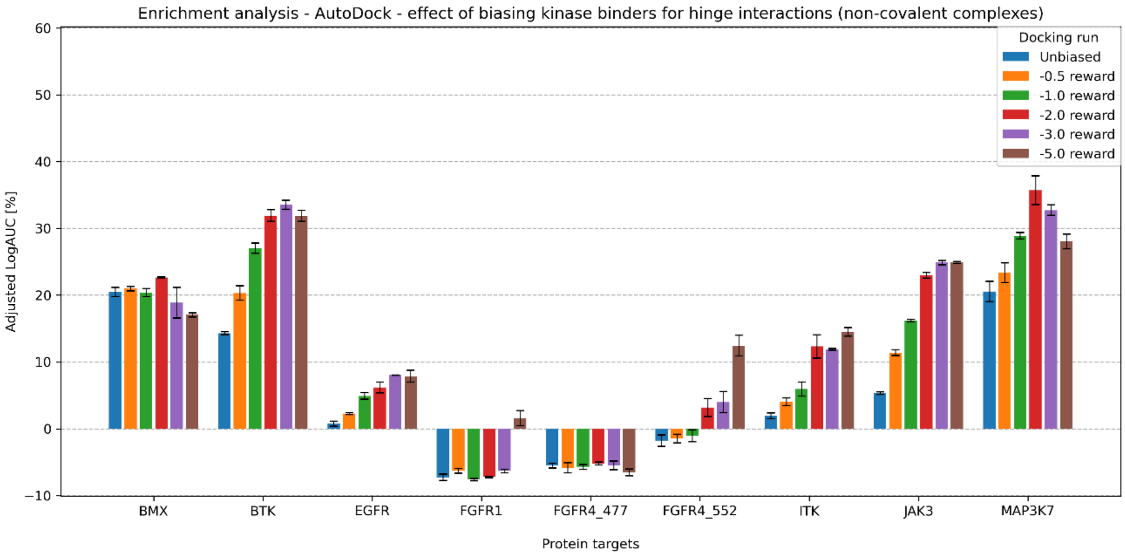


**Figure S34. AutoDock Bias effect on enrichment.** Adjusted LogAUC values from docking to the ten COValid Cys sites, reported as the average of three AutoDock replicates. An unbiased run is compared with runs that utilize AutoDock Bias for the formation of hydrogen bonds between the ligand and the two canonical hinge residues. Multiple different levels of energy rewards were used, ranging from -0.5 kcal/mol (least biased) to -5.0 kcal/mol (most biased). The decay radius was set to 1Å for all runs. The average adjusted LogAUC values, starting from unbiased to increasing bias, are 7.0±7.6%, 11.3±8.8%, 15.6±10.2% ,20.7±11.1%, 21.8±11.4%, and 22.7±11.2%, respectively. Fixed parameters: 10 GA runs, no bond angle diversification, maximal number of energy evaluations=250K.

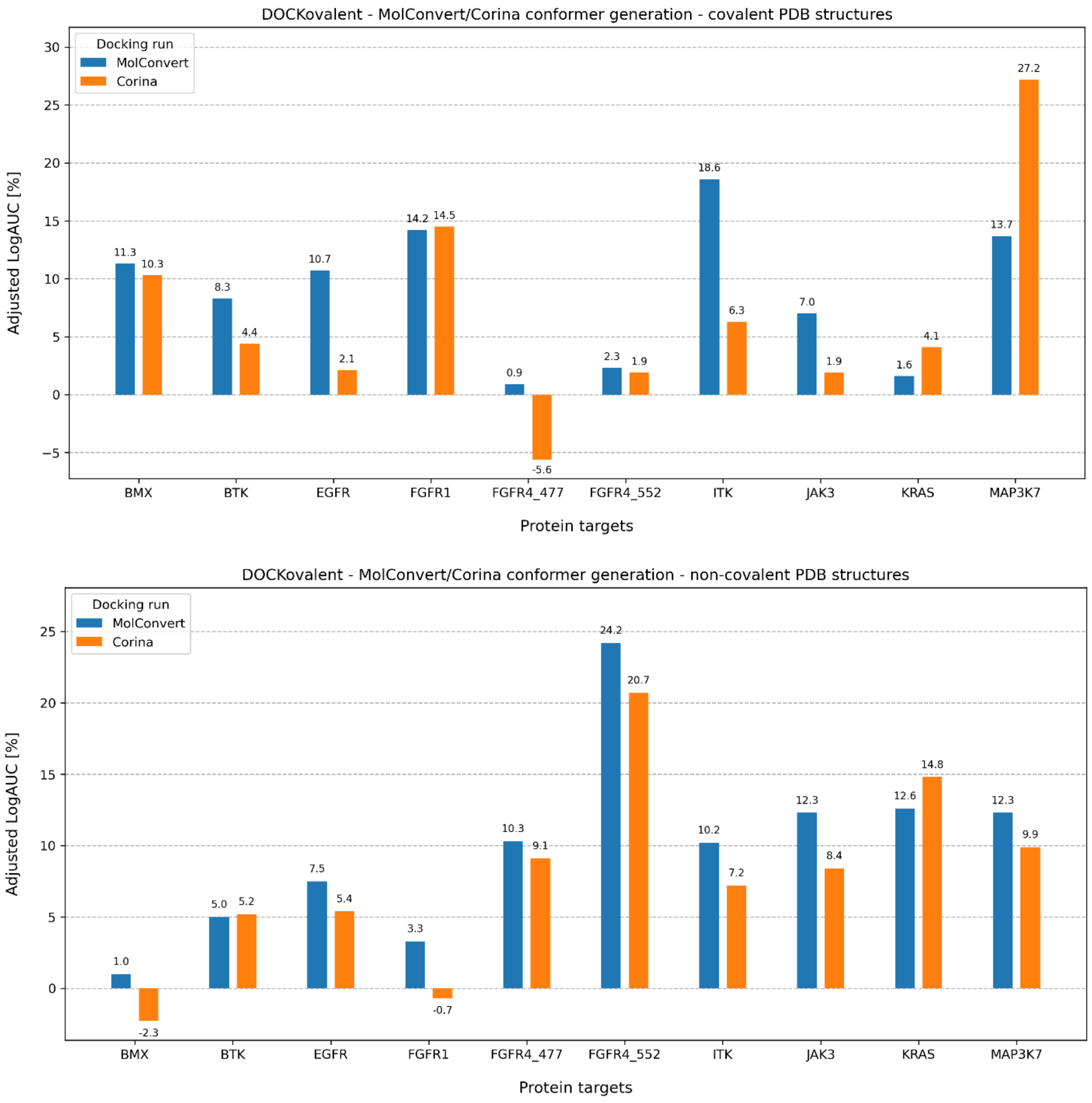


**Figure S35. DOCKovalent - effect of conformer generation software on enrichment.** We compared two tools for the generation of the initial 3D conformer for each protomer in the DOCKovalent ligand generation pipeline. The weighted average adjusted LogAUC values for MolConvert and Corina were 9.0±4.9% and 5.9±5.2%, respectively (p-value: 0.0014). Fixed parameters: maximal number of Omega conformers = 200.

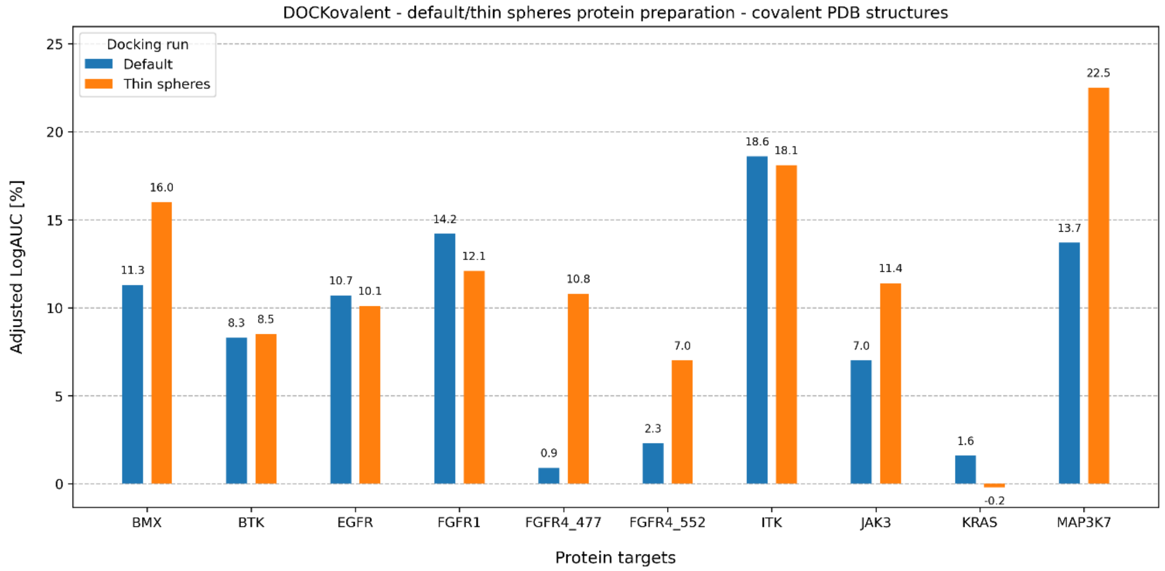


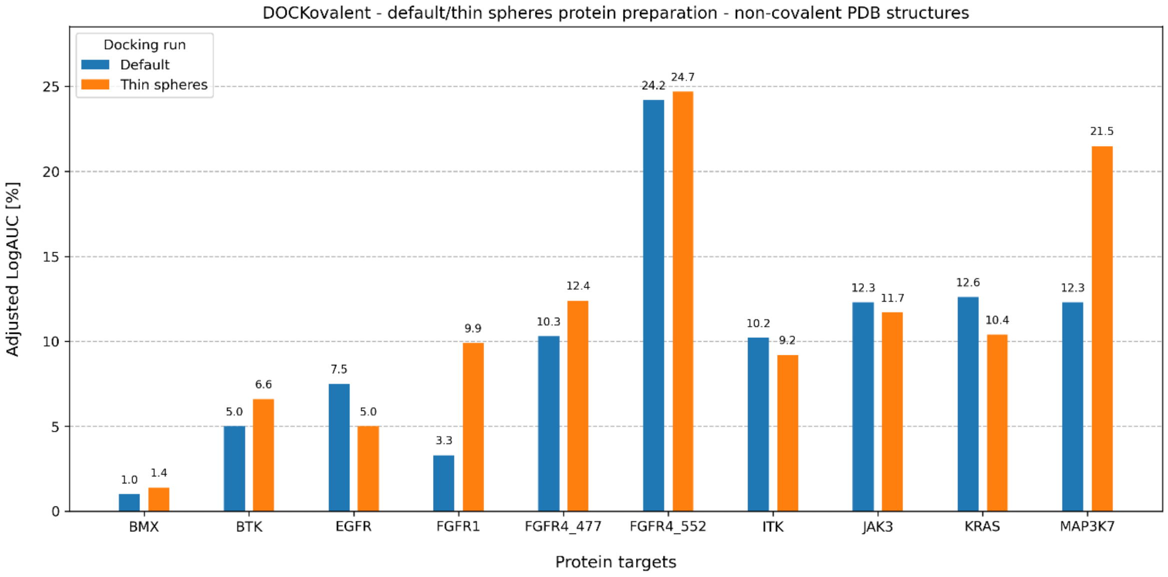


**Figure S36. DOCKovalent - effect of thin sphere protein preparation on enrichment.** We compared two protocols for the protein preparation step of DOCKovalent - the default BlasterMaster protocol, and the thin spheres protocol. The thin sphere protocol modifies the protein-solvent dielectric interface during the calculation of the electrostatics and desolvation grids (*16*). The weighted average adjusted LogAUC values for the default and thin sphere runs were 9.0±4.9% and 9.9±4.8%, respectively (*p-value*: 0.17). Fixed parameters: MolConvert for 3D conformer generation; maximal number of Omega conformers = 200.

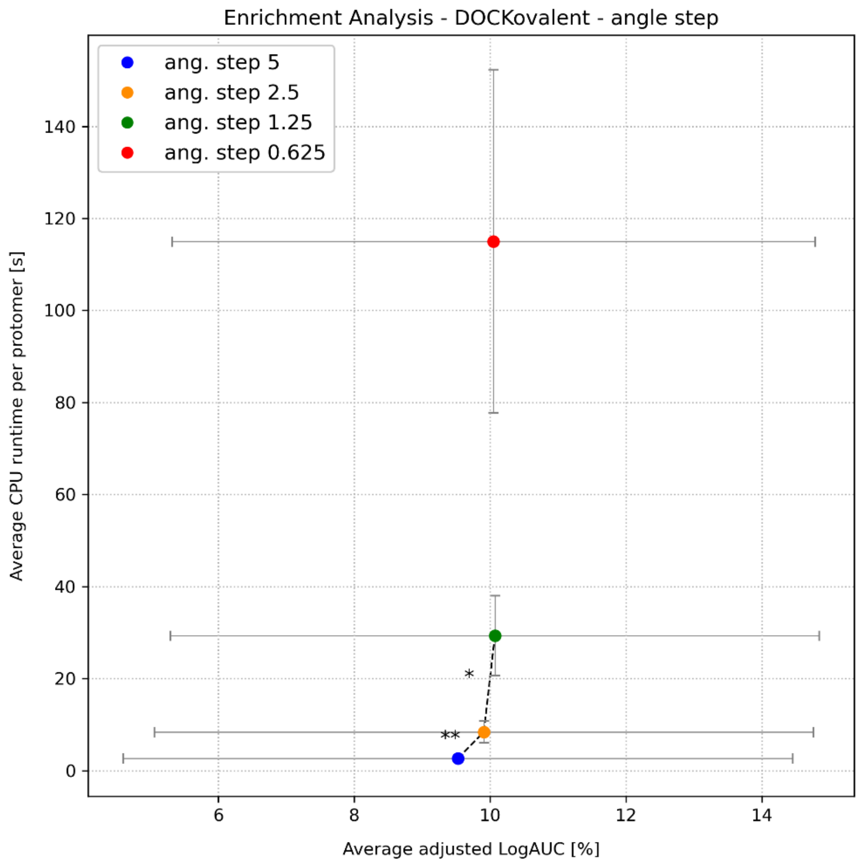


**Figure S37. DOCKovalent - effect of angle step parameter on enrichment.** This parameter determines the step size during sampling of the two bond angles at the covalent attachment site. Increasing the exhaustiveness from a step size of 5° to 2.5° and further to 1.25° leads to higher average adjusted LogAUC values with statistical significance (*p-values*: 0.0056 and 0.0172, respectively), while requiring significantly longer runtime. Fixed parameters: MolConvert for 3D conformer generation; maximal number of Omega conformers = 200; protein preparation with thin spheres; bond angle range = 10°.

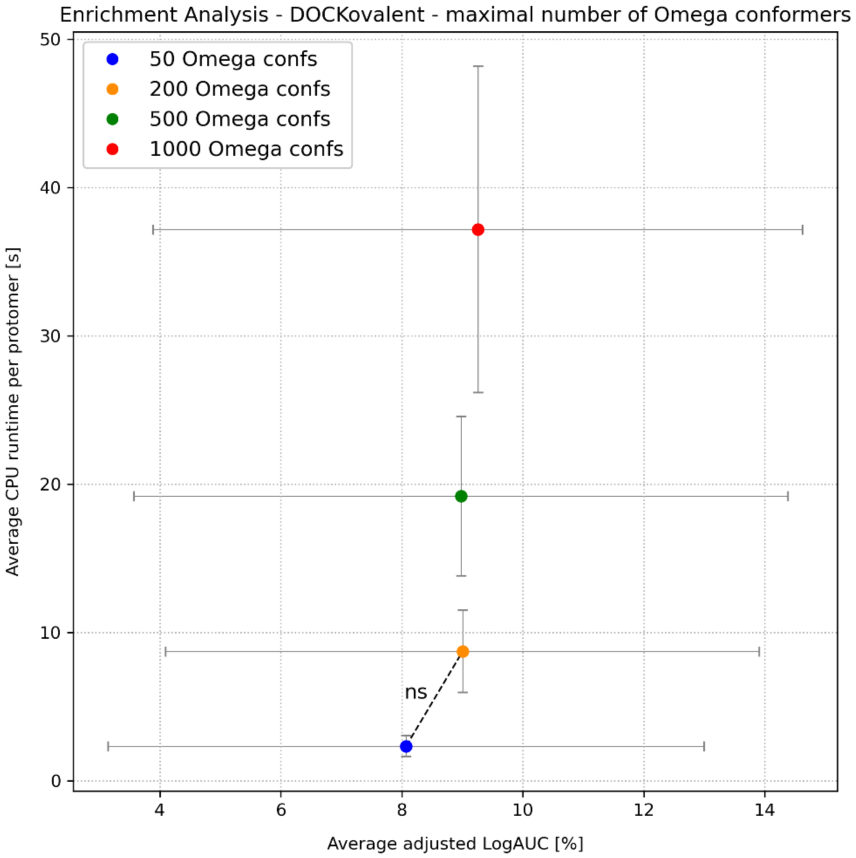


**Figure S38. DOCKovalent - effect of maximal number of Omega conformers on enrichment.** DOCKovalent uses Omega to expand the initial 3D conformer. We compared values for the maximal number of Omega conformers expanded per protomer. The increase in the average adjusted LogAUC when increasing the value from 50 to 200 is statistically insignificant (*p-value*: 0.1). Fixed parameters: MolConvert for 3D conformer generation.

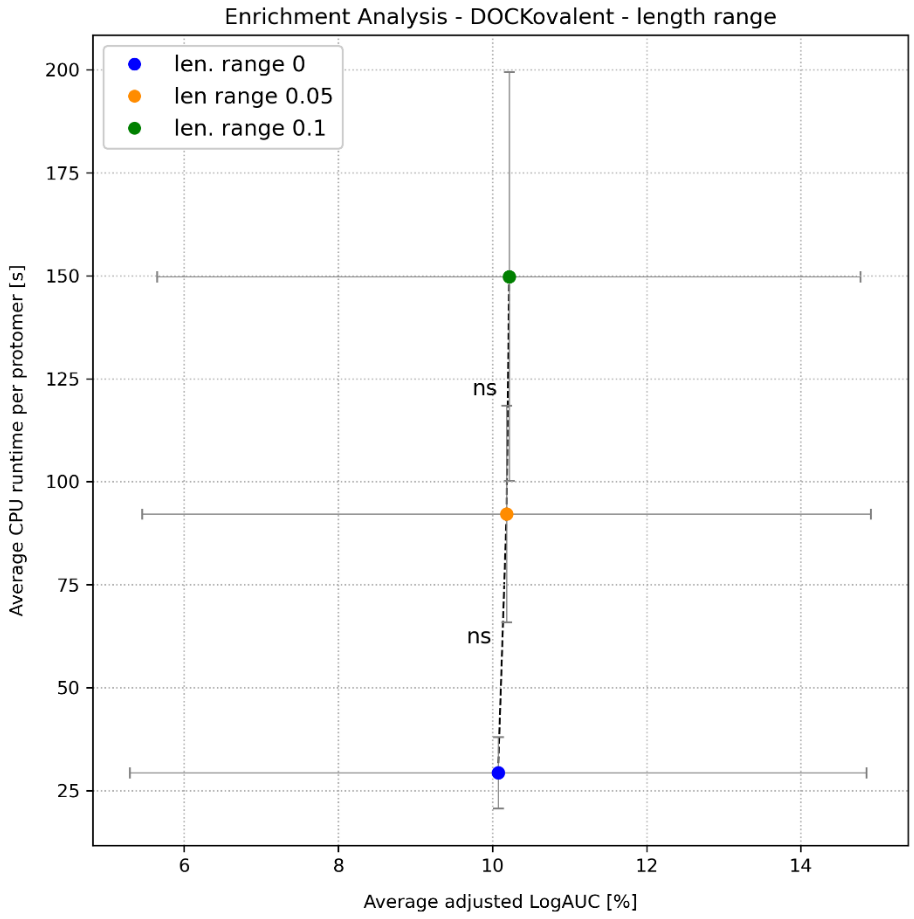


**Figure S39. DOCKovalent - effect of length range parameter on enrichment.** This parameter defines a range Δ around a user-defined bond length (1.8Å was used as the ideal), such that the cysteine Sγ-acrylamide Cβ bond is sampled ±Δ. Sampling with ±0.05Å or further with ±0.1Å yielded no significant increase to the average adjusted LogAUC value (*p-values*: 0.2 and 0.7, respectively). Fixed parameters: MolConvert for 3D conformer generation; maximal number of Omega conformers = 200; protein preparation with thin spheres; bond angle range = 10°; angle step = 1.25°.

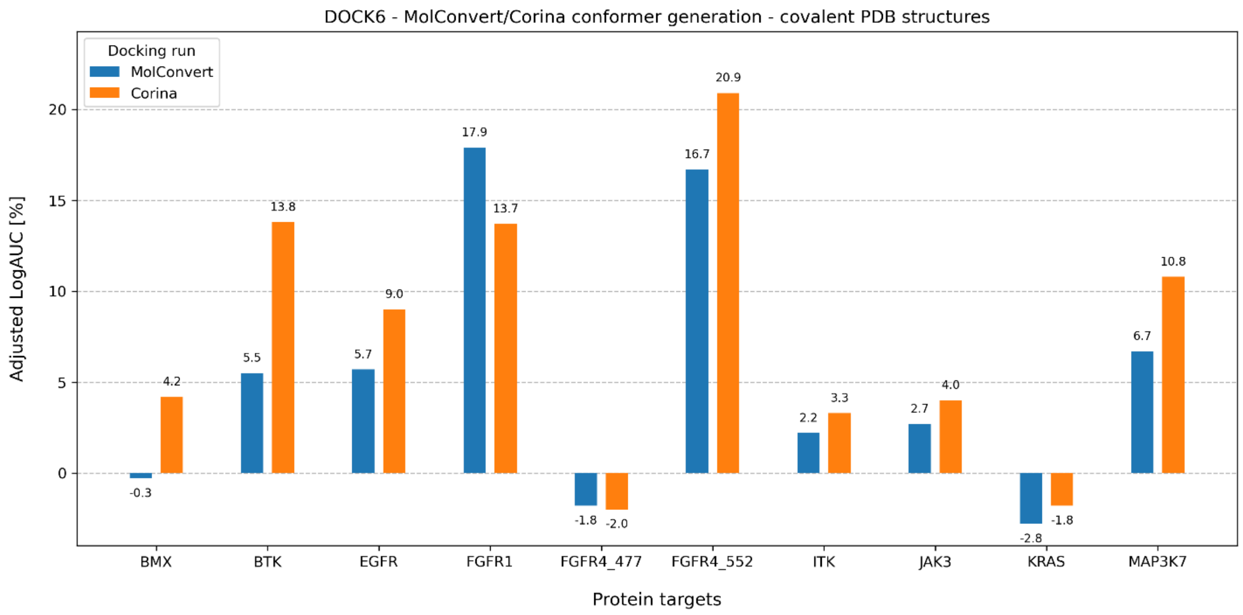


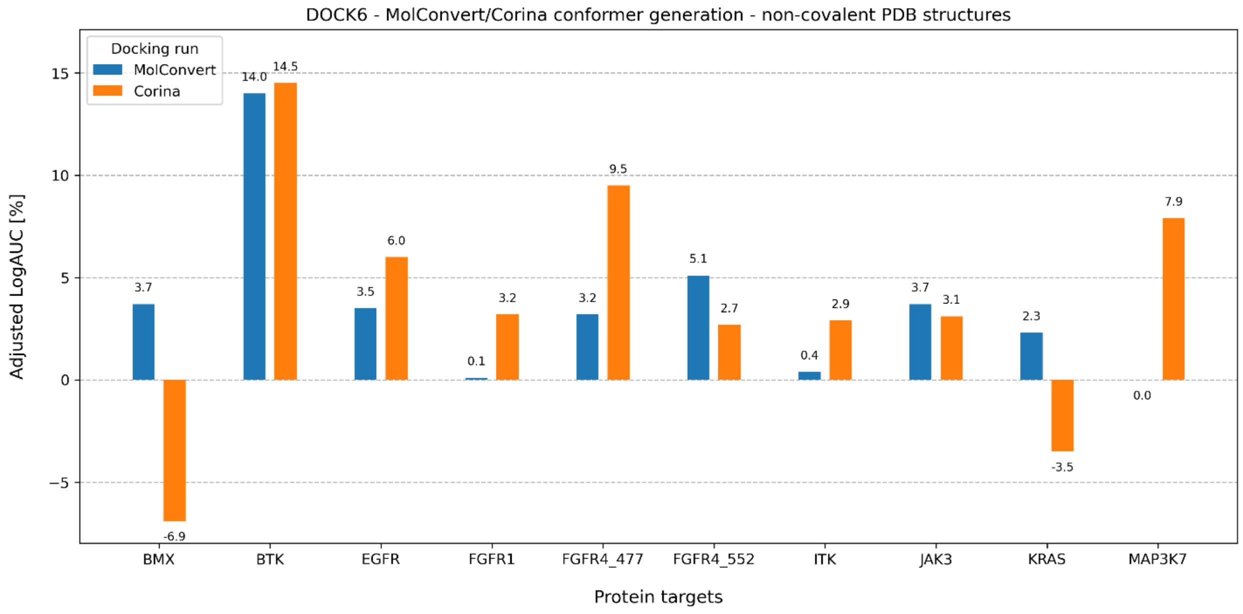


**Figure S40. DOCK6 - effect of conformer generation software on enrichment.** We compared two tools for the generation of the initial 3D conformer for each protomer in the DOCK6 ligand generation pipeline. The weighted average adjusted LogAUC values for MolConvert and Corina were 5.1±5.2% and 6.7±6.3%, respectively (*p-value*: 0.086). Fixed parameters: maximal number of Omega conformers = 200.

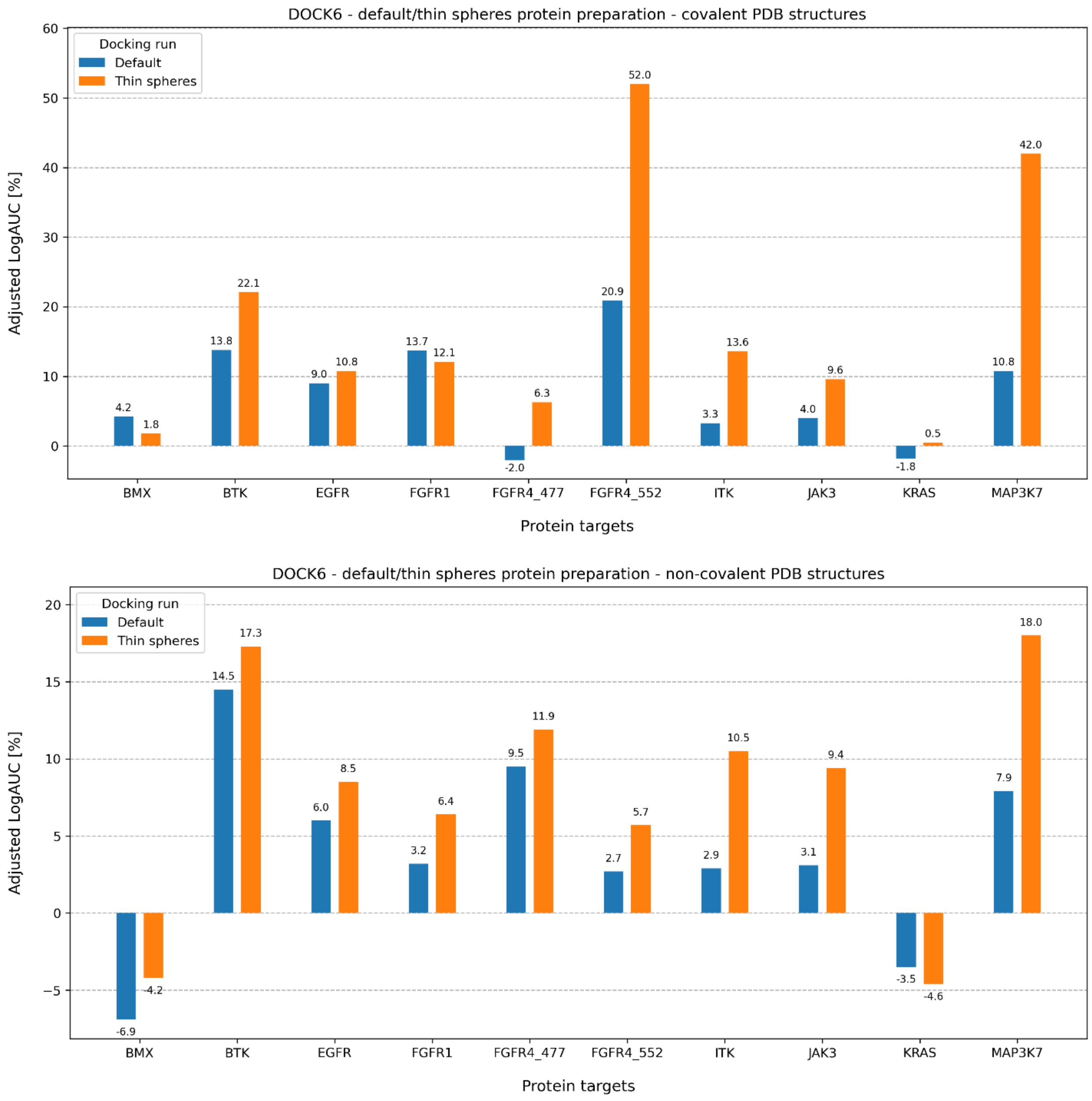


**Figure S41. DOCK6 - effect of thin sphere protein preparation on enrichment.** We compared two protocols for the protein preparation step of DOCK6 - the default BlasterMaster protocol, and the thin spheres protocol. The thin sphere protocol modifies the protein-solvent dielectric interface during the calculation of the electrostatics and desolvation grids (*16*). The weighted average adjusted LogAUC values for the default and thin sphere runs were 6.7±6.3% and 11.9±10.0%, respectively (*p-value*: 0.0006). Fixed parameters: Corina for 3D conformer generation; maximal number of Omega conformers = 200.

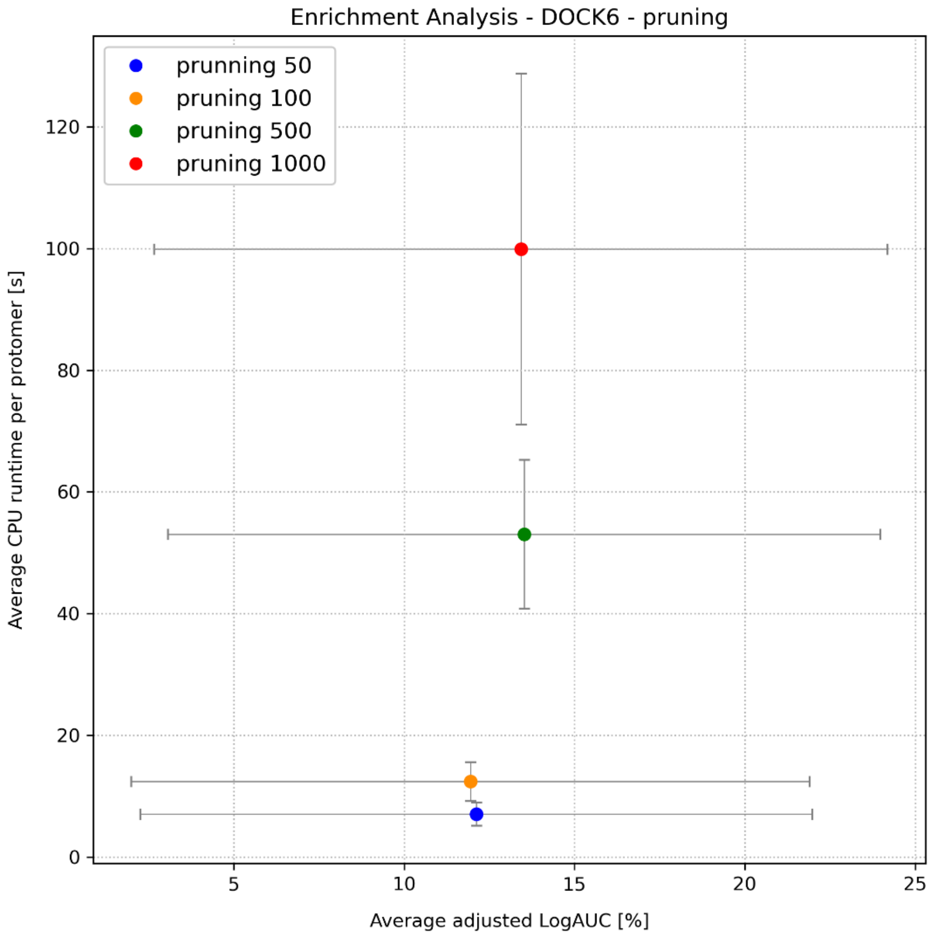


**Figure S42. DOCK6 - effect of pruning parameter on enrichment.** DOCK6.12 uses the attach-and-grow method for covalent docking. In this setting, torsions are sampled iteratively, with each conformer being evaluated with multiple dihedral positions. The torsionally sampled conformers are then pruned by a score cutoff as well as the pruning_clustering_cutoff parameter, which defines maximal number of conformers retained from the pruning process, according to a clustering process (*47*). Increasing the value of this parameter beyond 50 did not yield an increase to the average adjusted LogAUC. Fixed parameters: Corina for 3D conformer generation; maximal number of Omega conformers = 200; protein preparation with thin spheres.

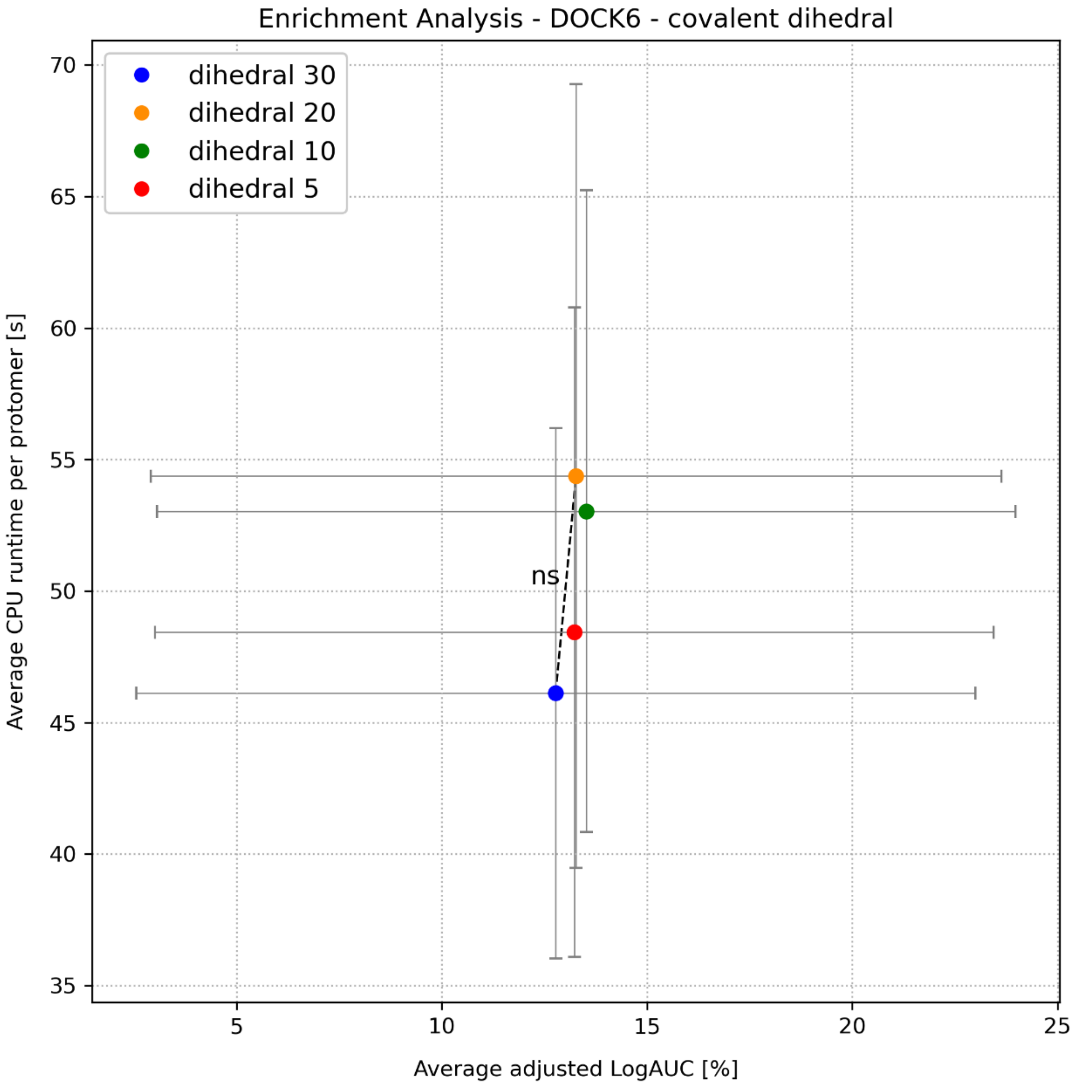


**Figure S43. DOCK6 - effect of covalent dihedral parameter on enrichment.** In the attach-and-grow protocol fo DOCK6.12, on top of the internal degrees of freedom of the ligand itself, the algorithm samples a few parameters at the attachment to the protein, including a torsion defined by Cys Cα, Cys Cβ, acrylamide Cβ, and acrylamide Cα atoms. The covalent_dihedral_step controls the step size of this sampling process. Decreasing the step size from 30° to 20° yielded an insignificant increase to the average adjusted LogAUC (p-value: 0.2). Fixed parameters: Corina for 3D conformer generation; maximal number of Omega conformers = 200; protein preparation with thin spheres; pruning parameter = 500.

### Supplementary Tables

| |
|---|
| **AutoDock** |
| 10 genetic algorithm iterations |
| Ligand generation – no bond angle diversification |
| Maximal number of energy evaluations = 50 |
| No use of Autodock Bias |
| **DOCKovalent** |
| MolConvert for initial conformer generation |
| Protein preparation with thin spheres |
| Maximal number of Omega conformers = 200 |
| Angle range = 10 |
| Angle step = 2.5 |
| no sampling of bond length range |
| **DOCK6** |
| Corina for conformer generation |
| Protein preparation with thin spheres |
| Pruning parameter = 50 |
| Covalent dihedral sampling parameter = 10 |

**Table S1. Docking parameters of leading covalent docking run configurations.** The results of runs using the above parameters were selected for comparison with AF3 predictions.

| Protein name | AUC-ROC | Adjusted LogAUC [%] |
|---|---|---|
| **BMX** | 0.9742 | 56.4 |
| **FGFR1** | 0.9962 | 79.4 |
| **FGFR4 C477** | 0.9996 | 83.8 |
| **FGFR4 C552** | 0.9988 | 82.5 |
| **JAK3** | 0.9696 | 71.6 |
| **EGFR** | 0.975 | 68.1 |
| **MAP3K7** | 0.9935 | 78.4 |
| **KRAS G12C** | 0.9967 | 74.7 |
| **BTK** | 0.9581 | 72.3 |
| **ITK** | 0.9082 | 65.8 |

**Table S2. AUC and LogAUC for AF3 predictions ranked by mPAE (worst-case scoring).** Reported is the area under the curve of the receiver operating curve, prior to log transformation of the x axis.

| protein target | residue name | residue index |
|---|---|---|
| **BTK** | GLU | 475 |
| **BTK** | MET | 477 |
| **JAK3** | LEU | 905 |
| **JAK3** | GLU | 903 |
| **ITK** | GLU | 436 |
| **ITK** | MET | 438 |
| **BMX** | GLU | 490 |
| **BMX** | ILE | 492 |
| **EGFR** | GLN | 791 |
| **EGFR** | MET | 793 |
| **FGFR1** | GLU | 562 |
| **FGFR1** | ALA | 564 |
| **MAP3K7** | ALA | 107 |
| **MAP3K7** | GLU | 105 |
| **FGFR4 C477** | ALA | 553 |
| **FGFR4 C477** | GLU | 551 |
| **FGFR4 C552** | ALA | 553 |
| **FGFR4 C552** | GLU | 551 |

**Table S3. Hinge interaction annotations for COValid kinases.** Listed are two canonical hinge residues of each kinase, known to form hydrogen bonds with ligands in the ATP binding pocket.

**Table S4. crystallographic data For YS1-BTK and YS2-BTK complexes.** § Statistics for the highest resolution shell are shown in parentheses.

| | **BTK in complex with YS1** | **BTK in complex with YS2** |
|---|---|---|
| **PDB code** | 9ZLJ | 9ZLM |
| **Data collection** | | |
| Resolution range§ | 59.21 - 1.496 (1.54 - 1.5) | 59.17 - 1.107 (1.12 - 1.11) |
| Space group | P 21 21 2 | P 21 21 2 |
| Cell dimensions | | |
| a, b, c (Å) | 72.024 103.974 38.069 | 71.97 103.943 37.973 |
| α, β, γ (°) | 90 90 90 | 90 90 90 |
| Multiplicity§ | 13.5 (13.0) | 8.9 (5.2) |
| Completeness (%)§ | 80.22 (3.38) | 67.43 (0.70) |
| Mean I/sigma (I) § | 5.25 (0.22) | 4.12 (0.26) |
| R-merge§ | 0.2576 (2.322) | 0.2064 (1.434) |
| CC1/2§ | 0.995 (0.504) | 0.99 (0.421) |
| **Refinement** | | |
| Resolution range§ | 38.07 – 1.60 (1.64 – 1.60) | 37.97 - 1.27 (1.29 - 1.27) |
| Unique reflections§ | 36621 (1600) | 71601 (1573) |
| R-work§ | 0.1846 (0.2484) | 0.1983 (0.2679) |
| R-free§ | 0.2177 (0.3177) | 0.2125 (0.2582) |
| Non-hydrogen atoms (n) | 2694 | 2603 |
| Protein | 2283 | 2271 |
| Ligand | 40 | 39 |
| Water | 371 | 293 |
| Protein residues | 268 | 268 |

| | BTK in complex with YS1 | BTK in complex with YS2 |
|---|---|---|
| Average B-factors (Å2) | 21.47 | 15.65 |
| Protein | 19.82 | 14.59 |
| Ligand | 18.59 | 14.84 |
| Water | 31.89 | 23.97 |
| RMS bonds (Å) | 0.002 | 0.095 |
| RMS angles (°) | 0.66 | 2.41 |
| Ramachandran favored (%) | 98.5 | 98.13 |
| Ramachandran allowed (%) | 1.5 | 1.87 |
| Ramachandran outliers (%) | 0.00 | 0.00 |

### Supplementary References

1. B. Zdrazil, E. Felix, F. Hunter, E. J. Manners, J. Blackshaw, S. Corbett, M. de Veij, H. Ioannidis, D. M. Lopez, J. F. Mosquera, M. P. Magarinos, N. Bosc, R. Arcila, T. Kizilören, A. Gaulton, A. P. Bento, M. F. Adasme, P. Monecke, G. A. Landrum, A. R. Leach, The ChEMBL Database in 2023: a drug discovery platform spanning multiple bioactivity data types and time periods. *Nucleic Acids Res.* **52**, D1180–D1192 (2024).

2. M. K. Gilson, T. Liu, M. Baitaluk, G. Nicola, L. Hwang, J. Chong, BindingDB in 2015: A public database for medicinal chemistry, computational chemistry and systems pharmacology. *Nucleic Acids Res.* **44**, D1045–53 (2016).

3. M. M. Mysinger, M. Carchia, J. J. Irwin, B. K. Shoichet, Directory of useful decoys, enhanced (DUD-E): better ligands and decoys for better benchmarking. *J. Med. Chem.* **55**, 6582–6594 (2012).

4. J. M. Bradshaw, J. M. McFarland, V. O. Paavilainen, A. Bisconte, D. Tam, V. T. Phan, S. Romanov, D. Finkle, J. Shu, V. Patel, T. Ton, X. Li, D. G. Loughhead, P. A. Nunn, D. E. Karr, M. E. Gerritsen, J. O. Funk, T. D. Owens, E. Verner, K. A. Brameld, R. J. Hill, D. M. Goldstein, J. Taunton, Prolonged and tunable residence time using reversible covalent kinase inhibitors. *Nat. Chem. Biol.* **11**, 525–531 (2015).

5. I. M. Serafimova, M. A. Pufall, S. Krishnan, K. Duda, M. S. Cohen, R. L. Maglathlin, J. M. McFarland, R. M. Miller, M. Frödin, J. Taunton, Reversible targeting of noncatalytic cysteines with chemically tuned electrophiles. *Nat. Chem. Biol.* **8**, 471–476 (2012).

6. R. T. Dungo, G. M. Keating, Afatinib: first global approval. *Drugs* **73**, 1503–1515 (2013).

7. M. Shirley, Dacomitinib: First global approval. *Drugs* **78**, 1947–1953 (2018).

8. P. J. Ropp, J. C. Kaminsky, S. Yablonski, J. D. Durrant, Dimorphite-DL: an open-source program for enumerating the ionization states of drug-like small molecules. *J. Cheminform.* **11**, 14 (2019).

9. R. A. Fairhurst, T. Knoepfel, C. Leblanc, N. Buschmann, C. Gaul, J. Blank, I. Galuba, J. Trappe, C. Zou, J. Voshol, C. Genick, P. Brunet-Lefeuvre, F. Bitsch, D. Graus-Porta, P. Furet, Approaches to selective fibroblast growth factor receptor 4 inhibition through targeting the ATP-pocket middle-hinge region. *Medchemcomm* **8**, 1604–1613 (2017).

10. Y. Wang, Y. Dai, X. Wu, F. Li, B. Liu, C. Li, Q. Liu, Y. Zhou, B. Wang, M. Zhu, R. Cui, X. Tan, Z. Xiong, J. Liu, M. Tan, Y. Xu, M. Geng, H. Jiang, H. Liu, J. Ai, M. Zheng, Discovery and development of a series of pyrazolo[3,4-d]pyridazinone compounds as the novel covalent fibroblast growth factor receptor inhibitors by the rational drug design. *J. Med. Chem.* **62**, 7473–7488 (2019).

11. H. Liu, D. Niu, R. T. Tham Sjin, A. Dubrovskiy, Z. Zhu, J. J. McDonald, K. Fahnoe, Z. Wang, M. Munson, A. Scholte, M. Barrague, M. Fitzgerald, J. Liu, M. Kothe, F. Sun, J. Murtie, J. Ge, J. Rocnik, D. Harvey, B. Ospina, K. Perron, G. Zheng, E. Shehu, L. A. D'Agostino, Discovery of selective, covalent FGFR4 inhibitors with antitumor activity in models of hepatocellular carcinoma. *ACS Med. Chem. Lett.* **11**, 1899–1904 (2020).

12. J. J. Irwin, K. G. Tang, J. Young, C. Dandarchuluun, B. R. Wong, M. Khurelbaatar, Y. S. Moroz, J. Mayfield, R. A. Sayle, ZINC20-A free ultralarge-scale chemical database for ligand discovery. *J. Chem. Inf. Model.* **60**, 6065–6073 (2020).

13. N. M. O'Boyle, M. Banck, C. A. James, C. Morley, T. Vandermeersch, G. R. Hutchison, Open Babel: An open chemical toolbox. *J. Cheminform.* **3**, 33 (2011).

14. R. M. Stein, Y. Yang, T. E. Balius, M. J. O'Meara, J. Lyu, J. Young, K. Tang, B. K. Shoichet, J. J. Irwin, Property-unmatched decoys in docking benchmarks. *J. Chem. Inf. Model.* **61**, 699–714 (2021).

15. D. Rogers, M. Hahn, Extended-connectivity fingerprints. *J. Chem. Inf. Model.* **50**, 742–754 (2010).

16. B. J. Bender, S. Gahbauer, A. Luttens, J. Lyu, C. M. Webb, R. M. Stein, E. A. Fink, T. E. Balius, J. Carlsson, J. J. Irwin, B. K. Shoichet, A practical guide to large-scale docking. *Nat. Protoc.* **16**, 4799–4832 (2021).

17. I. S. Knight, S. Naprienko, J. J. Irwin, Enrichment Score: a better quantitative metric for evaluating the enrichment capacity of molecular docking models, *arXiv [q-bio.QM]* (2022). http://arxiv.org/abs/2210.10905.

18. M. M. Mysinger, B. K. Shoichet, Rapid context-dependent ligand desolvation in molecular docking. *J. Chem. Inf. Model.* **50**, 1561–1573 (2010).

19. S. Wang, J. Witek, G. A. Landrum, S. Riniker, Improving conformer generation for small rings and macrocycles based on distance geometry and experimental torsional-angle preferences. *J. Chem. Inf. Model.* **60**, 2044–2058 (2020).

20. P. Tosco, N. Stiefl, G. Landrum, Bringing the MMFF force field to the RDKit: implementation and validation. *J. Cheminform.* **6**, 1–4 (2014).

21. J. Gasteiger, M. Marsili, Iterative partial equalization of orbital electronegativity—a rapid access to atomic charges. *Tetrahedron* **36**, 3219–3228 (1980).

22. J. Gasteiger, C. Rudolph, J. Sadowski, Automatic generation of 3D-atomic coordinates for organic molecules. *Tetrahedron Comput. Methodol.* **3**, 537–547 (1990).

23. P. C. D. Hawkins, A. G. Skillman, G. L. Warren, B. A. Ellingson, M. T. Stahl, Conformer generation with OMEGA: algorithm and validation using high quality structures from the Protein Databank and Cambridge Structural Database. *J. Chem. Inf. Model.* **50**, 572–584 (2010).

24. G. Bianco, S. Forli, D. S. Goodsell, A. J. Olson, Covalent docking using autodock: Two-point attractor and flexible side chain methods: Covalent Docking with AutoDock. *Protein Sci.* **25**, 295–301 (2016).

25. V. Sharma, M. Gupta, Designing of kinase hinge binders: A medicinal chemistry perspective. *Chem. Biol. Drug Des.* **100**, 968–980 (2022).

26. L. Xing, J. Klug-Mcleod, B. Rai, E. A. Lunney, Kinase hinge binding scaffolds and their hydrogen bond patterns. *Bioorg. Med. Chem.* **23**, 6520–6527 (2015).

27. J. P. Arcon, C. P. Modenutti, D. Avendaño, E. D. Lopez, L. A. Defelipe, F. A. Ambrosio, A. G. Turjanski, S. Forli, M. A. Marti, AutoDock Bias: improving binding mode prediction and virtual screening using known protein-ligand interactions. *Bioinformatics* **35**, 3836–3838 (2019).

28. J. P. Arcon, A. G. Turjanski, M. A. Martí, S. Forli, Biased docking for protein-ligand pose prediction. *Methods Mol. Biol.* **2266**, 39–72 (2021).

29. N. London, R. M. Miller, S. Krishnan, K. Uchida, J. J. Irwin, O. Eidam, L. Gibold, P. Cimermančič, R. Bonnet, B. K. Shoichet, J. Taunton, Covalent docking of large libraries for the discovery of chemical probes. *Nat. Chem. Biol.* **10**, 1066–1072 (2014).

30. H. Park, P. Bradley, P. Greisen Jr, Y. Liu, V. K. Mulligan, D. E. Kim, D. Baker, F. DiMaio, Simultaneous optimization of biomolecular energy functions on features from small molecules and macromolecules. *J. Chem. Theory Comput.* **12**, 6201–6212 (2016).

31. Y. Zhang, J. Skolnick, Scoring function for automated assessment of protein structure template quality. *Proteins* **57**, 702–710 (2004).

32. R. E. Joseph, I. Kleino, T. E. Wales, Q. Xie, D. B. Fulton, J. R. Engen, L. J. Berg, A. H. Andreotti, Activation loop dynamics determine the different catalytic efficiencies of B cell- and T cell-specific tec kinases. *Sci. Signal.* **6**, ra76 (2013).

33. D. Y. Lin, A. H. Andreotti, Structure of BTK kinase domain with the second-generation inhibitors acalabrutinib and tirabrutinib. *PLoS One* **18**, e0290872 (2023).

34. P. R. Evans, G. N. Murshudov, How good are my data and what is the resolution? *Acta Crystallogr. D Biol. Crystallogr.* **69**, 1204–1214 (2013).

35. W. Kabsch, XDS. *Acta Crystallogr. D Biol. Crystallogr.* **66**, 125–132 (2010).

36. C. Vonrhein, C. Flensburg, P. Keller, A. Sharff, O. Smart, W. Paciorek, T. Womack, G. Bricogne, Data processing and analysis with the autoPROC toolbox. *Acta Crystallogr. D Biol. Crystallogr.* **67**, 293–302 (2011).

37. A. J. McCoy, R. W. Grosse-Kunstleve, P. D. Adams, M. D. Winn, L. C. Storoni, R. J. Read, Phaser crystallographic software. *J. Appl. Crystallogr.* **40**, 658–674 (2007).

38. Smart, O. S. , Sharff A. , Holstein, J. , Womack, T. O. , Flensburg, C. , Keller, P. , Paciorek, W. , Vonrhein, C. and Bricogne G. , Cambridge, United Kingdom: Global Phasing Ltd., *Grade2 Version 1.7.1* (2021).

39. N. W. Moriarty, R. W. Grosse-Kunstleve, P. D. Adams, electronic Ligand Builder and Optimization Workbench (eLBOW): a tool for ligand coordinate and restraint generation. *Acta Crystallogr. D Biol. Crystallogr.* **65**, 1074–1080 (2009).

40. D. Liebschner, P. V. Afonine, M. L. Baker, G. Bunkoczi, V. B. Chen, T. I. Croll, B. Hintze, L.-W. Hung, S. Jain, A. J. McCoy, N. W. Moriarty, R. D. Oeffner, B. K. Poon, M. G. Prisant, R. J. Read, J. S. Richardson, D. C. Richardson, Sammito, O. V. Sobolev, D. H. Stockwell, T. C. Terwilliger, A. G. Urzhumtsev, L. L. Videau, C. J. Williams, P. D. Adams, Macromolecular structure determination using X-rays, neutrons and electrons: recent developments in Phenix. International Union of Crystallography [Preprint].

https://doi.org/10.1107/S2059798319011471.

41. P. Emsley, K. Cowtan, Coot: model-building tools for molecular graphics. *Acta Crystallogr. D Biol. Crystallogr.* **60**, 2126–2132 (2004).

42. J. Sadowski, J. Gasteiger, G. Klebe, Comparison of automatic three-dimensional model builders using 639 X-ray structures. *J. Chem. Inf. Comput. Sci.* **34**, 1000–1008 (1994).

43. MolConvert, Chemaxon (https://www.chemaxon.com).

44. S. Passaro, G. Corso, J. Wohlwend, M. Reveiz, S. Thaler, V. R. Somnath, N. Getz, T. Portnoi, J. Roy, H. Stark, D. Kwabi-Addo, D. Beaini, T. Jaakkola, R. Barzilay, Boltz-2: Towards accurate and efficient binding affinity prediction, *bioRxivorg* (2025)p. 2025.06.14.659707.

45. The PyMOL Molecular Graphics System, Version 3.7.12 Schrödinger, LLC.

46. S. Eid, S. Turk, A. Volkamer, F. Rippmann, S. Fulle, KinMap: a web-based tool for interactive navigation through human kinome data. *BMC Bioinformatics* **18**, 16 (2017).

47. Y. S. Tan, M. Chakrabarti, R. M. Stein, L. E. Prentis, R. C. Rizzo, T. Kurtzman, M. Fischer, T. E. Balius, Development of receptor desolvation scoring and covalent sampling in DOCK 6: Methods evaluated on a RAS test set. *J. Chem. Inf. Model.*, doi: 10.1021/acs.jcim.4c01623 (2025).
